# From tiny to massive: exploring genome evolution in the model grass genus *Brachypodium*

**DOI:** 10.64898/2026.07.16.738966

**Authors:** Chunlin Chen, Rubén Sancho, Miguel Campos, Bruno Contreras-Moreira, Jeremy Schmutz, Jerry Jenkins, Kerrie Barry, Anna Lipzen, Maxim Koriabine, Joanne Eichenberger, Li Lei, John P. Vogel, David L. Des Marais, Pilar Catalán

## Abstract

*Brachypodium* is a powerful model system for investigating grass genome evolution, yet genomic resources remain concentrated in the three annual species, whereas perennial species are less sampled. Here, we present chromosome-level assemblies for two perennial species, *Brachypodium mexicanum* and *B. arbuscula*, which represent the earliest-diverging lineages of the genus and the earliest-diverging lineage of the core perennial clade, respectively. Synteny-based phylogenomics indicate that *B. mexicanum* is a meso-allotetraploid composed of two closely related but temporally distinct x=10 subgenomes, here designed as P and U, each carrying subgenome-specific chromosome rearrangements. We further show that the unusually large *B. mexicanum* genome, in contrast to the reduced genomes of most other *Brachypodium* species, is primarily due to transposable elements distributed across all chromosomal regions. By contrast, the diploid genome of the earliest-diverging core perennial, *B. arbuscula*, contains few transposable elements, whereas the most recent diverged diploid perennial *B. sylvaticum* shows evidence of a secondary TEs proliferation. Comparisons of lineage-specific and functionally enriched orthogroups among *B. mexicanum*, core perennial species and annual species suggest that ancestral hybridization between annual and perennial lineages may have contributed to the origin of allotetraploid *B. mexicanum*. These assemblies provide a framework for testing how polyploidy, descending dysploidy, transposable-element turnover, and life-history evolution jointly shaped genome architecture in *Brachypodium*.

## Introduction

Genome evolution in cool-season grasses of subfamily Pooideae has been shaped by recurrent interactions among polyploidy, post-polyploid diploidization, descending dysploidy, and transposable element turn-over. These processes are associated with major shifts in base chromosome numbers, genome size, gene presence/absence variation (PAV), copy-number variation (CNV), and life-history strategies. Earlier-diverging tribal lineages generally retain higher chromosome base numbers (x=12, 11, or 10; Nardeae, Lygeae, Stipeae), whereas intermediate lineages, including Brachypodieae, display x=10, 9, 8, or 5, and the more recently evolved core pooids often have a reduced basic chromosome number of x=7 or lower (Triticeae, Bromeae, or Poeae)^1,2,3^. Successive rounds of whole genome duplication (WGD) events and diploidization have further produced paleopolyploids, diploidized paleopolyploids, and more recent meso– and neopolyploids^4,5^. *Brachypodium* (tribe Brachypodieae) provides a critical comparative system for studying genome restructuring and life-history evolution during grass diversification^6,7^ because of its intermediate phylogenetic position between earlier-diverging and core pooids.

*Brachypodium* genus has therefore become a model system for functional genomics, allopolyploidization, and the evolutionary transition between annual and perennial growth forms in temperate cereals and forage grasses^8–11^. The genus also contains the largest karyotypic variation documented within the Pooideae, with at least seven karyotypes that differ in chromosome base number and chromosomal rearrangements history. These range from the more ancestral x=10 karyotypes of *B. mexicanum*-4x and *B. stacei*-2x, through intermediate x=8 and x = 5 karyotypes, to more recently evolved x= 9, x=8, and x=5 karyotypes in core perennial species^12,13^. A common feature of the core Pooideae is genome-size expansion, apparently driven largely by bursts of repetitive-elements following or accompanying WGDs^4,14,15^. By contrast, *Brachypodium* is generally characterized by the small monoploid genomes, including *B. distachyon* (272 Mbp; 0.36 pg/1C)^8,16^ and *B. stacei* (234 Mbp; 0.28 pg/1C)^9,17,18^, with the striking exception of its most ancestral polyploid species *B. mexicanum* (0.94 pg/1C)^18^ whose monoploid genome is more than three times larger than the smallest *Brachypodium* haploid genome. The massive size difference is associated with extensive repetitive-element accumulation^18^. However, previous comparative analysis based on transcriptomic or genome skimming data could not fully resolve the repetitive region of *B. mexicanum*^13,18^.

*Brachypodium mexicanum* presents unusual combination of perennial features and annual-like traits. It has slender growth habit, highly selfing capacity, and rDNA-marker affinities with annual *Brachypodium* species^6,19,20^. Its incompatibility with other perennial *Brachypodium* species^20^ and its phylogenetic position identify it as one of the earliest-diverging lineages of the genus^13,21^. *Brachypodium arbuscula* is likewise important for understanding perennial genome evolution because it shares the x=9 karyotype with diploid perennials *B. sylvaticum* and *B. pinnatum*^6,13^, while retaining branched stems, a relict trait shared with the ancestral *B. boissieri* and *B. retusum* species^6^. Phylogenetic analyses consistently resolve *B. arbuscula* as the earliest-diverging lineage within the recently evolved core perennial clade^13,21^.

*Brachypodium* lineages have undergone repeated hybridization and polyploidization throughout their evolutionary history^6,13,21^. Their genomes show complex chromosomal rearrangements and an overall pattern of descending dysploidy trend, with chromosome base numbers ranging from ten to five and with distinct karyotypic configurations among genomes and subgenomes^12,13^. Post-polyploid diploidizations and descending dysploidies have likely contributed to genome-size change among descendant species and cytotypes^13,18^. However, chromosome size variation also reflects the proliferation and loss of transposable elements, especially long terminal repeats (LTR) retrotransposons^18^. LTR retrotransposons are major drivers of genome evolution among closely related plant species^22,23^, and their accumulation can contribute more strongly than WGD to genome-size differences across plants^24,25^. Dramatic genome-size variation, extensive chromosome rearrangements, diverse life histories, and established annual and perennial model taxa^8–11^ make *Brachypodium* well suited for disentangling how gene-content variation and TE dynamics shape genome architecture.

Here, we assembled and annotated high-quality chromosome-level genomes for two perennial *Brachypodium* species, *B. mexicanum* and *B. arbuscula*, which represent the largest and smallest perennial genomes described for the genus, respectively^13,18^. Both k-mer– and phylogeny-based approaches reveal that *B. mexicanum* is a meso-allotetraploid composed of two divergent x=10 subgenomes: a more ancestral P subgenome and a more recently derived U subgenome. We further demonstrate that *B. arbuscula*, as a representative of the ancestral core perennials, retains a gene content similar to that of *B. sylvaticum* but contains fewer repeat elements. The *Brachypodium* pangenome reveals how karyotype structure, gene PAV, and TE dynamics differ between annual and perennial lineages and between diploid and polyploid genomes. Across *Brachypodium,* variation in LTR abundance explains genome-size differences more strongly than coding-sequence length, highlighting TE turnover as a major driver of genome architectural evolution in this model grass genus.

## Results

### Genome assembly and annotation of *B. arbuscula* and *B. mexicanum*

The haploid genome of *B. arbuscula* (https://phytozome-next.jgi.doe.gov/info/BarbusculaBARB1_v3_1) contains nine pseudochromosomes (Fig. 1a and Supplementary Fig. 1a) with a total size of 283.55 Mb and an N50 scaffold length of 30.19 Mb (Supplementary Table 1). A total of 33,605 protein-coding genes were annotated, encompassing 99.10% of the complete BUSCO genes (Supplementary Table 1). The haploid genome of *B. mexicanum* (https://phytozome-next.jgi.doe.gov/info/Bmexicanum_v2_1) contains 20 pseudochromosomes (Fig. 1b and Supplementary Fig. S1b) with a total size of 1,530.5 Mb and an N50 scaffold of 78.41 Mb (Supplementary Table 1). A total of 70,845 protein-coding genes were annotated for *B. mexicanum*, including 99.50% of the complete BUSCO genes (Supplementary Table 1). Both SubPhaser and PhyloSD approaches resolved the same subgenomic chromosome assignments for the two subgenomes of *B. mexicanum* (Supplementary Figs. 2a and 2b), the more ancestral one of which is here denoted as subgenome “P” (paleo), while the putatively younger one is denoted “U” (Supplementary Fig. 2b). Our synonymous substitution distance analysis identified shared *Ks* peaks between paralogous genes for the *B. arbuscula* genome and the two respective subgenomes of *B. mexicanum* at around *Ks* = 0.8, t = 97.63-104.40 million years ago (Ma), coinciding with the WGD (rho) of the proto-grass ancestor (Supplementary Fig. 3). In addition, a more recent and higher *Ks* peak between homeologous genes of the P and U subgenomes was identified at around *Ks* = 0.1, t = 13.58 Ma, corroborating its meso-polyploid origin.

**Fig. 1.**
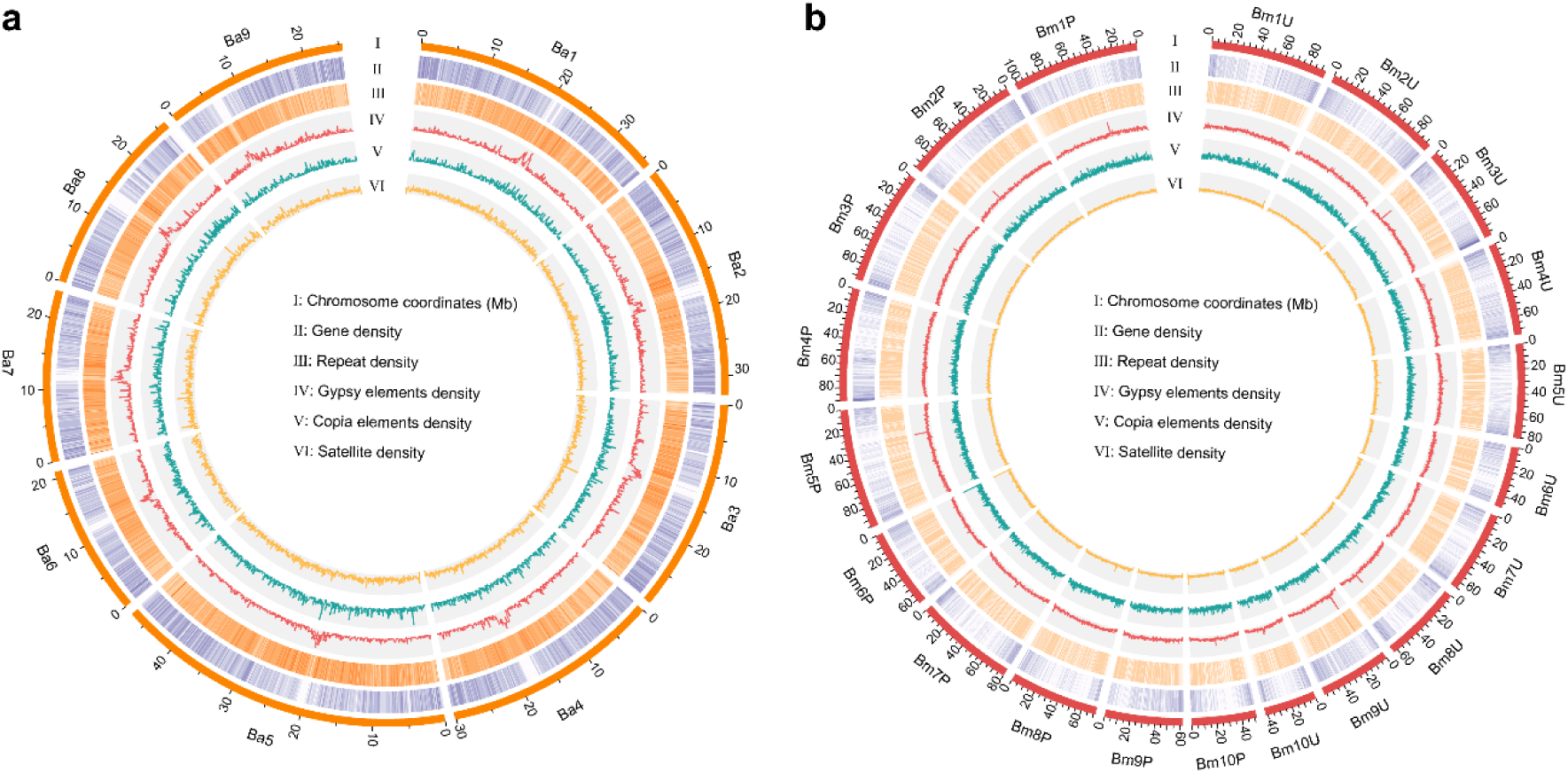
Circos plot of diploid x=9 *Brachypodium arbuscula*. (**a**) and allotetraploid x=10P+10U *B. mexicanum* **(b**) genomes. Information on layers ‘a’ to ‘f’ are shown inside the circle. From layers ‘b’ to ‘f’, the densities are shown in 200 kbp windows for *B. arbuscula* and 500 kbp windows for *B. mexicanum,* respectively.

**Fig. 2.**
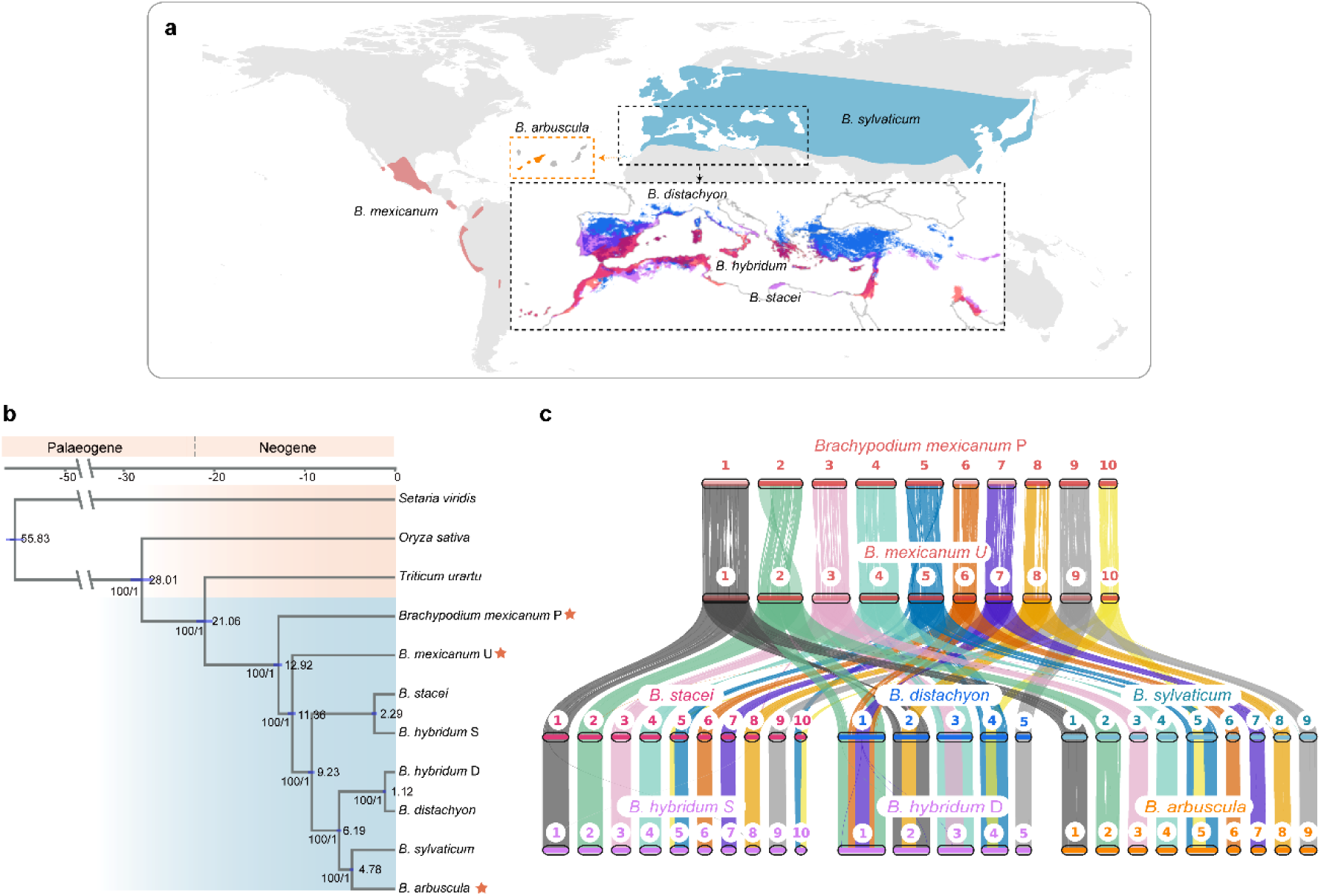
Phylogenomics and synteny of six *Brachypodium* species’ reference genomes, the newly assembled genomes of the perennials *B. arbuscula* (2x) and *B. mexicanum* (4x), plus those of annuals *B. distachyon* (2x), *B. stacei* (2x) and *B. hybridum* (4x) and the perennial *B. sylvaticum* (2x). (**a**) Geographic distribution of the six *Brachypodium* species studied here. (**b)** Dated genomic phylogeny of the eight *Brachypodium* (sub)genomes corresponding to the six *Brachypodium* species (allotetraploids *B. mexicanum* (P and U) and *B. hybridum* (S and D) contribute with two subgenomes each). Values on nodes indicate the estimated divergence times and blue bars show their 95% high credibility interval. Support values for all nodes, indicated below branches, reached 100 bootstrap percentages or posterior probability of 1 in all cases. Orange stars highlight the (sub)genomes generated in this study. (**c)** Syntenic relationship among the eight (sub)genomes of the six *Brachypodium* species under study, showing chromosomal rearrangements between the ancestral (sub)genomes P and U of *B. mexicanum* and S of *B. stacei* (and its derived *B. hybridum*-S subgenome) (x=10), and different nested chromosome fusions involved in the descendant dysploidies that originated the more recently evolved genomes D (x=5) in *B. distachyon* (and its derived *B. hybridum*-D subgenome) and G (x=9) in core-perennial taxa (*B. sylvaticum*, *B. arbuscula*).

### Comparative genomics and phylogenomics of perennial and annual *Brachypodium* species and genomes

All six *Brachypodium* genomes studied (Fig. 2a) had high BUSCO completeness (Supplementary Fig. 4). A total of 2,131 no-missing single-copy orthologous (ortho-homeologous) gene clusters across the (sub)genomes were used for phylogeny construction. For maximum likelihood concatenation-based method a subsequent species tree was estimated. Both concatenated and coalescent based phylogenies yielded consistent tree topologies with all nodes resolved with full support (i.e., bootstrap support percentage = 100, or posterior probability = 1; Fig. 2b). A total of 832 genes that showed a congruent topology with the species tree were kept for divergence time estimation. Phylogenomic analysis resolved the *B. mexicanum* P and U subgenomes as the earliest-diverging lineages (12.92 and 11.36 Ma), followed by *B. stacei* (9.23 Ma), *B. distachyon* (6.19 Ma), and the core-perennial clade (4.78 Ma). The two *B. hybridum* subgenomes diverged from their respective diploid progenitors at 2.29 and 1.12 Ma (Fig. 2b).

### Synteny among the *Brachypodium* genomes and subgenomes

Our synteny analysis identified largely consistent syntenic chromosomal blocks among all the investigated *Brachypodium* (sub)genomes (Fig. 2c). When interpreted within the evolutionary framework of our phylogenomic tree, we observed an overall descending dysploidy trend from the ancestral karyotype x=10 of *B. mexicanum*-P to that of *B. mexicanum*-U, and that of *B. stacei*, with each process experiencing different chromosomal rearrangements. We also inferred two independent reductions, a pronounced reduction from x = 10 to the x = 5 karyotype of the *B. distachyon* lineage, and a less drastic reduction to the x = 9 karyotype of core perennial clade species (Fig. 2c). Within the initial chromosomal rearrangements of the x = 10 (sub)genomes of *B. mexicanum*, the syntenic blocks identified two translocations that occurred between the chromosomes Bmex2P and Bmex2U, and Bmex5P and Bmex5U, and multiple inversions between homologous chromosomes pairs between the two subgenomes (Fig. 2c and Supplementary Fig. 5). In addition, a block consisting of around 400 genes was translocated from Bmex8P to Bmex3U (Fig. 2c, and Supplementary Fig. 6). A reciprocal translocation between chromosomes Bmex5U and Bmex10U of *B. mexicanum* gave rise to the x=10 karyotype of *B. stacei* (Fig. 2c). The strong descending dysploidy reduction from x=10 chromosomes to the x=5 chromosomes of *B. distachyon* likely occurred through five nested chromosome fusions; (i) chromosome Bd1 resulted from two nested fusions of chromosomes Bmex2U, Bmex6U and Bmex7U of *B. mexicanum*, (ii) chromosome Bd3 from chromosomes Bmex1U and Bmex8U, (iii) chromosome Bd3 from chromosomes Bmex3U and Bmex4U, and (iv) chromosome Bd4 from chromosomes Bmex5U and Bmex10U, while chromosome Bd5 of *B. distachyon* is fully collinear to chromosome Bmex9U (Fig. 2c). On the other hand, the descendant dysploidy trajectory that reduced x=10 to x = 9 chromosomes present in the core perennial species implied one nested fusion of chromosomes Bmex5U and Bmex10U of *B. mexicanum* to form the chromosome five of *B. arbuscula* (Ba5) and *B. sylvaticum* (Fig. 2c). The remaining eight chromosomes share a high synteny with the respective chromosomes of subgenome U of *B. mexicanum*. The two subgenomes of *B. hybridum* maintained high synteny with their respective progenitor species genomes (*B. stacei* and *B. distachyon*) (Fig. 2c).

### Presence/absence analysis of CDS orthogroups specific to annual and perennial species

A total of 106,180 orthogroups were detected in the CDS analysis of the six *Brachypodium* species. Of these, 346 contained sequences exclusive to the perennial species (*B. arbuscula, B. mexicanum, B. sylvaticum*), 105 specific to the annual species (*B. distachyon, B. stacei, B. hybridum*), and 135 from the annual species plus *B. mexicanum* (Supplementary Table 2). Because *B. mexicanum* shows some morphological and physiological intermediate features between the annual and perennial species^6^, we further examined the distribution of the gene sequences exclusive to each life history type within the P and U subgenomes. Of the 346 orthogroups exclusive to perennial species, 113 contained CDS found only in the P subgenome (128 CDS), while 169 contained CDS found only in the U subgenome (184 CDS), and only 58 orthogroups exclusive to perennials contained CDS from both the P and U subgenomes (69 + 63 CDS; Supplementary Tables 2a-d). Therefore, the total 346 perennial-type orthogroups included 200 and 250 CDS from the P and U subgenomes of *B. mexicanum*, respectively (Supplementary Tables 2a-c). These putatively perennial orthogroups were enriched in GO terms related to tolerance to stress conditions, cell wall development, endocytosis, and shoot system (Supplementary Fig. 7 and Supplementary Table 2d). In contrast, of the 135 orthogroups exclusive to annual species and *B. mexicanum*, 53 orthogroups presented sequences exclusive to the *B. mexicanum* P subgenome (68 CDS), 44 specific to its U subgenome (54 CDS), and 37 from both P and U subgenomes (44+42 CDS) (Supplementary Tables 2a, 2b, 2c and 2d). These orthogroups were enriched in GO terms related to cytokinesis, spindle organization, energy metabolism, reproduction, and shoot system development (Supplementary Fig. 8 and Supplementary Table 2d).

To assess whether *B. mexicanum* strongly influenced annual-associated orthogroup composition, we examined orthogroups containing sequences from the three annual species and one core perennial, but lacking the other core perennial (Supplementary Table 2c). Only 42 and 38 orthogroups contained *B. arbuscula* or *B. sylvaticum*, respectively, together with the three annuals. However, when *B. mexicanum* was also included, these numbers increased markedly to 138 and 143 orthogroups, respectively, indicating that *B. mexicanum* substantially contributes to orthogroups linking annual and perennial lineages.

### Mutation rates across growth habits

Between 11,084 and 14,819 orthologous gene pairs were detected among the four species/subgenome pairs (Supplementary Table 3). Between 2,373,883 and 3,334,746 4-fold degenerate sites were identified among these pairs. Mutation rate estimation showed that the four annual (sub)genomes exhibited higher (5.68E-09 for *B. distachyon* and *B. stacei)* or similar (5.75E-09 for the two subgenomes of *B. hybridum*) mutation rates, followed by lower but similar mutation rates obtained for perennial species (i.e., 3.79E-09 for *B. arbuscula* and *B. sylvaticum* genomes; 3.69E-09 for *B. mexicanum* subgenomes)(Supplementary Table 3).

### *Brachypodium* pan-gene dynamics

Based on the clustering results of the eight genomes/subgenomes from the six species, we identified a total of 12,545 core gene families with 147,238 genes, 5,689 softcore gene families with 57,122 genes, 8,304 dispensable gene families with 39,723 genes, and 1,305 private gene families totaling 3,807 genes (Fig. 3a). The proportion of each category of pan-gene count was largely consistent, showing average values of 57.2% (SD=5.36) for core genes, 20.5% (SD=3.11) for softcore, 15.0% (SD=2.58) for dispensable, and 1.3% (SD=1.21) for private of the total gene count (Supplementary Fig. 9 and Supplementary Table 4). Duplication mode analysis shows that most of the genes of *Brachypodium* are present as singletons (53.75%), followed by whole genome duplicated genes (15.65%), transposed genes (13.83%), tandem duplicated genes (9.58%), and proximal genes (7.02%) (Fig. 3b). The average gene length and coding sequence length of core genes was the longest (4072.05 bp for gene length and 1360.94 bp for coding sequence length), followed by softcore genes (i.e., 3387.64 bp; 1243.83 bp), dispensable genes (2128.35 bp; 792.16 bp) and private genes (1793.19 bp; 637.18 bp) (Figs. 3c and 3d).

**Fig. 3.**
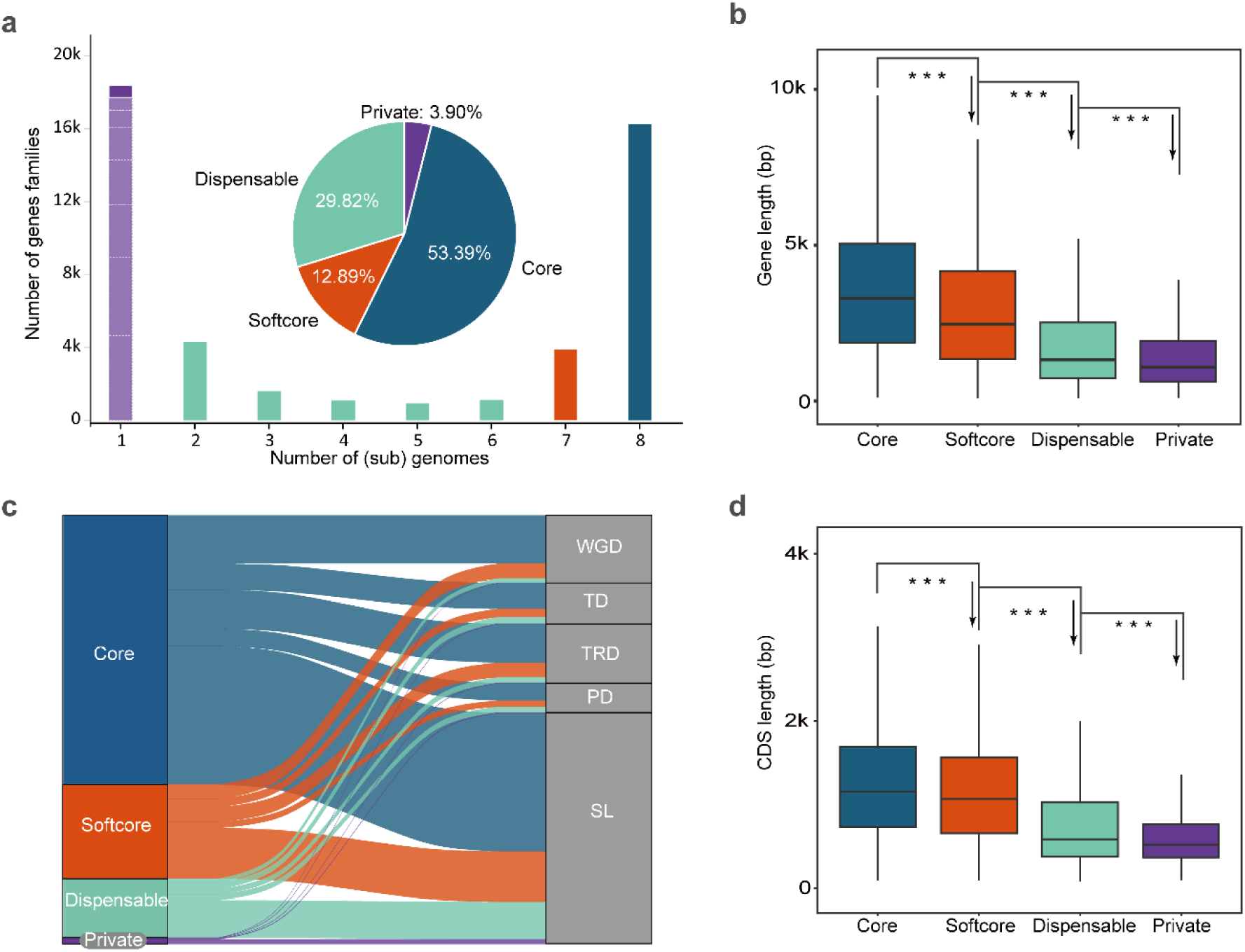
(**a**) Proportion of *Brachypodium* pan-genes in the core, softcore, dispensable, and private categories. Boxes filled with light purple show the unclustered genes for the eight (sub)genomes. (**b)** Proportions of the five types of duplicated genes, including genes from whole genome duplication (WGD), tandem duplication (TD), transposed duplication (TRD), proximal duplication (PD), and singleton genes (SL) across the four categories of pangenes (core, softcore, dispensable and private). Gene length (**c**) and CDS length (**d**) profiles of core, softcore, dispensable and private genes found in the *Brachypodium* pangenome. In Fig. c, the P-values for core vs softcore, softcore vs dispensable and dispensable vs private are <2.2E-16, <2.2E-16 and 8.588E-10, respectively. In Fig. d, the P-values for core vs softcore, softcore vs dispensable, dispensable vs private are <2.2E-16, <2.2E-16 and <2.2E-16 respectively. Three asterisks (***) indicate P < 0.001 (two-tailed Student’s t-test).

### Key role of transposable elements in the structure and evolution of the (sub)genomes of the *Brachypodium* pangenome

The proportion of repetitive regions varies greatly among the eight *Brachypodium* (sub)genomes ranging from 35.46% in *B. hybridum* S to 72.89% in *B. mexicanum* P (Fig. 4a and Supplementary Table 5). Most of the repetitive sequences are annotated as transposable elements, particularly LTRs (Fig. 4a). Total LTR proportions were highest in *B. mexicanum* P (60.00%) and *B. mexicanum* U (55.98%), followed by *B. sylvaticum* (27.59%), *B. hybridum D* (20.90%), *B. distachyon* (17.50%), *B. arbuscula* (16.44%), *B. stacei* (14.37%), and *B. hybridum* S (12.16%) (Supplementary Table 5). In general, the annual *Brachypodium* (sub)genomes have higher solo/intact LTR-RT ratios than do perennial genomes (Supplementary Table 6).

**Fig. 4.**
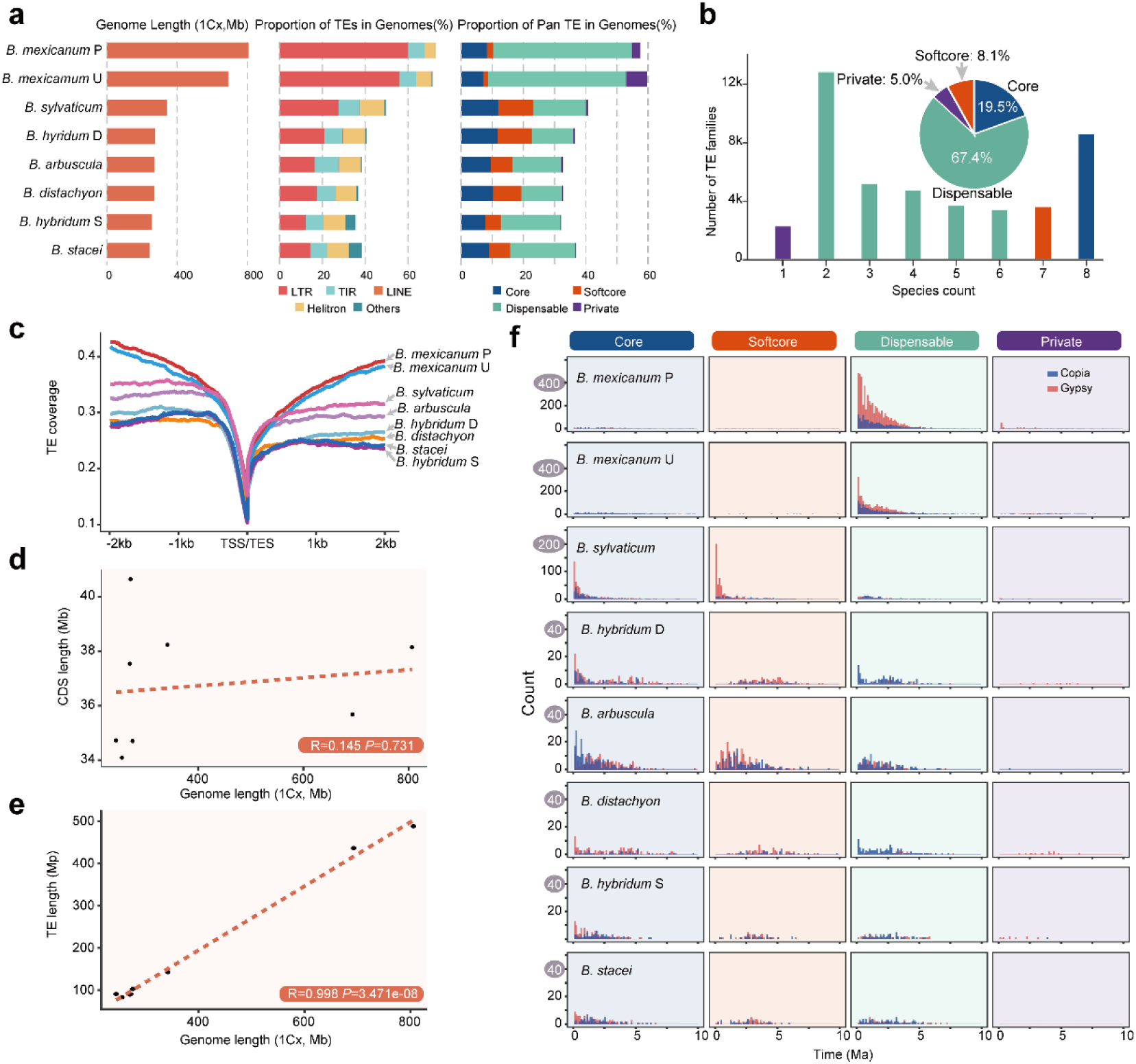
Genome sizes and transposable elements (TEs) of the eight *Brachypodium* (sub)genomes under study. (**a)** Genome length (1Cx, Mb), proportion of main TE families, and proportion of pan-TE categories in each sub/genome. (**b)** Proportion of pan-TEs in the core, softcore, dispensable and private categories of the *Brachypodium* pangenome. (**c**) TE coverage of the 2kb upstream (TSS, transcription start site) and 2kb downstream (TES, transcription end site) regions of coding-protein genes of the eight analyzed sub/genomes. (**d)** correlation between CDS length and genome length of the eight sub/genomes. (**e)** correlation between TE length and genome length of the eight subgenomes. For **d** and **e**, significance was determined using a two-sided Pearson’s correlation test. (**f)** Insertion times (Ma) of three categories of LTRs (Gypsy, Copia, Unknown) for core, softcore, dispensable and private TEs.

The Retand family was identified to be the most abundant transposable elements (Supplementary Figs. 10 and 11a-f) across all the eight (sub)genomes ranging from 11.80% in *B. mexicanum* subgenome P to 3.02% in *B. sylvaticum*, followed by Tekay family with the highest proportion identified in the two subgenomes of *B. mexicanum* (i.e., 5.82% in subgenome P and 5.06% in subgenome U). Gymco I and II TEs were identified exclusively in *B. stacei* and *B. hybridum* S subgenome, and Chlamyvir and TatII retrotransposons exclusively in *B. stacei*. The Ogre family was identified in all species except *B. distachyon* (Supplementary Fig. 10).

Analysis of pan-TE families revealed that dispensable TEs accounted for 67.4% of whole TE families, followed by 19.5% for core TEs, 8.1% for softcore TEs and 5.0% for private TEs (Fig. 4b). Retand were the most abundant TE family in softcore (52.1%), dispensable (42.0%) and private (70.1%) TEs, and the second most abundant in core TEs (17.9%) after Tekay (32.5%) (Supplementary Fig. 12). The pan-TE proportions for the eight (sub)genomes were also largely consistent, except the *B. mexicanum* U subgenome, which contained a higher proportion of private TE families (17.02%) than the others (0.28-5.69%) (Supplementary Fig. 13; Supplementary Table 7).

When pan-TE categories were compared based on genomic coverage, dispensable TEs were also the most abundant, especially the two subgenomes of *B. mexicanum*, accounting for 44.51% and 44.35% for subgenome P and subgenome U, respectively, whereas the remaining six (sub)genomes showed much lower proportions, ranging from 13.31% to 20.96% (Fig. 4a, Supplementary Table 8). Core TE is the second most abundant category, with proportions ranging from 7.11% in U subgenome of *B. mexicanum* to 11.92% in *B. sylvaticum*. Softcore TEs were generally the third most abundant TEs except in the two subgenomes of *B. mexicanum*, which has higher proportion of private TEs and softcore TEs. In all four categories of pan-TEs, the proportions of *B. sylvaticum* are higher than those of *B. arbuscula* (Supplementary Table 8).

The TE coverage of 2 kb upstream and downstream regions of the protein-coding genes (Fig. 4c) showed the same size order as that of their corresponding genome size (Fig. 4a). CDS length of protein coding genes exhibited a low positive correlation with genome size that was not statistically significant (R=0.145; P=0.731) (Fig. 4d), whereas total TE length across the eight (sub)genomes showed a strong positive correlation with genome size (R=0.998; P=3.47E-08) (Fig. 4e).

### Dynamics of TEs, insertion times, recent bursts, and phylogeny of the *Gypsy* and *Copia* LTR-retrotransposons in *Brachypodium*

The LTR insertion time based on each category of pan-TE showed that the ancestral BmexP subgenome had a significantly higher accumulation of *Gypsy* elements than BmexU over the last 5 million years (My; Fig. 4f). Furthermore, private *Gypsy* TEs in the BmexP subgenome experienced a recent surge in TE insertions. In addition, most of the *Gypsy* TEs inserted to the *B. mexicanum* subgenomes are dispensable (Fig. 4f). Most of the TE insertions in *B. sylvaticum* correspond to the core and softcore TEs, which peak in the last 5 Ma. As for the dispensable TEs, the insertions were accumulated more gradually and correspond to *Copia*. Unlike the other species, *B. arbuscula* shows a recent peak accumulation of *Copia* for the core category, compared to the accumulation of LTR-*Gypsy* insertions of the core and softcore present in the annual species and *B. sylvaticum* (Fig. 4f). Annual species and *B. arbuscula* showed much lower accumulation of retrotransposon insertions in the last 5 Ma compared to those accumulated in *B. mexicanum* and *B. sylvaticum* (Fig. 4f).

The unrooted RT domain-based phylogeny of *Copia*-like retrotransposons (Supplementary Fig. 14a) revealed the greatest expansion of the Ale, Bianca, Ikeros, Ivana, SIRE, and TAR families in *B. mexicanum* P, followed by *B. mexicanum* U and *B. sylvaticum*, which correspond to three of the largest (sub)genomes analyzed in this study. This expansion is especially marked in the SIRE and Ikeros retrotransposons of the ancestral *B. mexicanum* P. Furthermore, the unrooted phylogeny based on the RT domain of *Gypsy*-like retrotransposons (Supplementary Fig. 14b) identified the greatest number of expansions of the CRM, Ogre and Reina families in *B. mexicanum* P, followed by *B. mexicanum* U and *B. sylvaticum*. Most of the diversity of Ogre elements occurred in *B. mexicanum* P, with a much smaller presence in *B. mexicanum* U. Interestingly, for the most abundant Retand families, *B. mexicanum* U had the highest number of expansions, followed by *B. sylvaticum*, while the expansion of *B. mexicanum* P was minimal, although the Retand family took up the highest proportion in *B. mexicanum* P (Supplementary Fig. 14b).

## Discussion

This study displays the representative pan-genome of the genus *Brachypodium*, integrating previously characterized genomes from the annual species *B. distachyon*, *B. stacei*, and *B. hybridum*^8,9,10,16,26,27^, the perennial *B. sylvaticum*^11^, and the newly assembled chromosome-level genomes of the perennial species *B. arbuscula* and the allopolyploid *B. mexicanum* (Fig. 1 and Supplementary Table 1). Our sampling enables comparative analyses at both genomic and subgenomic levels, revealing major chromosomal rearrangements, genic PAV, and TE-family distributions across representative lineages of the genus (Figs. 2-4).

Both *B. mexicanum* subgenomes carry lineage-specific rearrangements, but the more recently derived U subgenome appears to have accumulated more derived structural variations within the phylogenomic framework inferred here (Fig. 2). Our comparative genomic analyses also refine earlier comparative chromosomal barcoding (CCB) hypotheses for subgenomes^12,13^. Some CCB-inferred syntenic patterns in the BmexP subgenome (Bmex A1.1^13^) and the BmexU subgenome (Bmex A1.2^13^) are corroborated by genome-scale synteny (Fig. 2c, and Supplementary Figs. 5, 6), whereas others are not supported. The discrepancies likely reflect limited probe specificity and the inability of barcoding data to recover complete chromosomal syntenic blocks in earlier analyses. Complete chromosome-level assemblies therefore allow a more accurate resolution of inversions and translocation between the P and U chromosomes of *B. mexicanum* (Fig. 1 and Supplementary Figs. 5, 6). Thus, Bmex7P and Bmex7U are highly collinear (Fig. 2c and Supplementary Fig. 5), contradicting the earlier inference of a reciprocal translocation between Bmex7U and Bmex3U^13^. Synteny and subgenome assignment also support an inversion in Bmex2P rather than in Bmex2U^13^, because the other analyzed *Brachypodium* genomes are collinear with the Bmex2U chromosome (Fig. 2c). In addition to the telomeric translocation in Bmex5U detected by CCB^13^, our analyses identify a second translocation on this chromosome (Fig. 2c and Supplementary Fig. 5). Together, these results indicate that many ancient structural variants within the x = 10 karyotypes arose during the transition from the ancestral P subgenome to the more derived U subgenome, with additional rearrangements within the P subgenome itself, including the large inversion on Bmex2P (Fig. 2c and Supplementary Figs. 5, 6).

Previous genome size estimates revealed that genome sizes among *Brachypodium* perennials vary more than seven-fold^13,18^. The ancestral *B. mexicanum* (x=10P + 10U) harbors the largest subgenome sizes within the genus, being approximately three times larger than those of the other allotetraploid, *B. hybridum* (x=10S+5D) (Figs. 2 and 4a). In contrast, *B. sylvaticum*, a recently evolved core-perennial diploid species (x=9G) that diverged after the split of *B. arbuscula*, around 4.78 million years ago (Ma) (Fig. 2b), has a genome 88 Mb larger and with 11.33% higher repeat content than *B. arbuscula* (Supplementary Table 5). The expansions of both genomes (Bmex, Bsyl) resulted from proliferations and retention of transposable elements, but at different evolutionary times (Fig. 4 and Supplementary Figs. 10, 11) and under different biological and cytological constraints.

Our repeat elements analysis based on chromosome level genome assembly revealed a higher proportion of repeat contents for all eight (sub)genomes of the six analyzed *Brachypodium* species (Fig. 4 and Supplementary Figs. 10, 11) than was previously documented^18^. Our data are consistent with previous findings indicating that the Retand retrotransposon is the most abundant TE family in *Brachypodium* genomes (Supplementary Figs. 10 and 12)^18,28^. However, our results also showed that Retand retrotransposons exhibit different dynamics within the two *B. mexicanum* subgenomes. Although the highest proportion of Retand retrotransposons is identified in the P subgenome (Supplementary Fig. 10), our gene tree based on the *Gypsy*-RT domain showed that the amount of P subgenome with an RT domain is the lowest (Supplementary Fig. 14b), indicating that most Retand elements are not autonomous in the P subgenome of *B. mexicanum*. In contrast, the highest number of Retand retrotransposons with an RT domain was identified for the U subgenome of *B. mexicanum*. For the remaining *Gypsy* and *Copia* retrotransposon lineages, *B. mexicanum* P showed the greatest diversity for those with RT domains (Supplementary Figs. 14a and 14b). We also found that Ogre retrotransposons are present in seven *Brachypodium* (sub)genomes, with the exception of that of *B. distachyon*. Although previously reported only in *B. mexicanum*^18^, our analyses reveal their occurrence also in *B. arbuscula*, *B. hybridum*, *B. stacei*, and *B. sylvaticum*, albeit at considerably lower proportions (Supplementary Figs. 10 and 14b).

TE length is strongly positive related to genome length across the eight (sub)genomes, in contrast to the CDS length (Figs. 4c and 4e). Our data corroborate that insertions of the most abundant LTRs cause variation in the sizes of *Brachypodium* (sub)genomes and their potential regulations, as in other species^26,29,30^. However, research on whether this variation was due to insertions of shared LTRs or to species-specific LTRs is scarce. Our pan-TE analysis here gives a different perspective showing that genome size variation could be driven by both shared TE variation and species-specific TE insertions. The three largest (sub)genomes of *Brachypodium* show distinct temporal and spatial TE dynamics. In *B. mexicanum*, the two largest subgenomes contain ∼44% dispensable TEs (Fig. 4a and Supplementary Table 8), which experienced their most active insertion events since ∼5 Ma (Fig. 4f), while LTR insertions in other plants such as maize (LTR insertion time 0.12-5.15 Ma), wheat (retrotransposon insertion time in 5A chromosome 0.8-4.5 Ma), rice (solo-LTR insertion time <4Ma) and cotton (LTR insertion times in A and D subgenomes 2 Ma) have been dated more recently^31,32^. In *B. sylvaticum*, although dispensable TEs also dominate, their activity, recorded since ∼3 Ma for LTR transposons, mainly involves shared core/soft-core families. PAV-rich regions have been associated with TE-rich regions in several species such as lettuce and soybean^33^, consistent with the high presence of dispensable TEs detected in the present study. In addition, different TE families have evolved to occupy specific genomic niches in centromeric, pericentromeric, or telomeric regions of chromosomes. LTR-*Gypsy* and *Copia* elements are usually located in centromeric and pericentromeric regions^34^, as shown in diploid *Brachypodium* and in other species with large genomes and centromeric regions such as tomato or melon^35^. However, LTR-*Gypsy* and *Copia* elements can also be found in telomeric and subtelomeric regions^36^, as observed in *B. mexicanum*, whose larger genome size of its P and U subgenomes is explained by the predominance of TE Retand and Tekay throughout its entire chromosomal length (Supplementary Figure 11a) and the next most abundant LINE elements were more sparsely dispersed. In contrast, the smaller genomes of *B. stacei* and *B. distachyon* showed preferentially the Retand and Tekay TE families in the centromeric regions, and LINE elements were distributed across the chromosomes (Supplementary Figs. 11e and 11f).

The large subgenome sizes of *B. mexicanum* may be explained by the “polyploid genome shock” hypothesis, which proposes that the sudden combination of distinct genomes leads to reshuffling and mobility of the repetitive elements in hybrid and polyploid plants and followed proliferation of repetitive elements^37^. Our results revealed that more LTR insertions were identified for the more recently formed subgenome D of *B. hybridum* than its diploid progenitor species *B. distachyon* and the subgenome S of *B. hybridum* also contain more conserved *Gypsy* insertions than its diploid progenitor species *B. stacei* (Fig. 4f). Previous studies have shown that selfing species often exhibit elevated TE copy numbers and insertion frequencies, largely due to reduced selection efficacy against TEs in the absence of heterozygosity-dependent ectopic recombination^38^. *Brachypodium mexicanum* and *B. sylvaticum* are the only known self-compatible species within the *Brachypodium* perennials^6,20^, potentially subjecting them to relaxed purifying selection against TEs and facilitating their accumulation and, as perennial species with longer generation times, both likely experience reduced efficacy of TE elimination compared to annual species. This pattern is consistent with our observation that, at equivalent ploidy levels, perennial *Brachypodium* species tend to have larger genomes than their annual counterparts (Fig. 4a and Supplementary Table 1)^13,18^.

The high abundance of TEs, whether as a cause or a consequence of meiotic crossover suppression in repeat-rich genomic regions, appears to be a far more complex and variable phenomenon across organisms. In plants, crossovers are generally suppressed within pericentromeric and centromeric regions, showing a strong negative correlation with DNA methylation levels and TE density, although the strength of this relationship depends on both the transposon family and the species,^39,40^. Similar to other angiosperm groups, where genomes of ancestral lineages show a significantly higher genome size than those of more recently evolved lineages (e. g., *Hesperis* clade; Asteraceae, *Lactuca*)^41,42^, the *B. mexicanum* subgenomes have maintained a large number of TEs, mostly LTRs. This could be due to the reproductive isolation of this ancient lineage, which does not cross with other *Brachypodium* species^20^, thus avoiding the mechanism of transposon deletion through hybridization and recombination. In addition, *B. mexicanum*, like other large plant genomes which experienced less transposon purging and could therefore maintain the excess of TE load and possess large centromeric regions. These larger centromeric region could compensate the respective lengths of chromosome arms and centromeres and facilitate the correct segregation of chromosomes during cell division^43^. Furthermore, in some plants, the reduction in genome size is associated with an adaptive transition to an annual life cycle, irrespective of the species’ age^44^. This variation—proliferation/reduction—of TEs and the transition from perenniality to annuality impacted the chromosome architecture and size of the analyzed *Brachypodium* (sub)genomes (Figs. 2 and 4). However, the relatively small percentage of repetitive elements in *B. arbuscula*, even lower than that of the annual diploids *B. distachyon* and *B. stacei*, and their relatively high proportion in the *B. sylvaticum* genome (Supplementary Table 5), points not only to self-fertility but also to other potential biological constraints, such as adaptation to broad, humid landscapes and its ubiquitous infection with its fungal endophyte *Epichloë sylvatica*^45^, as factors that may also have favored the expansion of recently acquired TEs (Fig. 4).

Regarding the total number of protein-coding genes, *B. arbuscula* has fewer genes than *B. sylvaticum* and also fewer than the annual *B. distachyon*, while the allotetraploid *B. mexicanum* shows a slightly higher number of genes than *B. hybridum* (Supplementary Table 1). Although the number of genes predicted per haplome does not vary considerably between genomes, some PAVs were exclusive for the P and U subgenomes of *B. mexicanum*, for annual species, for core perennial species, or for combinations thereof (Supplementary Table 2). Substitution rates were lower for perennial and higher for annual subgenomes (Supplementary Table 3), an expected correlation with the lengths of their respective reproductive cycles. For the pan-gene analysis, core gene families were the most abundant, followed by variable gene families (softcore and dispensable) and private gene families, and this downward trend from core to private gene families is shared by all eight (sub)genomes (Fig. 3 and Supplementary Fig. 9). However, subgenomes P and U of *B. mexicanum* accumulate the highest proportion of private genes and the lowest of core genes (Supplementary Table 4), in concordance with their earliest and subsequent diverging positions in the *Brachypodium* phylogeny constructed from shared genes (Fig. 2b).

The morphological, physiological and genomic features of *B. mexicanum* (intermediate between those of the annual and perennial *Brachypodium* species^6)^, together with the evolutionary divergences of its P and U subgenomes (close to those of the annual species; Fig. 2b), lead us to hypothesize the involvement of at least one annual species as an ancestral progenitor, or alternatively, a transition from an annual to a short-lived perennial during its evolutionary trajectory. This study supports the presence of genes in BmexP and BmexU chromosomes closely related to those of the annual species. We identified 135 orthogroups containing sequences from the three annual species and *B. mexicanum*, but none from *B. arbuscula* or *B. sylvaticum* (Supplementary Tables 2a, 2b, and 2d). This contrasts with the 42 and 38 orthogroups detected in the same analysis when *B. arbuscula* and *B. sylvaticum*, respectively, were included among the annual species (Supplementary Table 2c). The large genome size of *B. mexicanum* may partially account for these differences; however, even when orthogroups (Annuals + Bmex vs. Barb + Bsyl) are examined separately for the P and U subgenomes, both subgenomes still exhibit a higher number of “annual”-type orthogroups (37 containing sequences from both subgenomes, 53 from P only, and 44 from U only) (Supplementary Table 2d). Considering the number of sequences assigned to each orthogroup category (Perennial-specific vs. Annuals + Bmex-specific), subgenome P contains 200 sequences grouped within the perennial orthogroup, whereas subgenome U contains 250; in contrast, subgenome P contributes 112 coding sequences to the annual orthogroup, compared with 96 from subgenome U (Supplementary Table 2b). These patterns indicate that the more ancestral subgenome P harbors a relatively higher proportion of genes predominant in annual species of the genus, whereas subgenome U contains a greater number of sequences characteristic of perennial species (Supplementary Figs. 7 and 8).

Regarding the hypothetical life forms of the ancestors of *B. mexicanum*, presumably extinct but with genomes present today only in the orphan subgenomes of this meso-allopolyploid (Fig. 2b and 2c), our genic data supports a presumed annuality of ancestor P and a presumed perenniality of ancestor U (Supplementary Tables 2a, 2b and 2d). Spontaneous and artificial crosses between annual and perennial species are common in some Pooideae lineages, resulting in sterile interspecific/intergeneric hybrids, or allopolyploids after WGD^46,47^. In *Brachypodium*, allopolyploids account for more than half of the extant species^6,7^, and crosses have occurred between the annual species *B. distachyon* and perennial species of the core-perennial clade, either naturally or synthetically^6,20^, in some cases producing offspring with low fertility^20^. Due to the prolonged evolutionary and geographic isolation of *B. mexicanum*, crosses with extant annual or perennial congeners may be unfeasible. However, expanding the number of other genomes incorporated into the *Brachypodium* pangenome will be essential to test hypotheses about the origins and progenitors of *B. mexicanum* and other polyploid species within this model genus.

## Methods

### Taxon sampling, genome sequencing

*Brachypodium arbuscula* (accession ID: Barb1) were sampled in Spain: Canary Isles, La Gomera, Vallehermoso, and *B. mexicanum* (accession ID: Bmex347) was sampled in Hidalgo, Sierra de Pachuca, Mexico). Herbarium vouchers of these two accessions were preserved at the Unizar herbarium. We sequenced *B. arbuscula* (BARB1) and *B. mexicanum* (BMEX347) using a whole genome shotgun sequencing strategy and standard sequencing protocols. Sequencing reads were collected using Illumina and PACBIO platforms. Illumina and PACBIO reads were sequenced at the HudsonAlpha Institute in Huntsville, Alabama and the Department of Energy (DOE) Joint Genome Institute (JGI) in Berkeley, California, USA. Illumina reads were sequenced using the Illumina HiSeq-2500 and NovaSeq S4 platforms, and the PACBIO reads were sequenced using the Sequel II platform. For each of the two species, one 400bp insert 2×250 Illumina fragment library (388.63x coverage for *B. arbuscula*, 60.66x coverage for *B. mexicanum*) was sequenced along with one 2×150 HiC library (209.31x coverage for *B. arbuscula*, 26.90x coverage for *B. mexicanum*). Prior to assembly, Illumina fragment reads were screened for phix contamination. Reads composed of >95% simple sequence were removed. Illumina reads <50bp after trimming for adapter and quality (q<20) were removed. The final read sets consist of 848,070,262 high-quality Illumina reads, covering 597.94x for *B. arbuscula* and of 642,594,486 high-quality Illumina reads, covering 87.56x of bases for *B. mexicanum* respectively. For the PACBIO sequencing, a total raw sequence yield of 68.23Gb (113.71x coverage) and 163.48 Gb (90.82x) for *B. arbuscula* and *B. mexicanum*, respectively.

### Genome assembly and annotation

Genome assembly based on the sequenced reads entails genome assembly based on PacBio Sequel long reads, subsequent polishing, chromosome anchoring based on HiC reads, manual curation, redundancy reduction and final polishing using Illumine reads. Gene annotation of *Brachypodium mexicanum* and *B. arbuscula* was performed using an integrated pipeline combining transcriptome evidence, protein homology, and *ab initio* gene prediction. Detailed steps on genome assembly and annotation are provided in Supplementary Data 1.

### *Brachypodium mexicanum* subgenome assignment and synonymous distance estimation of paralogous genes

Due to controversy about the auto– or allo-polyploid nature of the two subgenomes of *B. mexicanum*^13^, we employed both a K-mer based method using SubPhaser v.1.2.6^48^, and a gene phylogeny-based method using an updated version of PhyloSD^13^, to identify and assign the assembled chromosomes of *B. mexicanum* into its subgenomes. For the SubPhaser assignment, the kmers were counted by Jellyfish v. 2.2.10^49^, followed by identification of differential kmers among homoeologous chromosomes and clustering of chromosomes into subgenomes P and U by a K-Means algorithm. All the analyses were implemented using the default parameters in SubPhaser. For PhyloSD analysis, a total of 9,627 clusters of ortho/homeologous coding sequences (CDS) selected by Orthofinder from the available reference genomes of *B. distachyon*, *B. hybridum* (subgenomes S and D), *B. stacei* and *B. sylvaticum*, and the new genomes of *B. arbuscula* and *B. mexicanum* (subgenomes P and U) were analyzed. Each of the clusters had to include a sequence of the diploid species and at least one of *B. mexicanum*. Each cluster was aligned using Mafft v.7.490^50^ and filtered using TrimAl v.1.4.1^51^ to be subsequently analyzed by PhyloSD^13^.

To estimate synonymous substitution rates, the paralogous genes of the *B. arbuscula* genome and of within and between *B. mexicanum* subgenomes were identified using synteny-based method implemented in Wgdi v.0.6.5^52^. Briefly, synteny blocks within *B. arbuscula* genome and *B. mexicanum* subgenomes were extracted and the synonymous substitution distances distribution between paralogous genes were estimated using PAML v.4.10.7^53^ implemented in Wgdi. We further dated the timing of the ancient whole genome duplication shared by all Poaceae species (rho WGD)^54^ based on the gaussian fitted peak *Ks* values implemented in Wgdi and mutation rates estimated with the equation T=*K*/2u, where *K* is the fitted peak *Ks* value, u is the mutation rate estimated and T is the time. Similarly, the divergence time between the two subgenomes of *B. mexicanum* was also estimated.

### Phylogenomics and molecular dating

We analyzed four available *Brachypodium* reference genomes (*B. distachyon* (Bd21), *B. stacei* (ABR114), *B. hybridum* (ABR113), *B. sylvaticum* (Ain1)) and the newly generated *B. arbuscula* (Barb1) and *B. mexicanum* (Bmex347) genomes, plus the genomes of outgroup species including *Oryza sativa* var. Japonica^55^, *Setaria viridis*^56^ and *Triticum urartu*^57^ downloaded from NCBI (https://www.ncbi.nlm.nih.gov/). We divided the two subgenomes of *B. mexicanum* and *B. hybridum* allotetraploids and treated them as separate lineages when the genes were clustered for comparative genomic analysis. We first filtered out those genes with less than 100 amino acids. Single copy orthologous genes were identified using OrthoFinder v.2.5.5^58^. Protein coding sequence of each identified single copy gene cluster was aligned using MAFFT v.7.490 with the automatic model, followed by alignment refinement using TrimAl with “-gt 0.6” parameter.

Phylogenetic inference was conducted using both concatenation and coalescent-based approaches. Multiple sequence alignment (MSA) from concatenated gene alignments was analyzed using IQ-TREE2 v.2.3.0^59^ under the best-fit model GTR+F+I+G4 selected by the program using “-m TEST” parameter based on BIC (Bayesian Information Criterion) with 1,000 ultrafast bootstrap replicates. For the coalescent approach, individual gene trees were inferred with IQ-TREE2 for each of the 2,131 genes used for phylogenomic reconstruction and the species tree was estimated with ASTRAL-III v.5.7.8^60^.

Due to the lack of fossil records for *Brachypodium* and its closely related taxa, divergence times of these taxa were dated using Beast v.2.7.1^61^ based on secondary calibrations. We filter out those genes that showed incongruent topologies regarding the placement of the two subgenomes for both *B. hybridum* and *B. mexicanum* with the species tree thus to have a more reliable dating. The remaining 832 genes were concatenated into an MSA, which was analyzed using Beast2 with the GTR mutation model, a lognormal relaxed clock, Yule tree models, and an exponential birth rate model. We constrained the crown age of BOP+PACMAD (normal prior mean=55.0Ma, SD=0.5Ma), the crown age of *Brachypodium* + *Triticum* + *Oryza* (BOP) group (normal prior mean=51.7Ma, SD=1.9Ma), and the crown age of Pooideae (*Triticum* + *Brachypodium*; normal distribution mean=33.2Ma, SD=9.5Ma). A total of one billion MCMC generations were run. The convergence of each parameter was checked with TRACER v.1.7.2^62^, ensuring that Effective Sample Size (ESS) > 200. Maximum clade credibility (MCC) trees were computed using TreeAnnotator v.1.7.2^61^ with the first 10% of trees discarded as burn-in.

We also compared and visualized the synteny relationship among the *Brachypodium* genomes using JCVI v.1.4.6 pipeline^63^. The pairwise synteny among all pairs of *Brachypodium* genomes was searched using “python-m jcvi.compara.catalog ortholog genome1 genome2”, followed by synteny blocks with command “*python-m jcvi.compara.catalog screen –minspan=30 ––simple genome1.genome2.anchors genome1.genome2.anchors.new*”. The filtered synteny blocks were visualized using the command “*python –m javi.graphics.karyotype seqids layout*”.

### Mutation rate estimation

As the mutation rate varies between closely related annual and perennial plants^64^, we estimated the mutation rate for the annual and perennial *Brachypodium* species separately based on the equation r=u/2T, where u is the proportion of mutational sites between two species and T is the divergence time of the two species. The one to one (ortho-homeologous genes for four diploid species genomes and the respective homeologous subgenomes of each allotetraploid, and paralogous genes for the respective subgenomes of the two allotetraploids) between pairs of genomes or subgenomes were determined based on the cluster information of OrthoFinder. The protein sequences of these pairs were aligned using MAFFT and used to generate the codon-based alignment using Pal2al v.14^65^. This step was followed by the identification of 4-fold degenerate sites using a custom script. Mutation rates were calculated by dividing the proportion of mutational sites by the divergence times estimated by Beast2.

### Construction of the *Brachypodium* pangenome

Gene orthogroups identified by OrthoFinder were classified into core (present in all eight genomes/subgenomes analyzed), softcore (present in 7), dispensable (present in ≥2 and ≤6) and private (present only in one genome/subgenome) based on the number of genomes/subgenomes contained in each cluster. In the pangenome analysis, we reran Orthofinder based only on the eight ingroup (sub)genomes including *B. arbuscula, B. distachyon, B. stacei, B. sylvaticum* and the respective subgenomes of *B. hybridum* (S and D) and *B*. *mexicanum* (P and U) without the length filtering step employed in the phylogenomic analysis. The gene and coding sequence (CDS) length of these three pangenome categories of genes were also compared. We further classified the duplication type of the genes from the six species based on their comparison with *Oryza sativa* using DupGen_finder v.1.0.0^66^.

Leveraging the reference genomes of three annual and three perennial plants, including the perennial *B. mexicanum* with a habit intermediate between annuals and perennials, potential genes related to perennial-to-annual (and/or vice versa) transition syndromes were explored. First, orthogroups were established from the CDS of the six *Brachypodium* species using the get_homologues-x86_64-20251130 software^67,68^ and the parameters –m local –M –A –t 0 –S 95 (min % sequence identity of 95%). The Get_Homologues-est scripts *compare_clusters.pl* (–m –n) and *parse_pangenome_matrix.pl* (–m clusters_cds_t0_S95/pangenome_matrix_t0.tab –g) were used to generate the pangenome matrix and to analyze pan-genome sets, respectively, enabling the identification of genes/CDSs present in annual species but absent in perennials, and vice versa. In addition, we also performed functional annotation of the *B. mexicanum* genes using the online portal of eggNOG-mapper (http://eggnog5.embl.de/#/app/seqscan, accessed in June, 2025)^69^. Gene enrichment of those gene families detected were carried out using topGO^70,71^ package emplemented in R. The enriched GO terms were further summarized in Revigo (http://revigo.irb.hr)^72^.

### Repeat annotation, pan-TE analysis and LTR insertion time estimation

We annotated the repeat sequence of eight *Brachypodium* (sub)genomes (six species) including *Brachypodium arbuscula*, *B. stacei*, *B. distachyon*, and *B. sylvaticum*, and two subgenomes each of *B. hybridum*, and *B. mexicanum* using EDTA v.2.1.0^73^. For each sub/genome, the *solo* and *intact* LTRs were identified by *solo_finder.pl* and *intact_finder_coarse.pl* provided by LTR_retriever v.3.0.4^74^, respectively, and the *solo/intact* ratio was subsequently estimated for each sub/genome. To conduct pan-TE analysis, the TE libraries identified by EDTA were clustered by OrthoFinder and classified into core, softcore, dispensable and private TE families following the previously defined class presence-criterion for protein coding-genes (core: 8; softcore: 7; dispensable: 2-6; private: 1). The TE length and insertion times were explored to inform their contributions to genome size variation. Insertion times (T) of different LTRs were calculated based on the equation T=D/2r, where D is the divergence level of intact LTRs and r is the mutation rate, which was estimated previously for each genome. To examine TE dynamics, LTR-retrotransposons were classified into the major superfamilies *Gypsy* and *Copia* respectively, their RT domains extracted and aligned with the *concatenate_domains.py* script from TEsorter v.1.4.7^75^. RT sequence-based phylogenies were inferred using IQTREE2 with 1,000 ultrafast bootstrap replicates and visualized using ITOL (https://itol.embl.de/). Based on EDTA output, we quantified the contribution of each TE subfamily to the total genome length. We also analyzed the correlation between total TE length, total coding sequence length and genome size using the linear model (lm function) implemented in R v.4.3.0^76^. The significance of these correlations was determined using two-sided Pearson’s correlation test implemented in R v.4.3.0.

## Data availability

Genome assemblies and annotations can be downloaded from Phytozome [https://phytozome.jgi.doe.gov/]. The direct links for the *B. mexicanum* and *B. arbuscula* reference genomes are [https://phytozome-next.jgi.doe.gov/info/Bmexicanum_v2_1] and [https://phytozome-next.jgi.doe.gov/info/BarbusculaBARB1_v3_1], respectively. Supporting data are available in the Supplementary materials and at the Github repository https://github.com/Bioflora/BmexicanumBarbuscula. The upgraded PhyloSD version is available on [https://github.com/eead-csic-compbio/allopolyploids].

## Authors’ contributions

D.L.D, JV and PC obtained funding for the project. PC, CC and RS designed the study. KB provided overall project management, AL conducted RNA analyses, MK and JE performed genome sequencing, CC conducted the comparative genomics, phylogenomic, TE’s and pangenomic analyses, RS performed the PAV analyses, BC and RS conducted the PhyloSD analyses, D.L.D and PC conducted and supervised the analyses. CC, RS and PC wrote the original draft, and all authors contributed to writing.

## Authors’ information

Chunlin Chen

Ruben Sancho

Miguel Campos

Bruno Contreras Moreira

John P. Vogel

David L. Des Marais

Pilar Catalan

## Supplementary Figures

**Supplementary Fig. 1.**
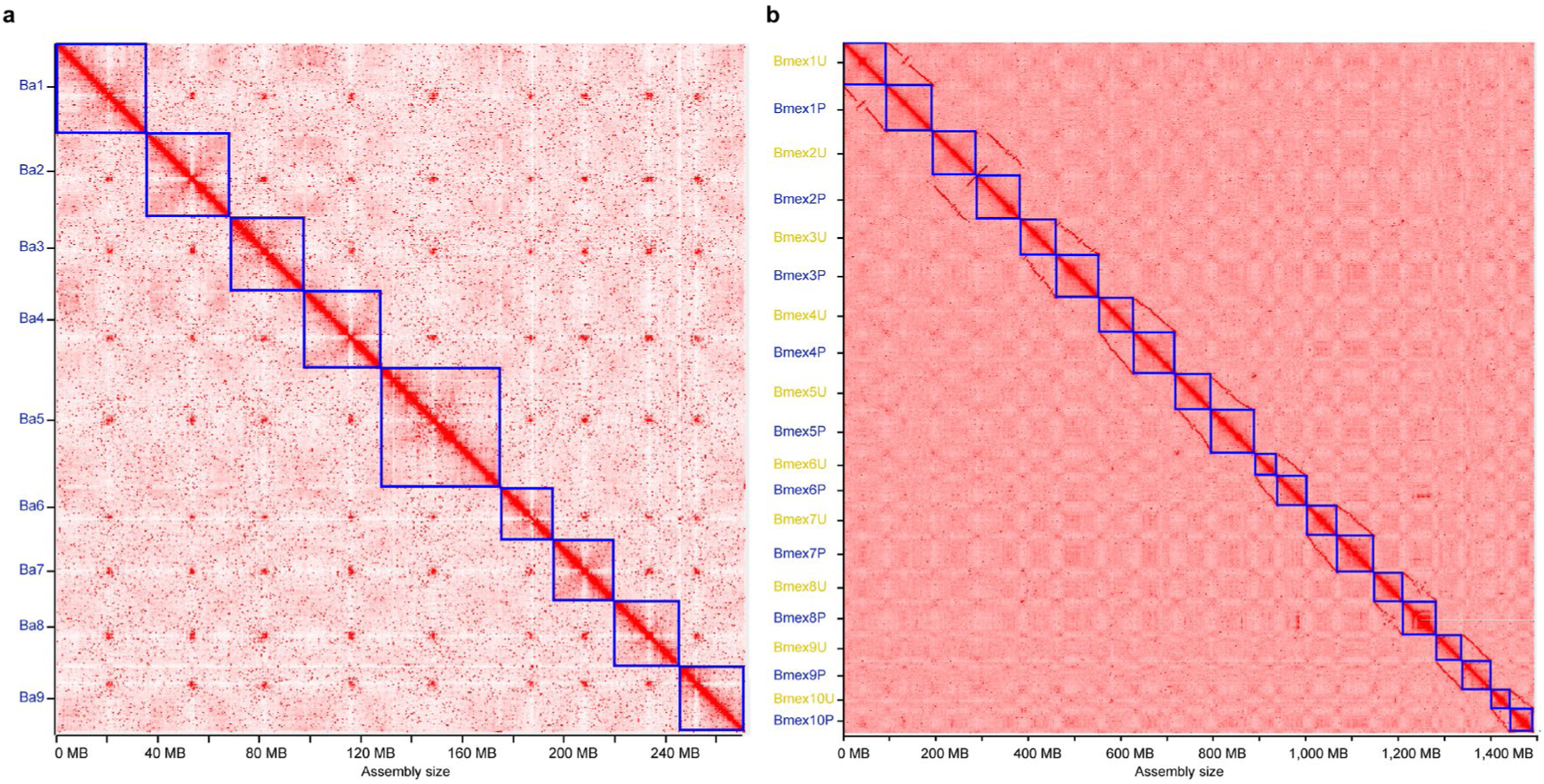
HiC interaction maps of *Brachypodium arbuscula*. **(a)** and *B. mexicanum* **(b)** genomes. For *B. mexicanum*, the blue and yellow letters indicate chromosomes assigned to P and U subgenomes, respectively.

**Supplementary Fig. 2.**
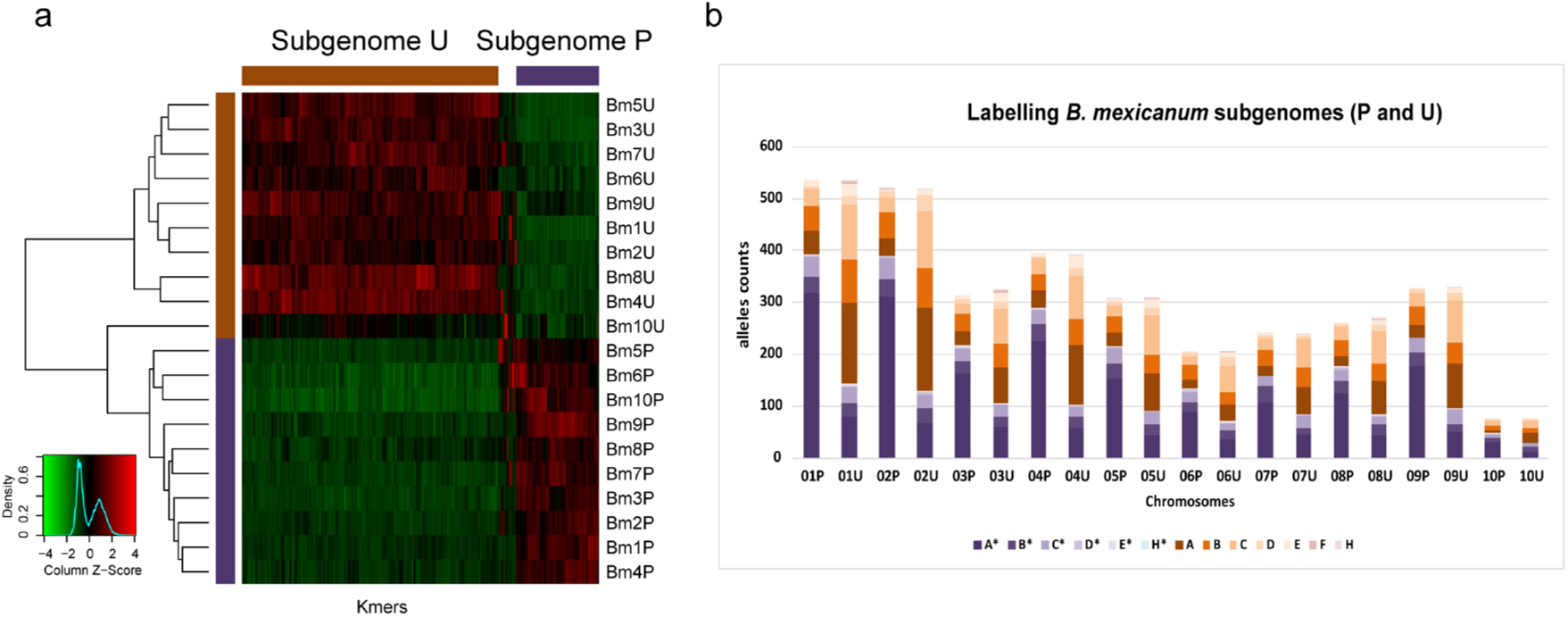
*Brachypodium mexicanum* P and U subgenomic assignments. (**a**) K-mer based approach using Subphaser^1^. Repetitive kmers that are uniquely expanded in specific subgenomes, showing high Z scores (red colors) in their respective chromosomes, differentiate the P and U subgenomes. (**b**) Maximum likelihood phylogeny-based approach using PhyloSD^2^. Letters with asterisks (blue range colors) indicate ancestral-labelled alleles and those without asterisks (orange range colors) more recently evolved-labelled alleles. The P subgenome chromosomes are enriched with ancestral alleles and the U subgenome chromosomes with recently evolved alleles.

**Supplementary Fig. 3.**
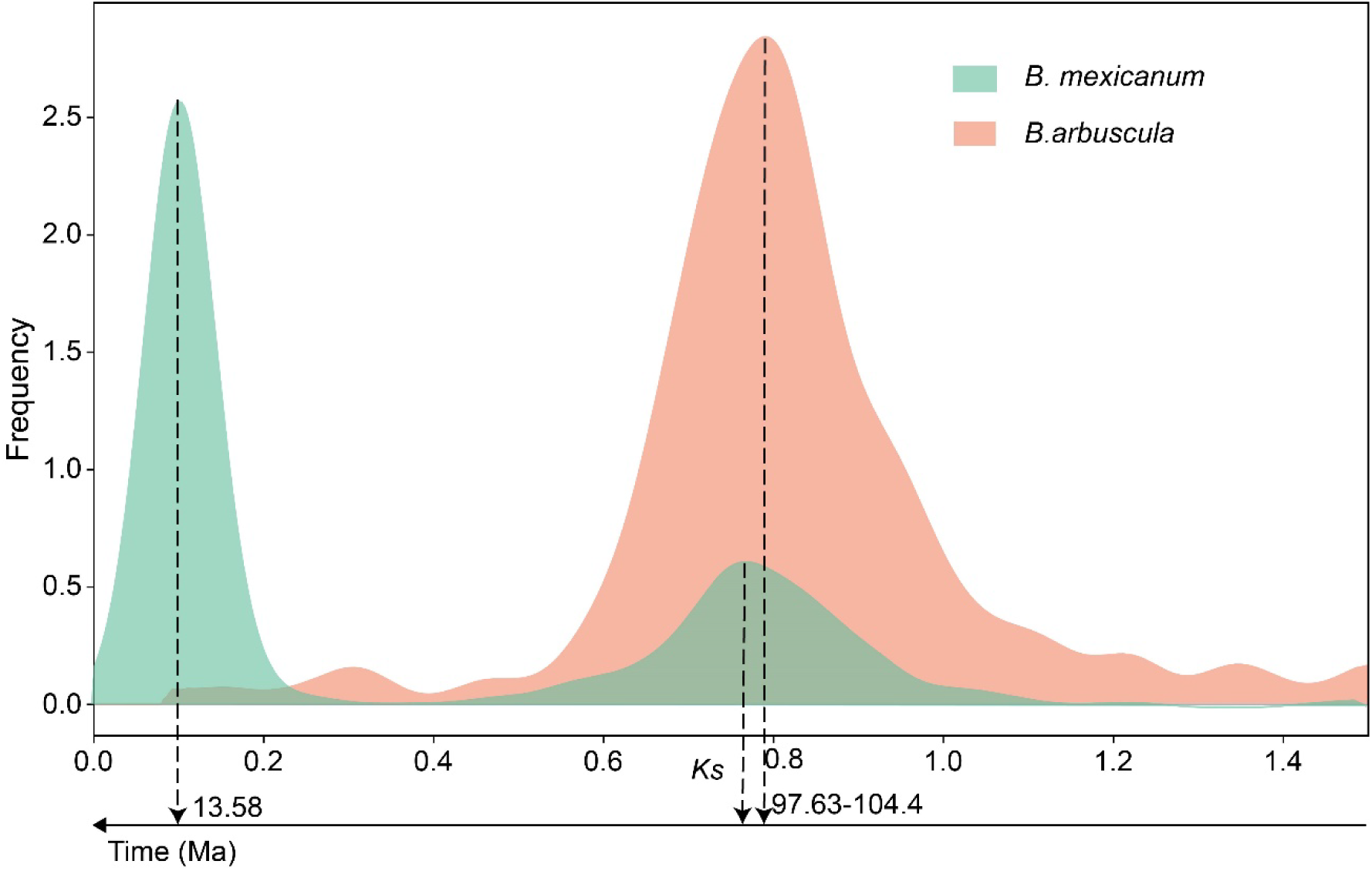
Distribution of pairwise synonymous substitution distances (*Ks*) between paralogous genes of the *Brachypodium arbuscula* genome and the two *B. mexicanum* subgenomes (BmexP, BmexU), and their corresponding time ages. Pairwise calculations within each genome or subgenome provide *Ks* values of ∼0.8 and inferred ages of 97.63-104.4 Ma, which agree with those of the ancient proto-grass polyploidization, while pairwise calculations between BmexP and BmexU provide a value of ∼0.1 and an inferred age of ∼13.58 Ma, which corresponds to the divergence time of its two subgenomes (parental lineages).

**Supplementary Fig. 4.**
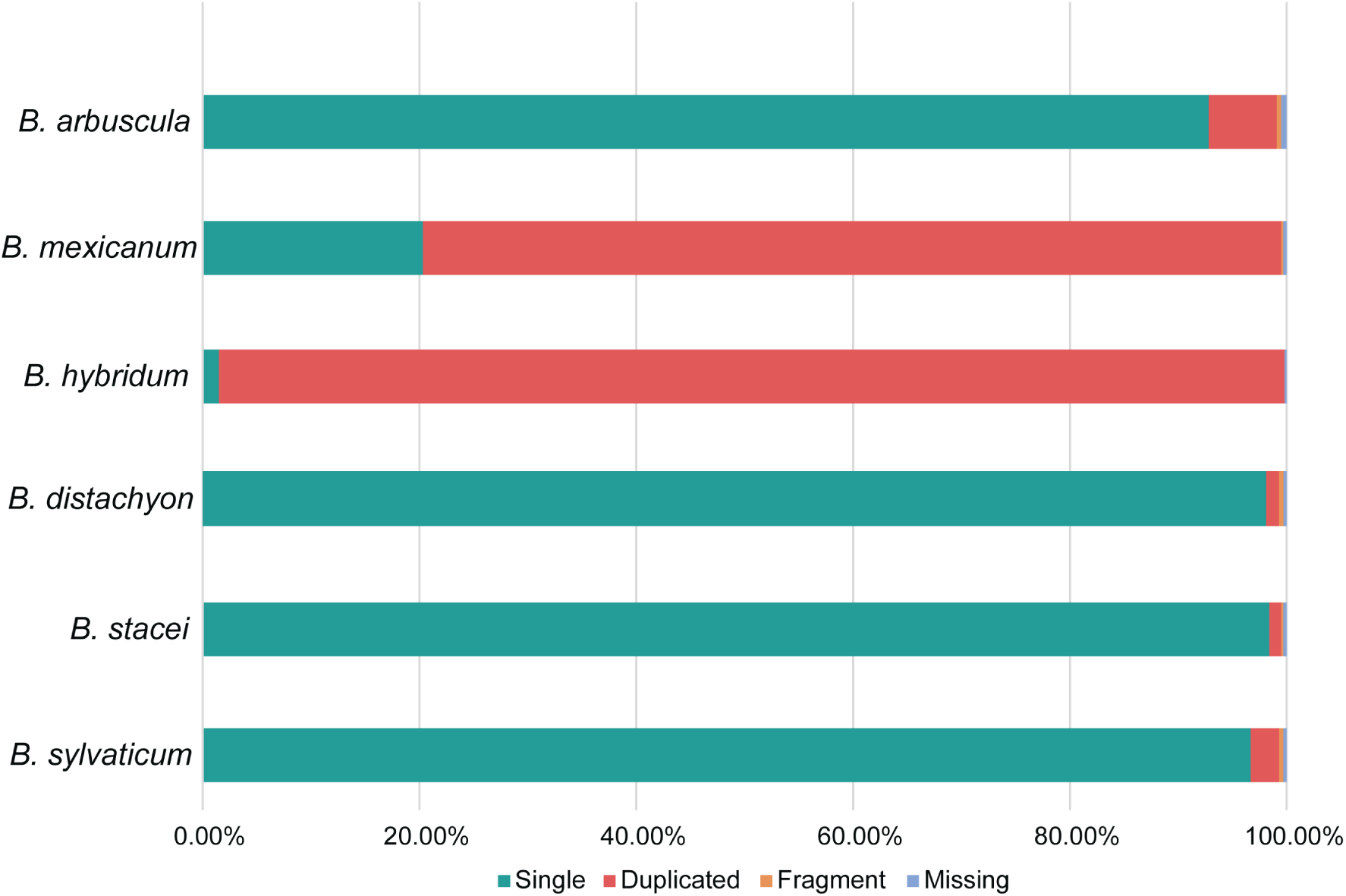
Results of BUSCO assessment of gene annotations for the genomes of the six *Brachypodium* species. Note the predominance of single-copy genes in diploid genomes and duplicated genes in allotetraploid genomes.

**Supplementary Fig. 5.**
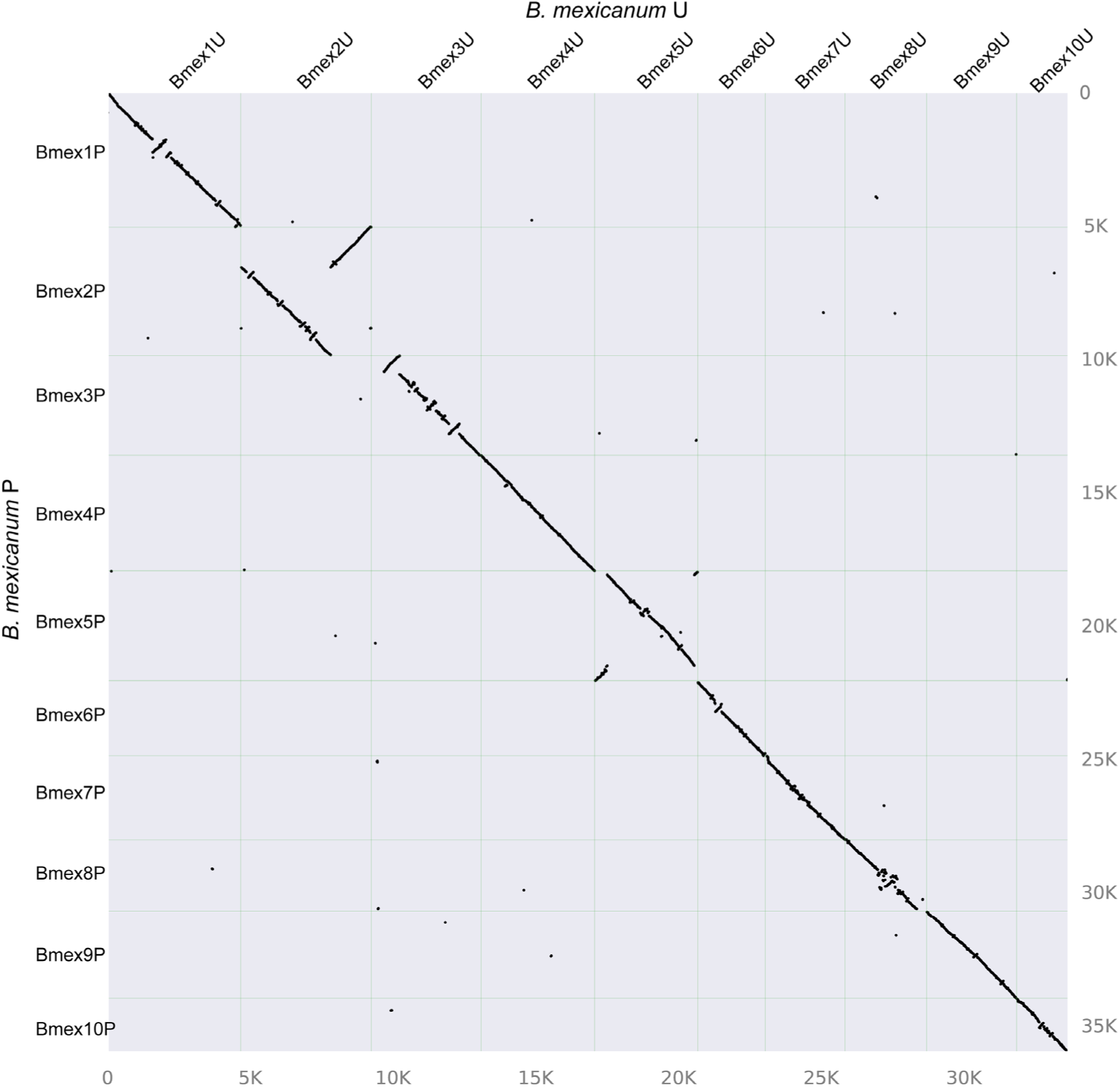
Dotplot of the chromosomes between *B. mexicanum* subgenome P and subgenome U.

**Supplementary Fig. 6.**
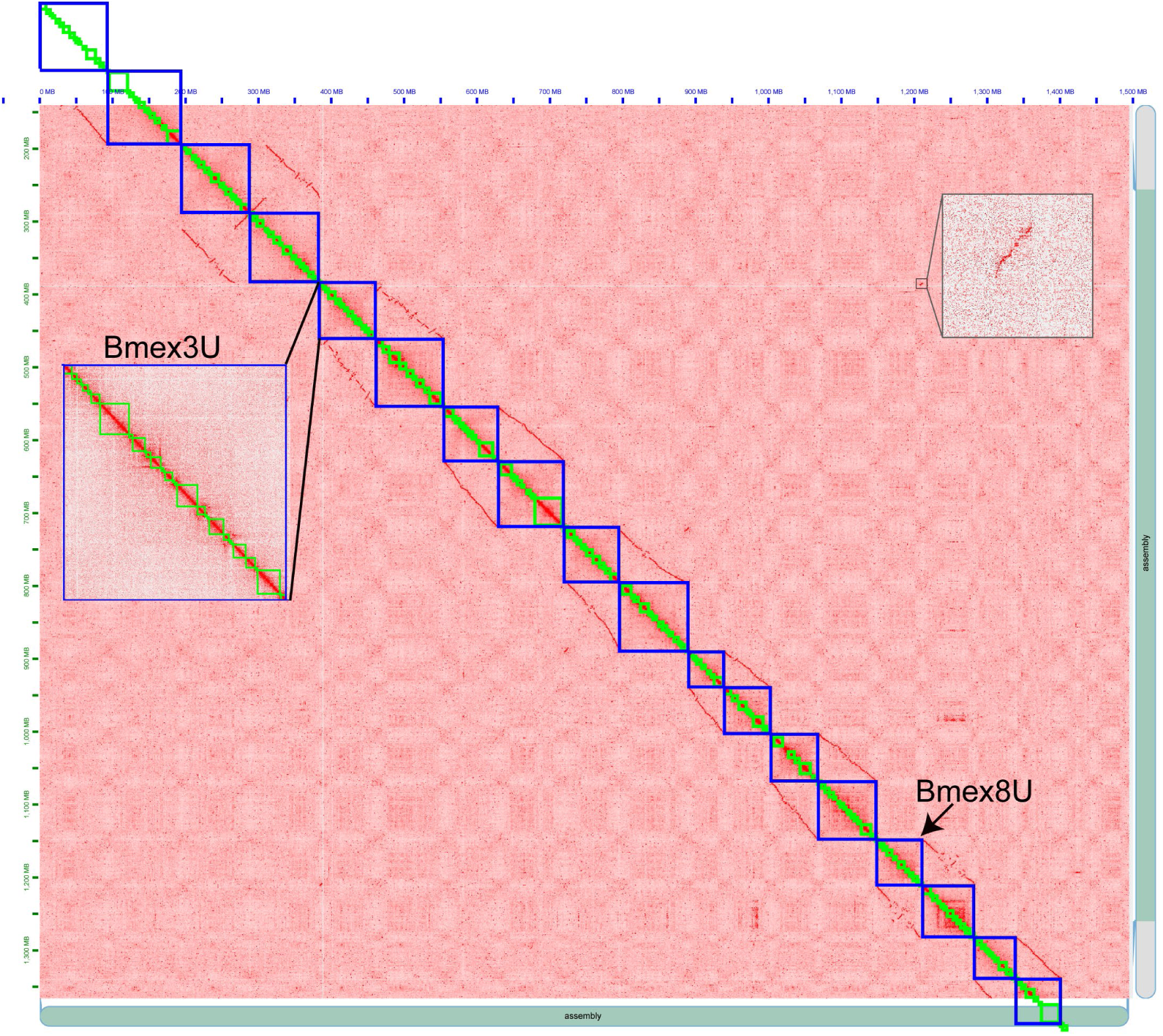
HiC interaction map supporting the translocation of a block from the tail of Bmex8P to Bmex3U.

**Supplementary Fig. 7.**
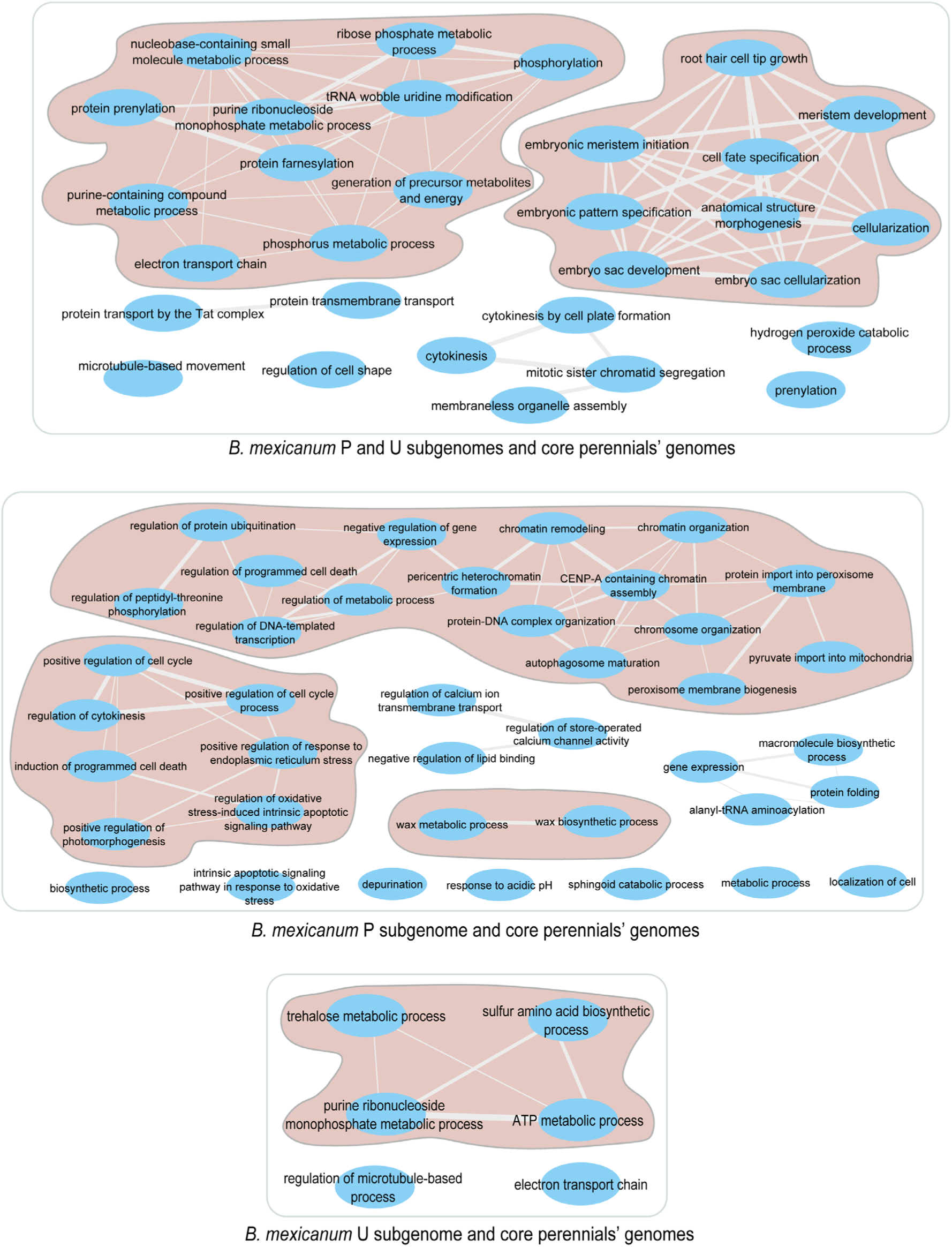
Summarized GO terms that were enriched for orthogroups shared exclusively by the *Brachypodium mexicanum* P and U subgenomes and the core perennial species. Shaded terms indicate those related to perennial habit.

**Supplementary Fig. 8.**
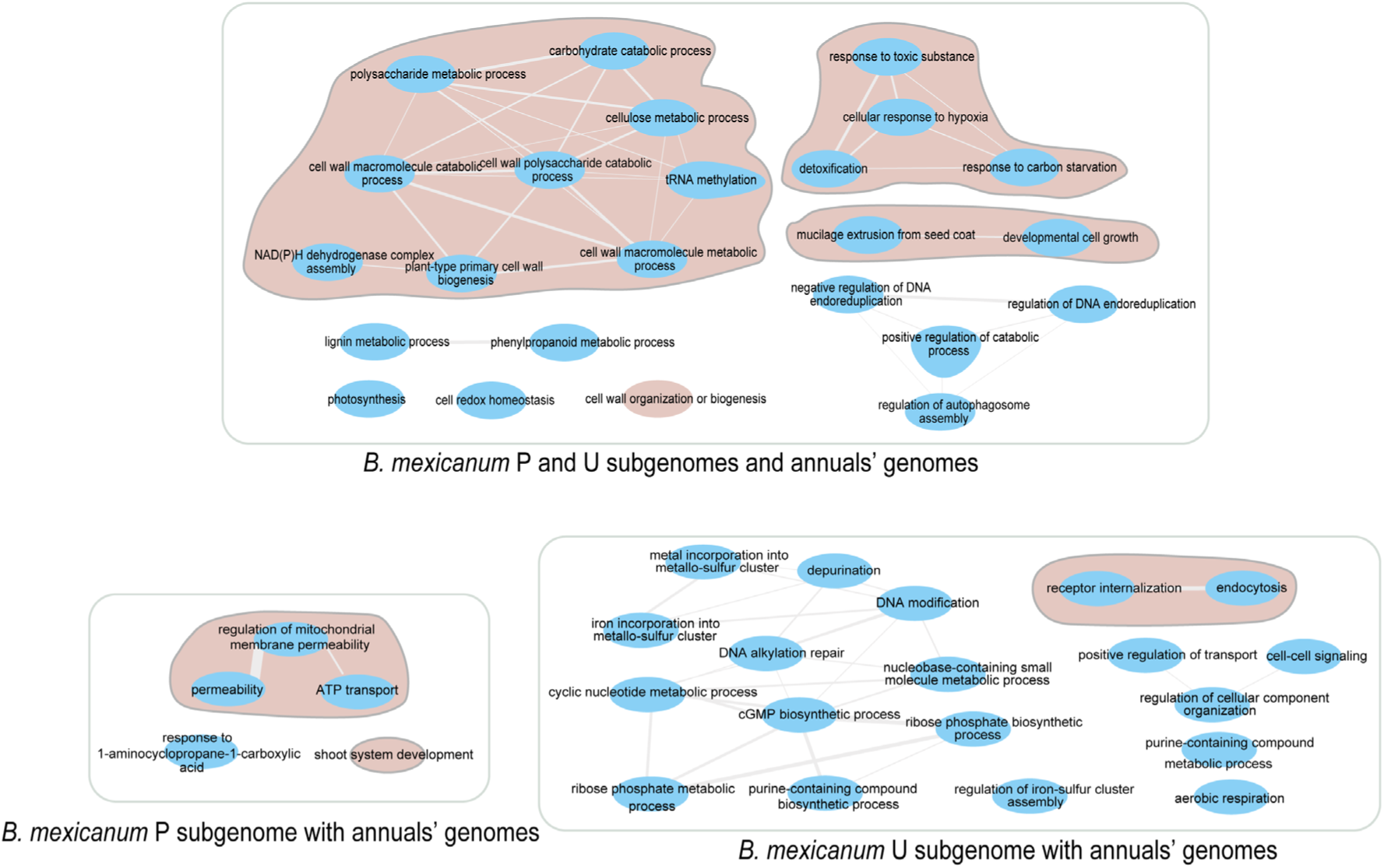
Summarized GO terms that were enriched for orthogroups shared exclusively by the *Brachypodium mexicanum* P and U subgenomes and the annual species. Shaded terms indicate those related to annual habit.

**Supplementary Fig. 9.**
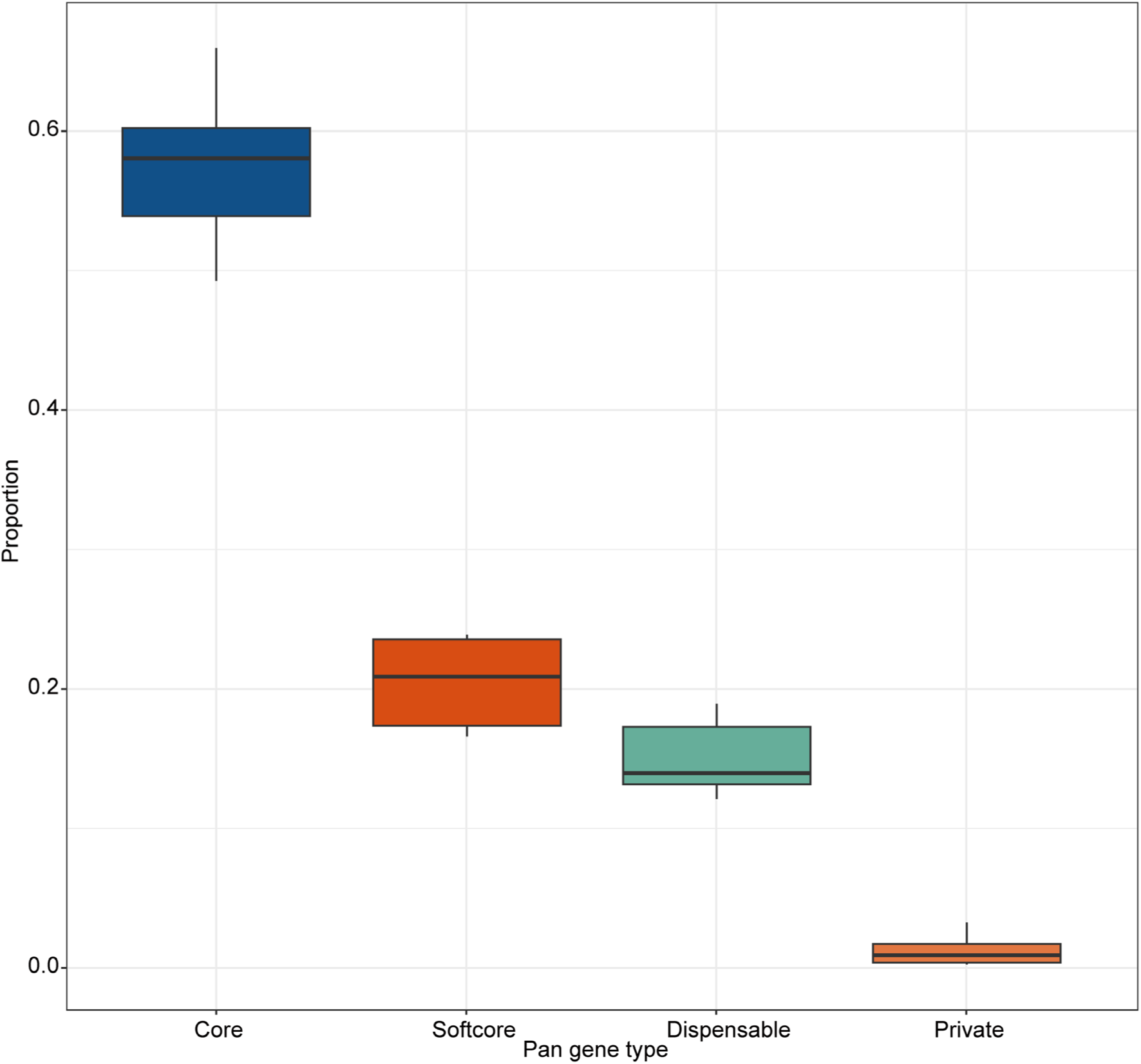
Proportion of each class of pangenes with respect to the total number of genes of the eight *Brachypodium* (sub)genomes under study and separately for each sub(genome) (see also Table S4).

**Supplementary Fig. 10.**
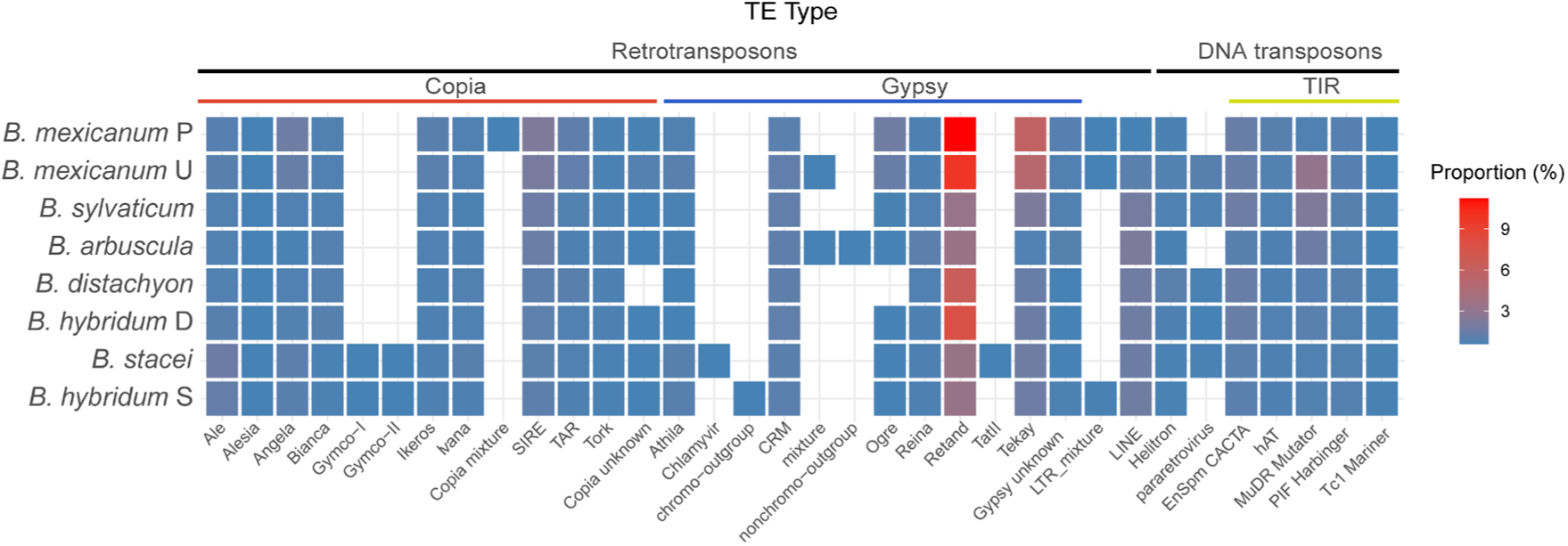
Heatmap showing the length proportion for each type of TE families in the respective sub(genomes) of the six *Brachypodium* species under study.

**Supplementary Fig. 11.**
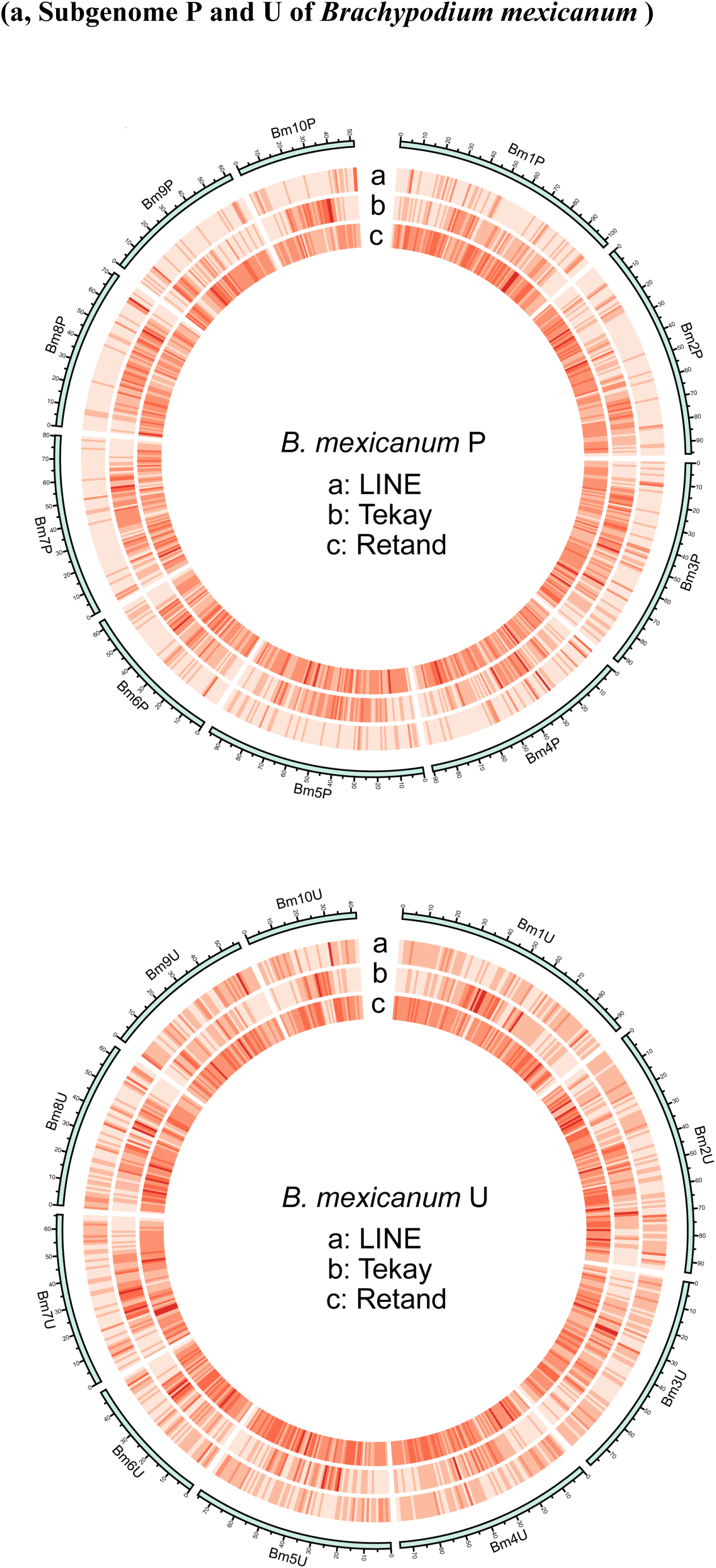
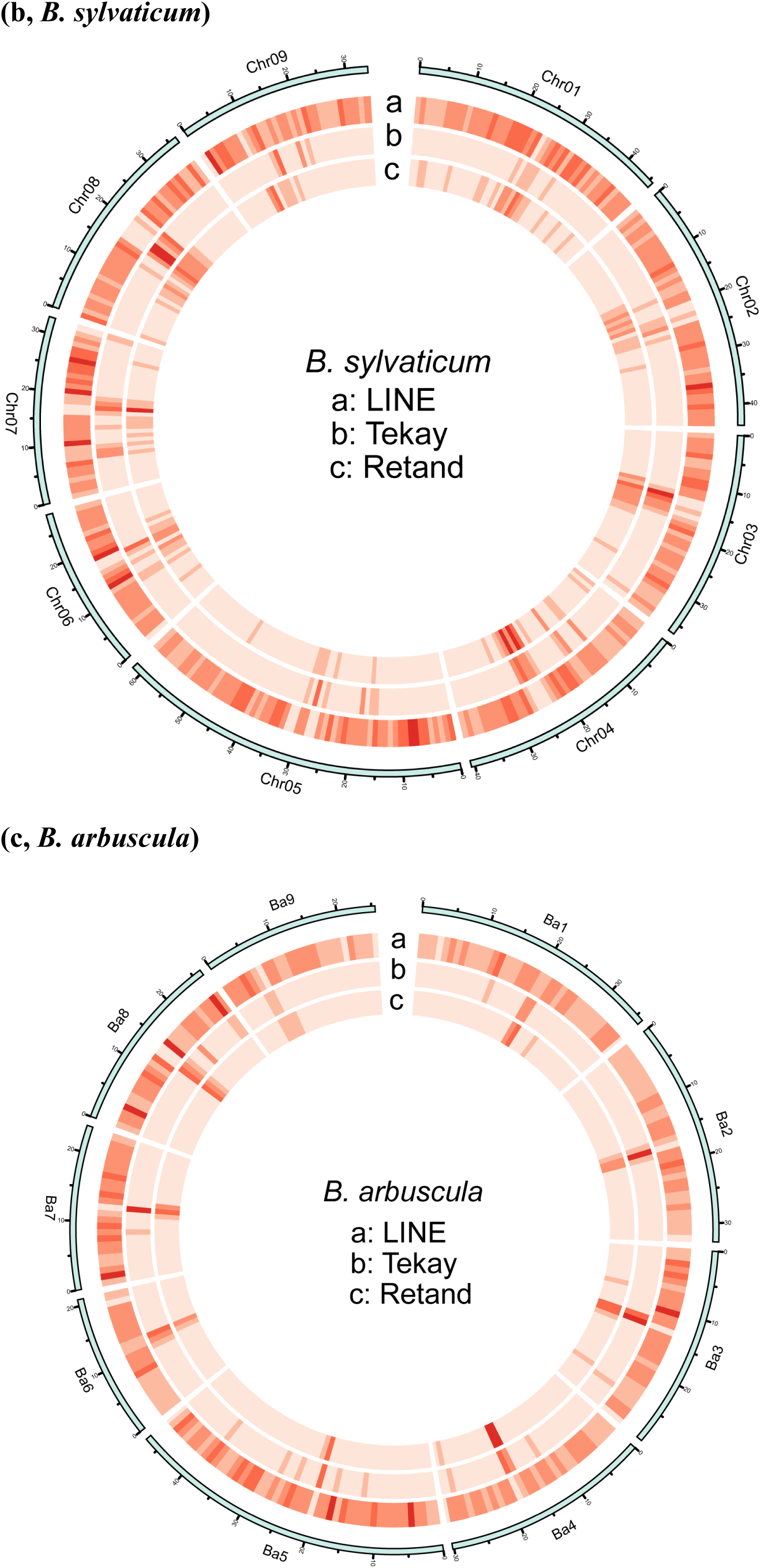
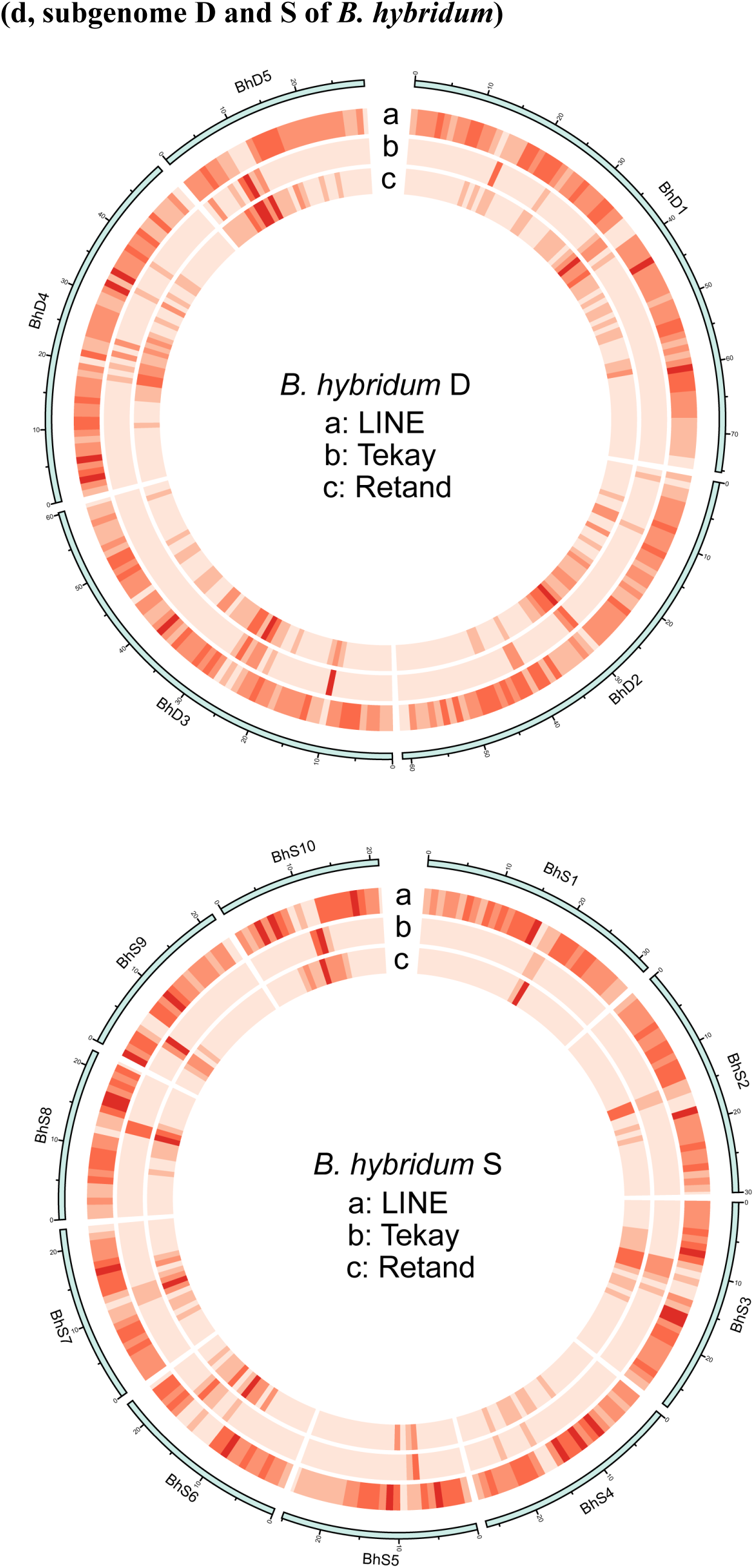
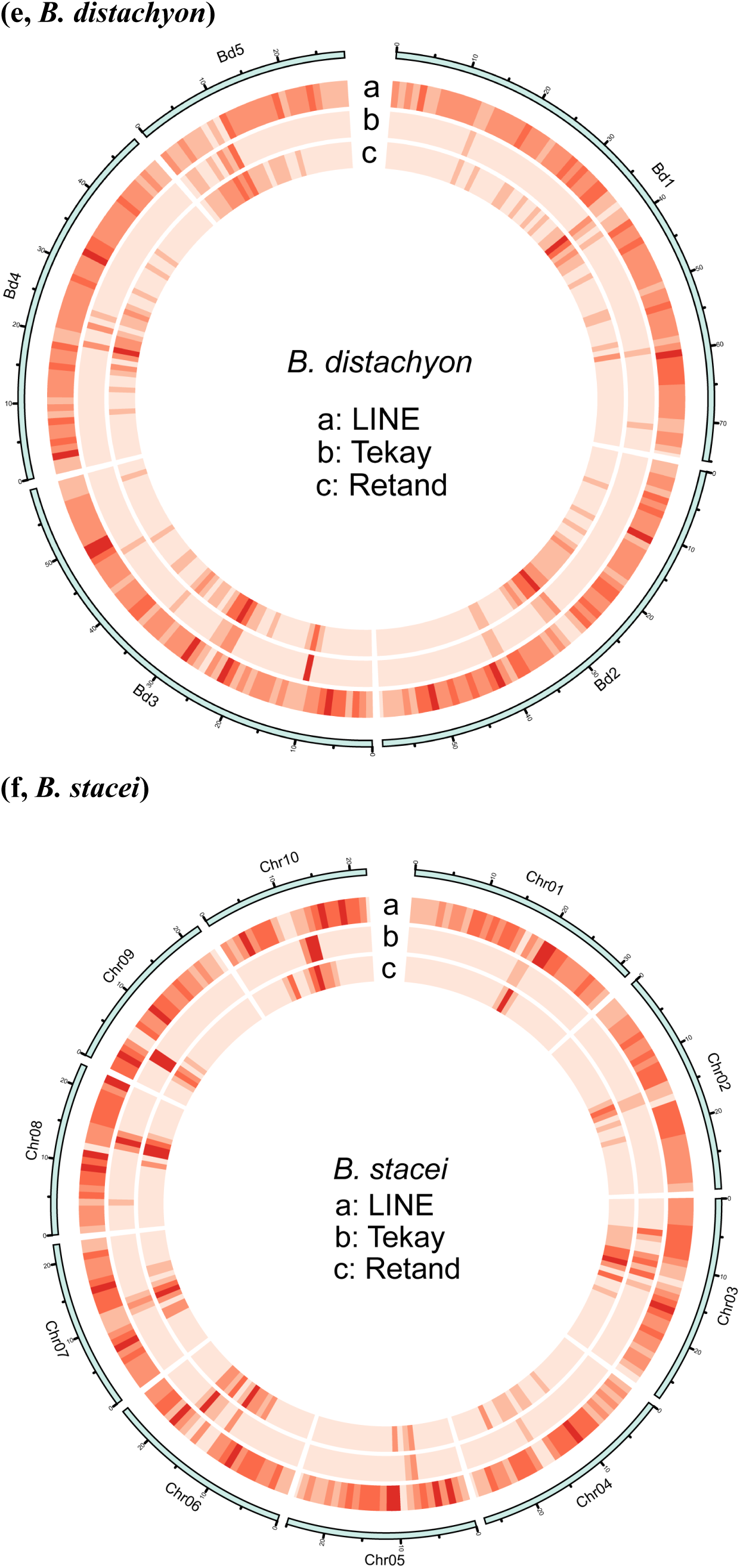
Density of TE copies for the top three subfamilies with the highest copies along the chromosomes of. **(a)** *B. mexicanum* subgenomes P and U, **(b)** *B. sylvaticum* and **(c)** *B. arbuscula,* **(d)** *B. hybridum* subgenomes D and S, **(e)** *B. distachyon* and **(f)** *B. stacei*.

**Supplementary Fig. 12.**
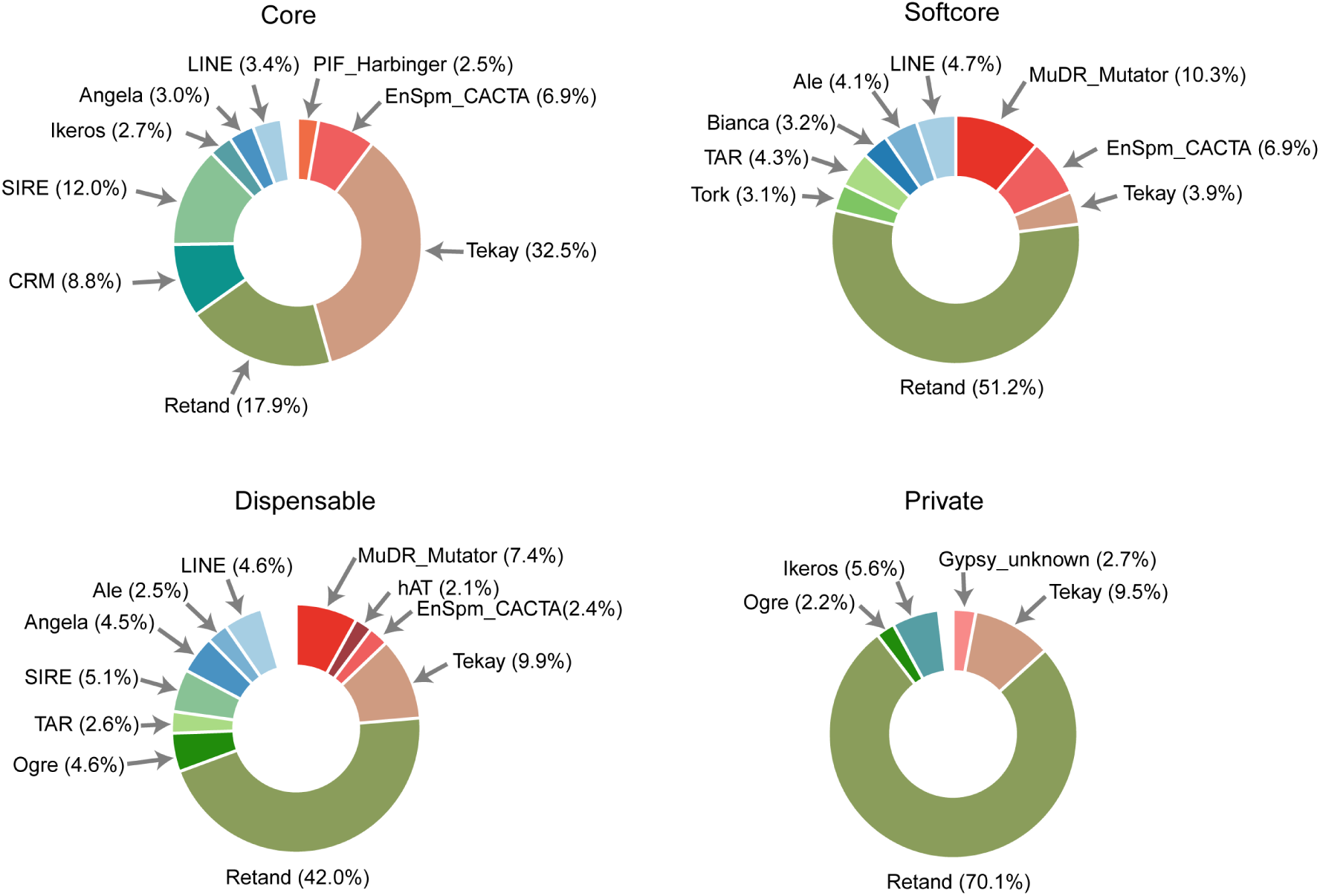
The *Brachypodium* TE family length proportions for each pan-TE type, only those with proportion higher than 2% are shown.

**Supplementary Fig. 13.**
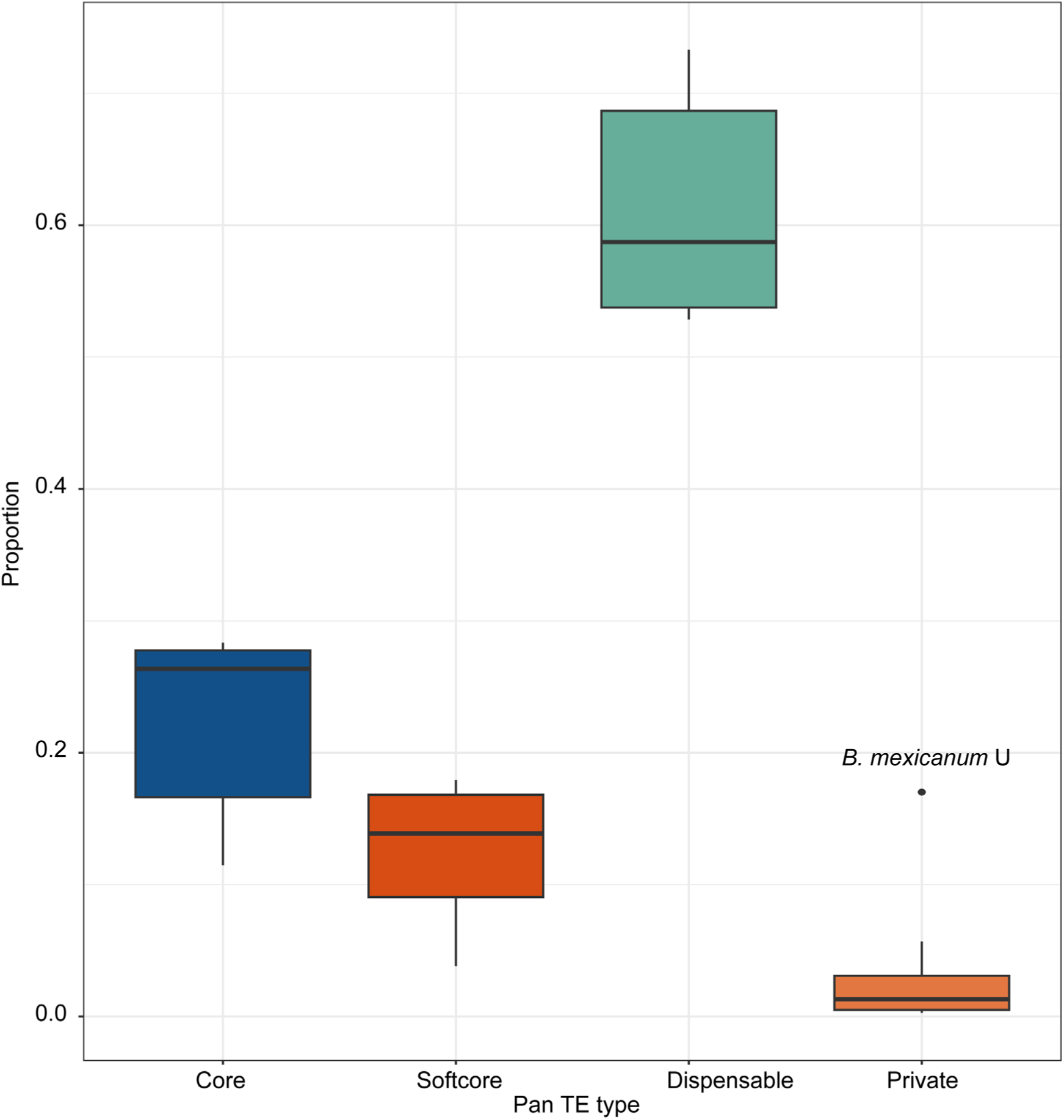
Proportion of each class of pan-TEs with respect to the total number of TEs of the eight *Brachypodium* sub(genomes) under study and separately for each (sub)genome (see also Table S7).

**Supplementary Fig. 14.**
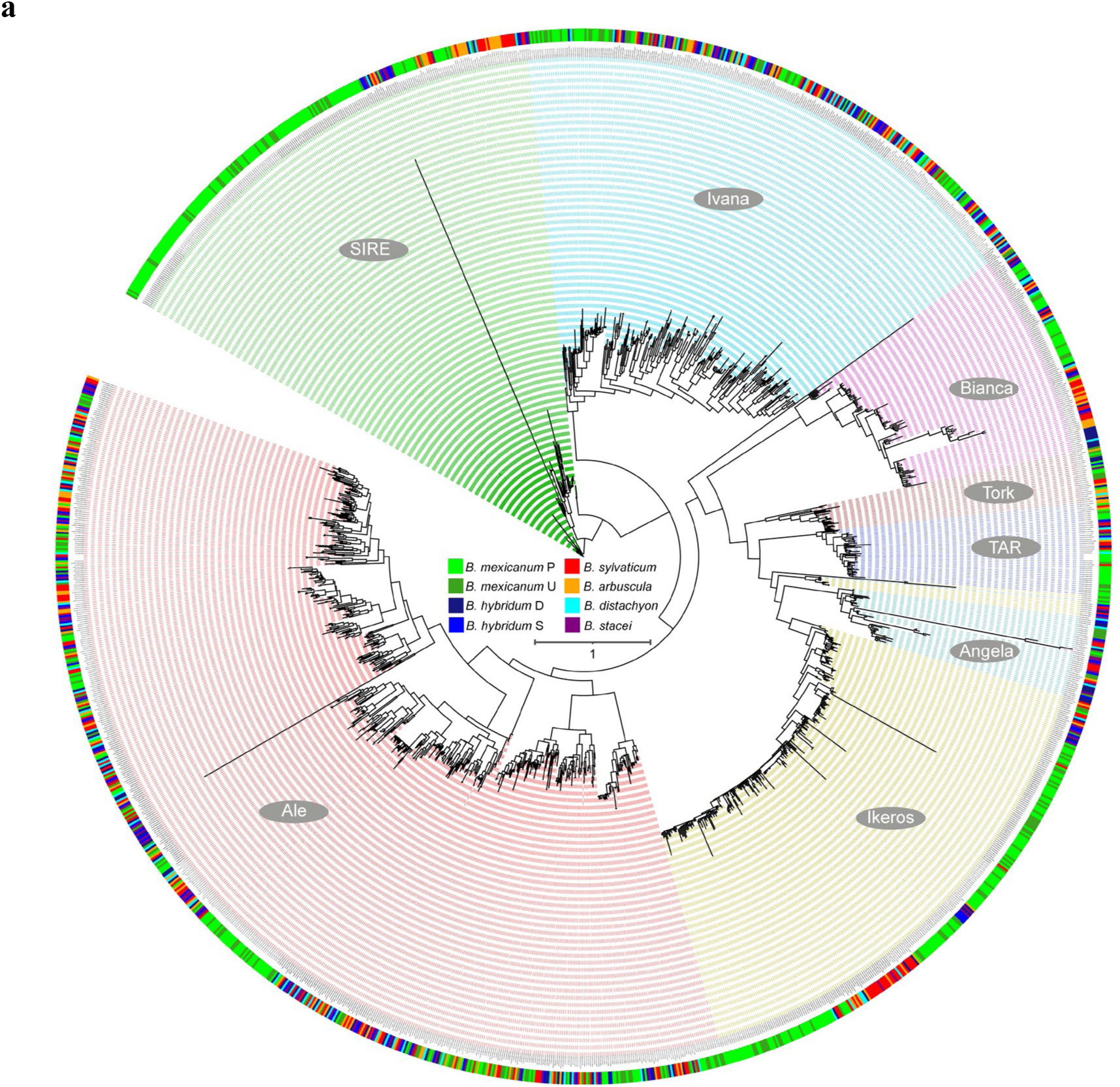
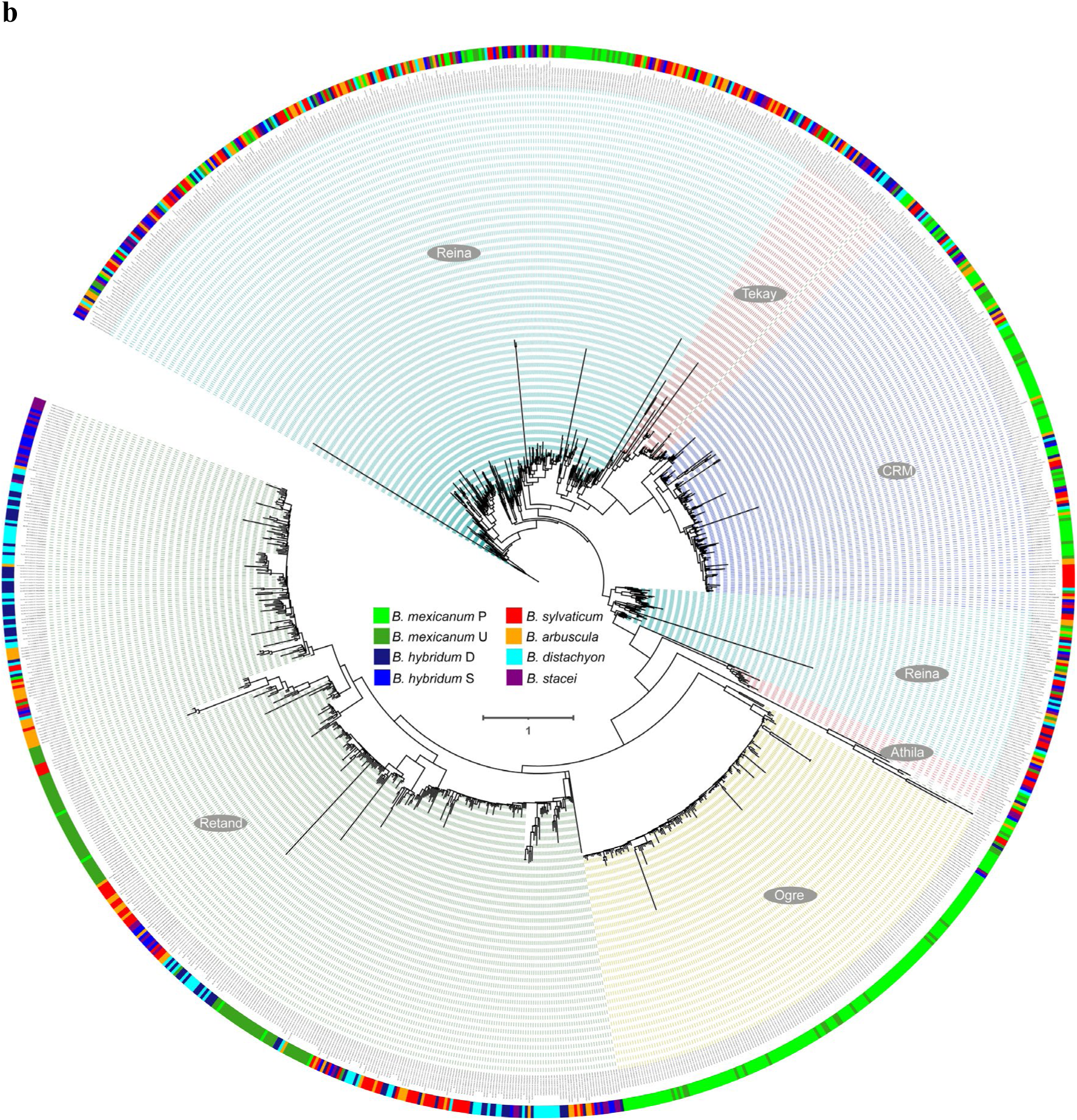
Unrooted phylogenies of (a) *Copia* and (b) *Gypsy* TE families from the eight *Brachypodium* (sub)genomes under study based on RT domains. Color codes for 5 (sub)genomic provenances are indicated in the corresponding charts.

---

**Supplementary Table 1.** Statistics on the assembly and annotation of the *Brachypodium arbuscula* Barb1 and *B. mexicanum* Bmex347 genomes (in bold), and comparison with data from other available *Brachypodium* reference genomes1,2. The BUSCO completeness of the six *Brachypodium* species were assessed using BUSCO v.5.3.23 against the “embryophyta_odb10” database.

|  | *Brachypodium*<br>*arbuscula*<br>**Barb1 (v3.1)** | *Brachypodium*<br>*mexicanum*<br>**Bmex347 (v2.1)** | *Brachypodium*<br>*distachyon*<br>Bd21 (v.3.2) | *Brachypodium*<br>*stacei*<br>BstaABR114 (v.1.1) | *Brachypodium*<br>*hybridum*<br>BhybABR113 (v.1.1) | *Brachypodium*<br>*sylvaticum*<br>Bsyl_Ain1 (v.1.1) |
| --- | --- | --- | --- | --- | --- | --- |
| Assembled genome size | **283.6MB** | **1.53GB** | 271MB | 234MB | 509MB | 358MB |
| Scaffold N50 | **30.19MB** | **78.41MB** | 59.13MB | 23.06MB | 30.6MB | 38.76MB |
| Percentage Single copy BUSCOS | **92.80%** | **20.30%** | 98.10% | 98.40% | 1.50% | 96.70% |
| Percentage of duplicated copy BUSCOS | **6.30%** | **79.20%** | 1.20% | 1.10% | 98.30% | 2.60% |
| Percentage of complete BUSCOS | **99.10%** | **99.50%** | 99.30% | 99.50% | 99.80% | 99.30% |
| Percentage of repeat elements | **38.26%** | **71.12%** | 36.51% | 38.25% | 45.45% | 49.60% |
| Number of predicted Protein-coding genes | **33,605** | **70,845** | 32,439 | 29,898 | 70,160 | 36,927 |

**Supplementary Table 2.**
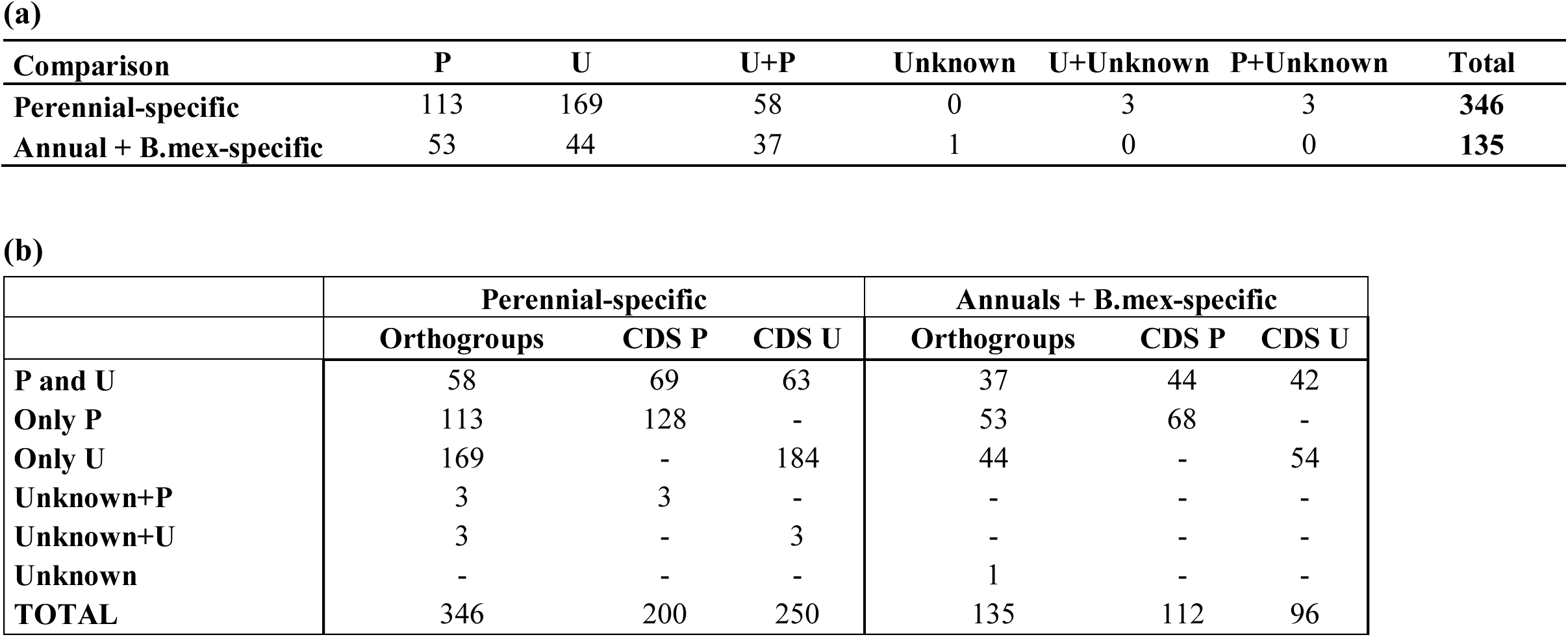
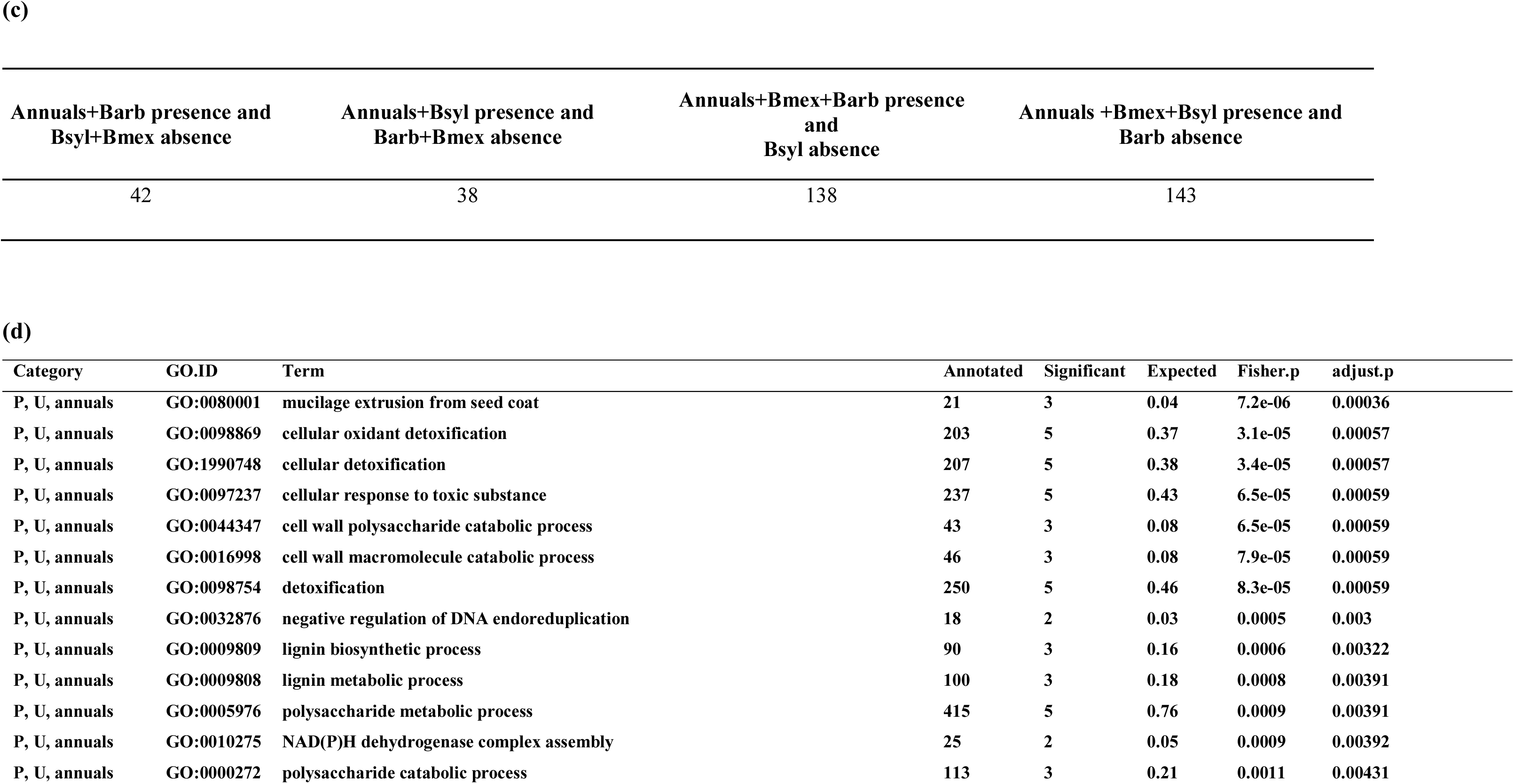
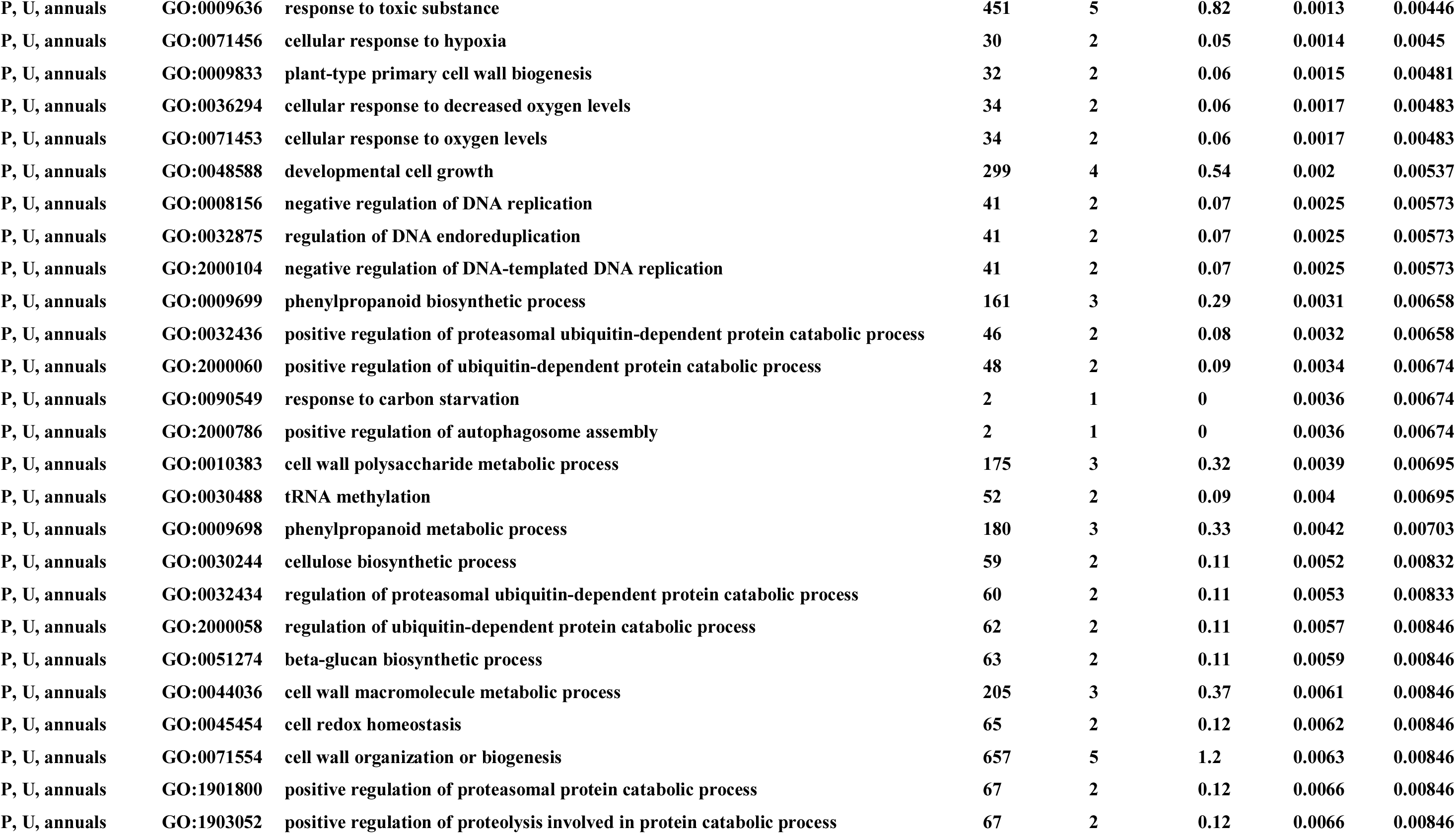
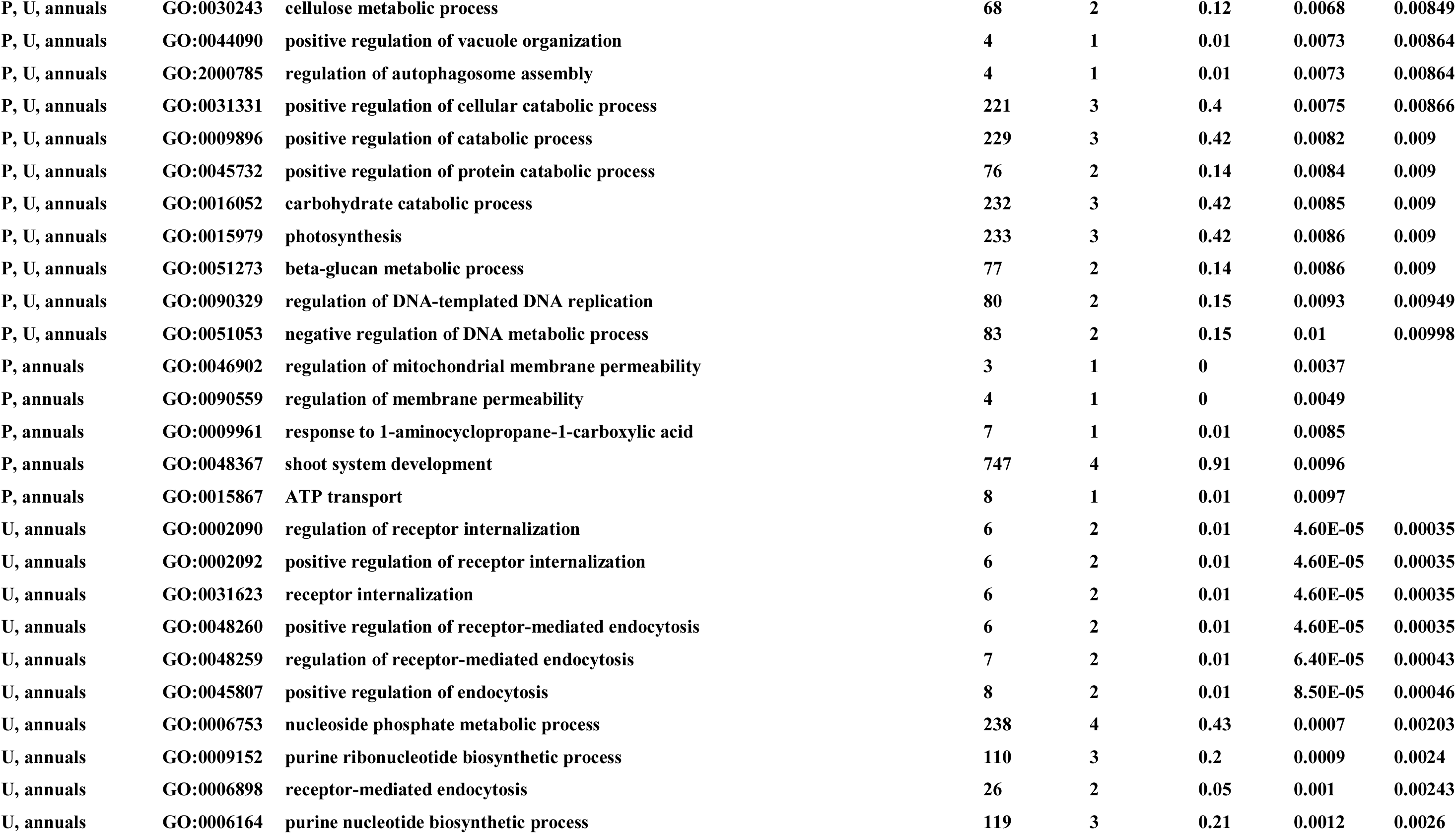
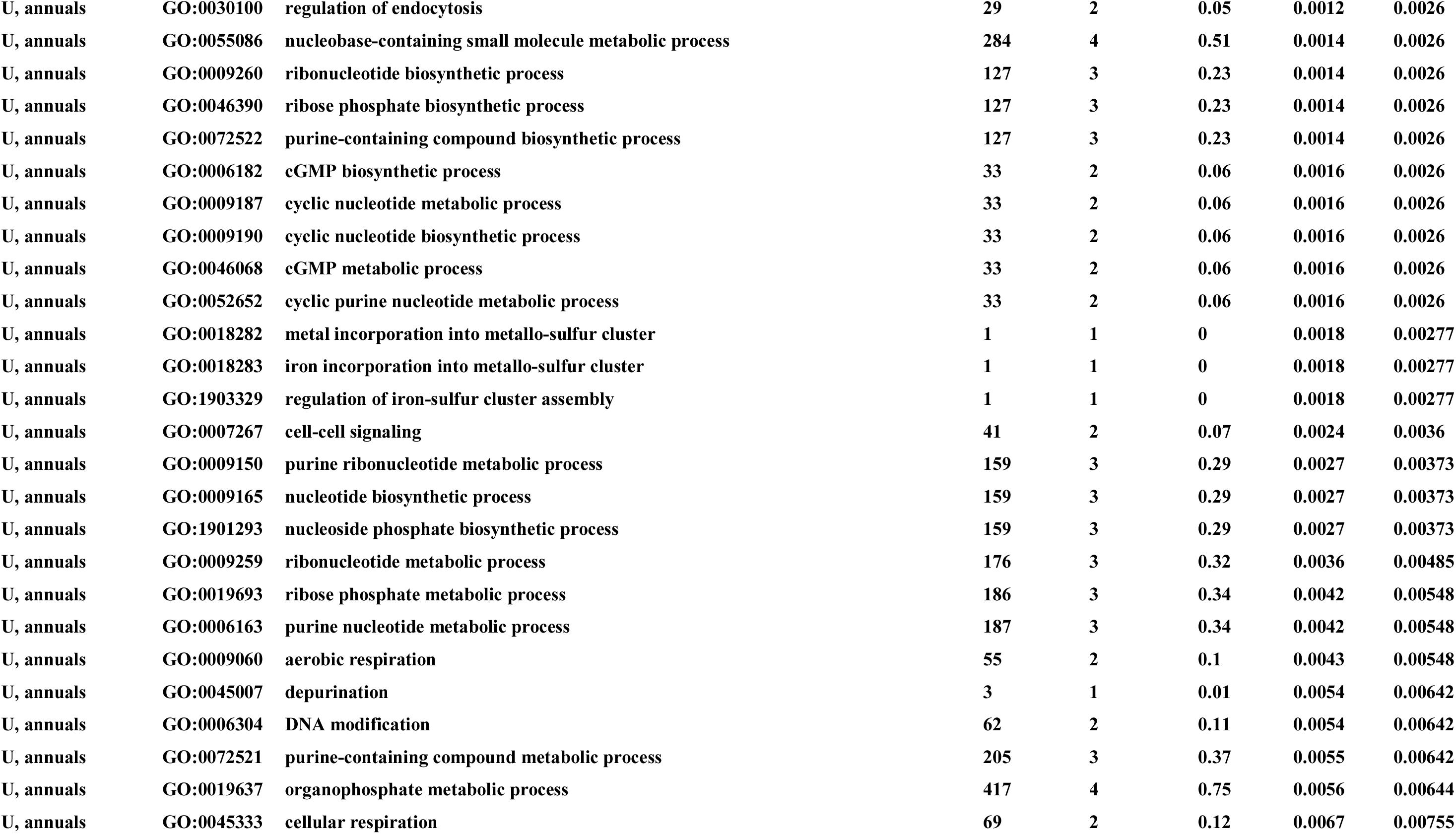
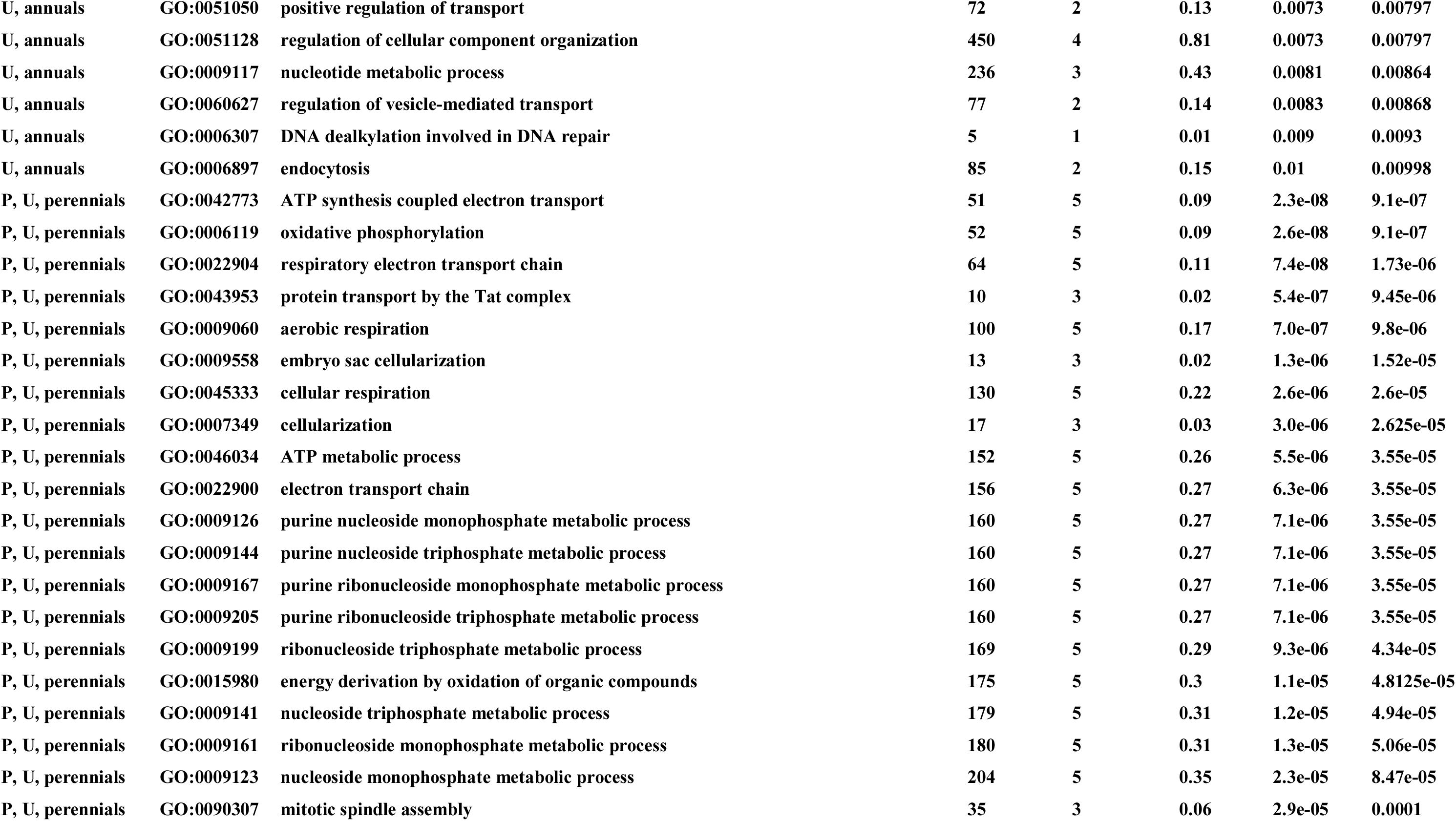
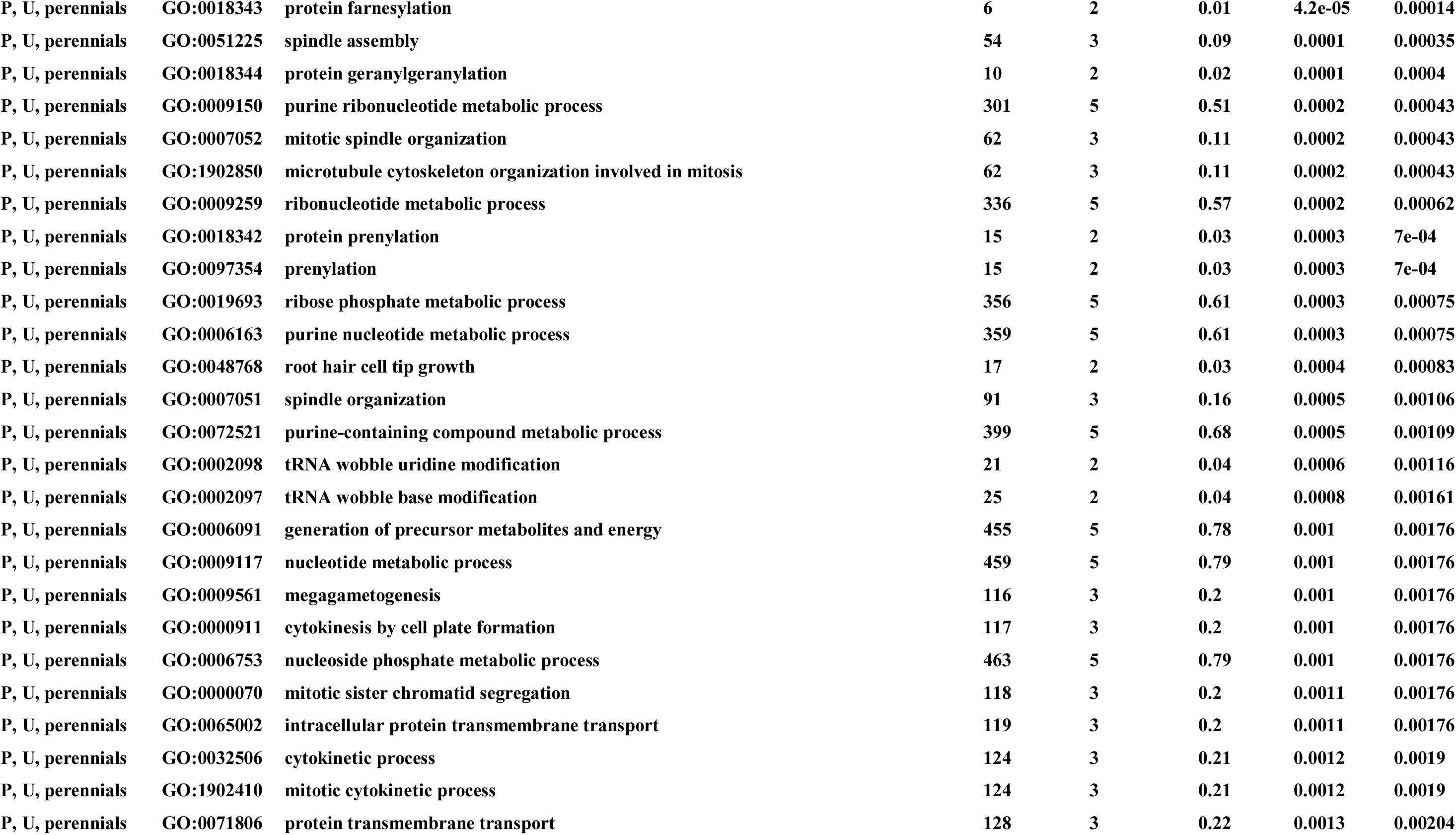
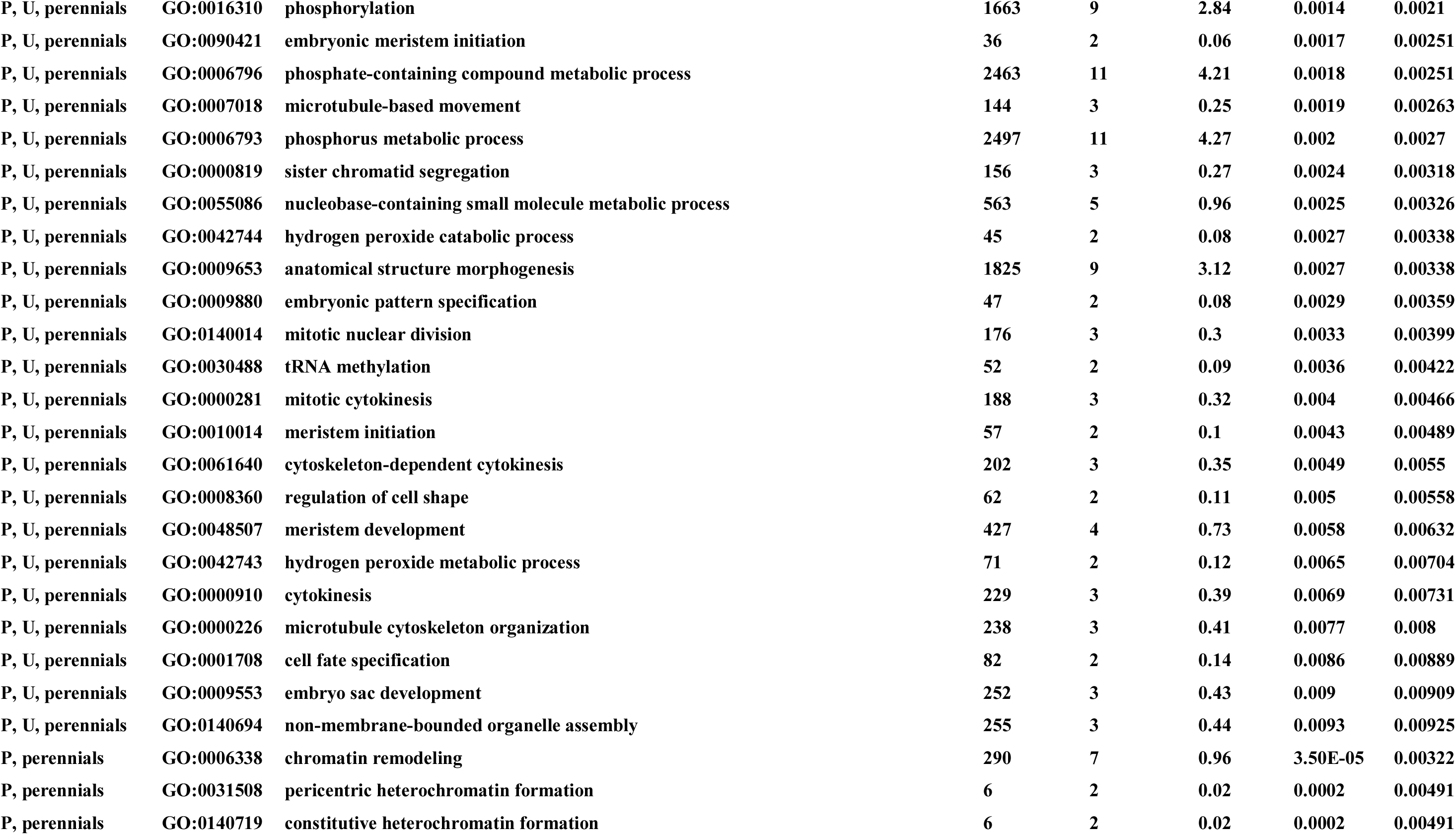
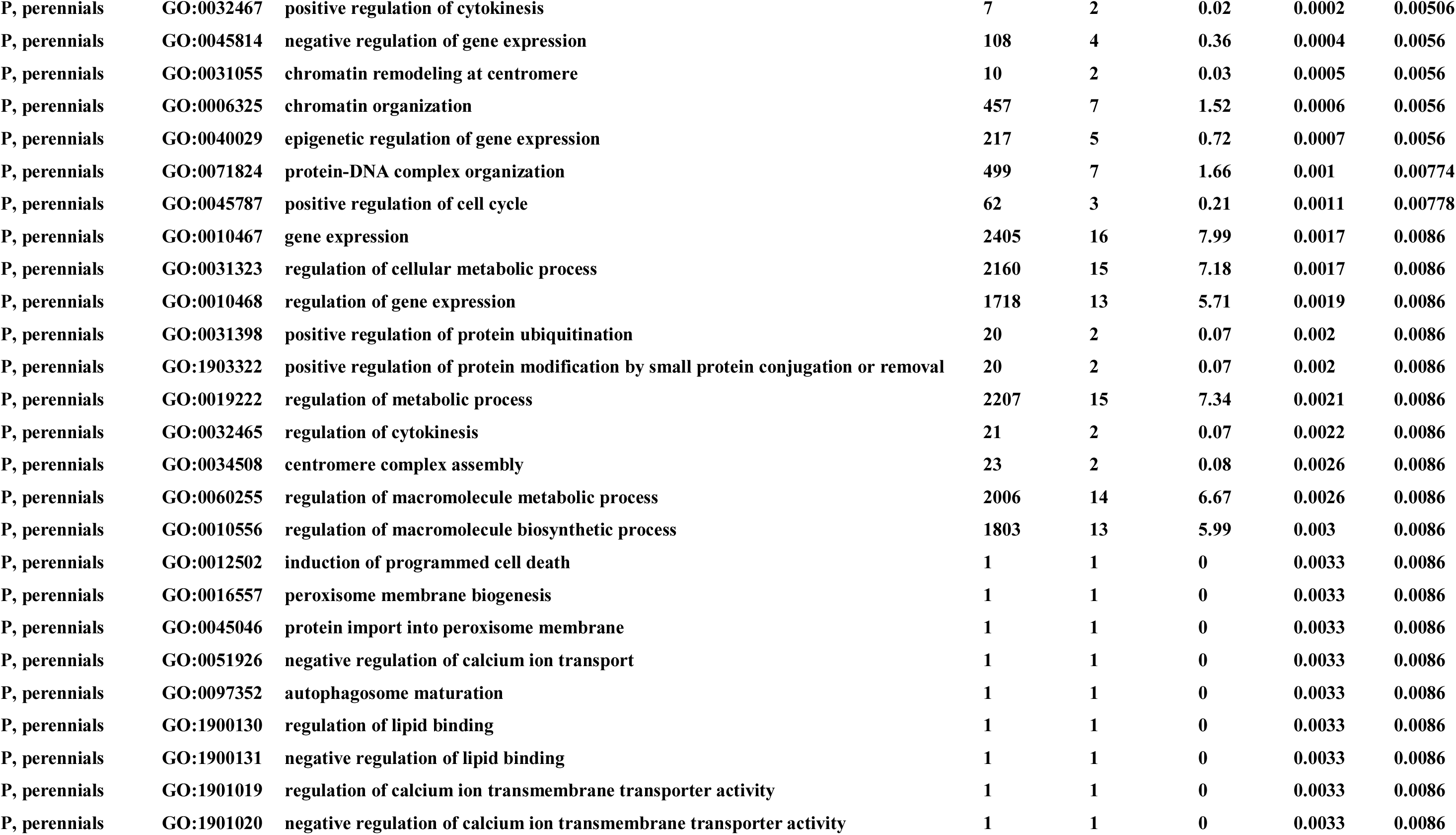
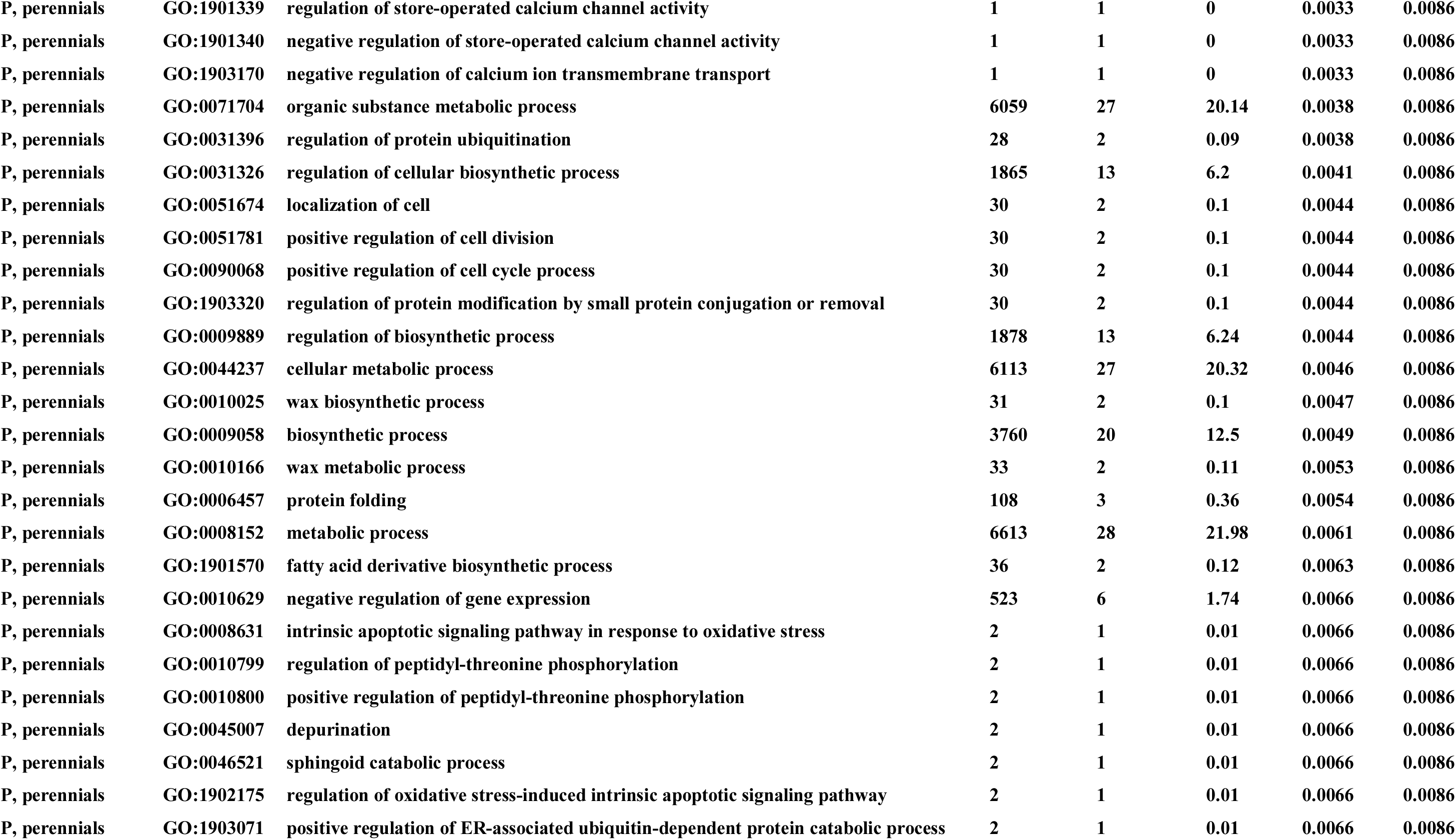
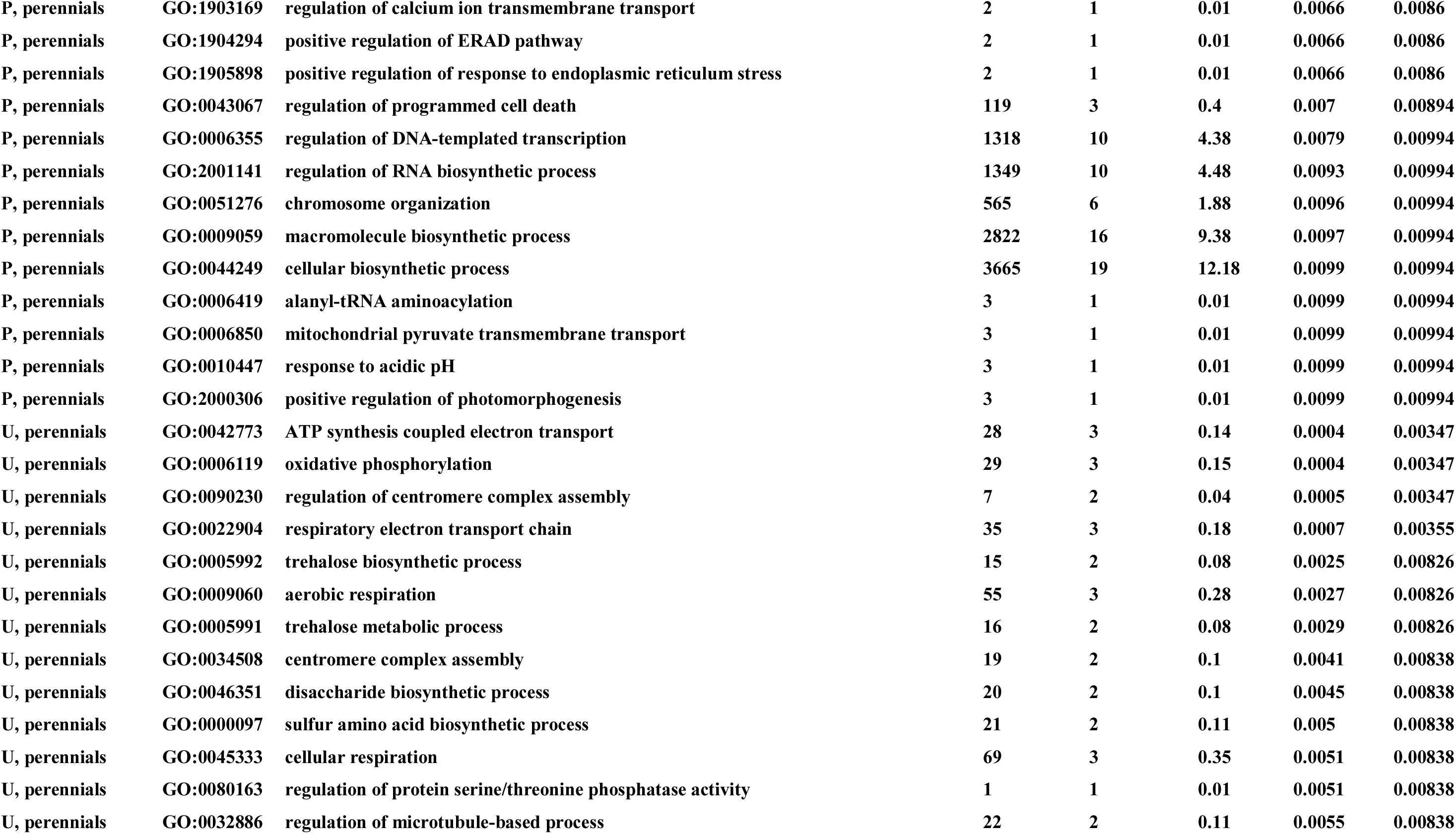
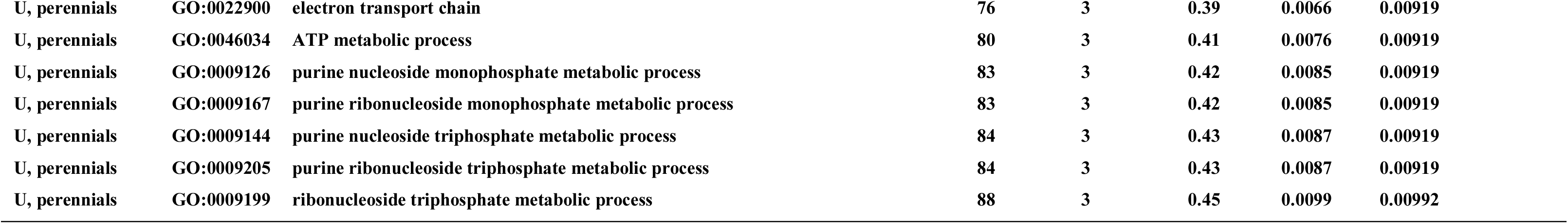
Comparative analyses of CDS-constructed orthogroups to determine the presence/absence of specific genes in perennial or annual *Brachypodium* species and their distribution in the *B. mexicanum* genome. Perennial-specific orthogroups include at least one sequence from all perennial species and none from annual species. Annuals + Bmex-specific orthogroups include at least one sequence from all annual species and from *B. mexicanum*, and none from *B. arbuscula* or *B. sylvaticum*. **(a)** Counts of specific perennial and annual orthogroups in the P and U subgenomes of *B. mexicanum*. **(b)** CDS counts of specific perennial and annual orthogroups in the P and U subgenomes of *B. mexicanum*. **(c)** Multiple comparisons between orthogroups containing all three annual species and orthogroups containing any of the perennial species. (**d**) GO enrichment results of the genes in six groups, including genes shared by the P and U subgenomes of *B. mexicanum* and annual species, by the P subgenome only and annual species, by the U subgenome only and annual species, by the P and U subgenomes and perennial species, by the P subgenome only and perennial species, and by the U subgenome only and perennial species.

**(a)**
| Comparison | P | U | U+P | Unknown | U+Unknown | P+Unknown | Total |
| --- | --- | --- | --- | --- | --- | --- | --- |
| Perennial-specific | 113 | 169 | 58 | 0 | 3 | 3 | 346 |
| Annual + B.mex-specific | 53 | 44 | 37 | 1 | 0 | 0 | 135 |

**(b)**
|  | Perennial-specific |  |  | Annuals + B.mex-specific |  |  |
| --- | --- | --- | --- | --- | --- | --- |
|  | Orthogroups | CDS P | CDS U | Orthogroups | CDS P | CDS U |
| P and U | 58 | 69 | 63 | 37 | 44 | 42 |
| Only P | 113 | 128 | - | 53 | 68 | - |
| Only U | 169 | - | 184 | 44 | - | 54 |
| Unknown+P | 3 | 3 | - | - | - | - |
| Unknown+U | 3 | - | 3 | - | - | - |
| Unknown | - | - | - | 1 | - | - |
| TOTAL | 346 | 200 | 250 | 135 | 112 | 96 |

(c)
| **Annuals+Barb presence and**<br>**Bsyl+Bmex absence** | **Annuals+Bsyl presence and**<br>**Barb+Bmex absence** | **Annuals+Bmex+Barb presence**<br>**and**<br>**Bsyl absence** | **Annuals +Bmex+Bsyl presence and**<br>**Barb absence** |
| --- | --- | --- | --- |
| 42 | 38 | 138 | 143 |

(d)
| **Category** | **GO.ID** | **Term** | **Annotated** | **Significant** | **Expected** | **Fisher.p** | **adjust.p** |
| --- | --- | --- | --- | --- | --- | --- | --- |
| P, U, annuals | GO:0080001 | mucilage extrusion from seed coat | 21 | 3 | 0.04 | 7.2e-06 | 0.00036 |
| P, U, annuals | GO:0098869 | cellular oxidant detoxification | 203 | 5 | 0.37 | 3.1e-05 | 0.00057 |
| P, U, annuals | GO:1990748 | cellular detoxification | 207 | 5 | 0.38 | 3.4e-05 | 0.00057 |
| P, U, annuals | GO:0097237 | cellular response to toxic substance | 237 | 5 | 0.43 | 6.5e-05 | 0.00059 |
| P, U, annuals | GO:0044347 | cell wall polysaccharide catabolic process | 43 | 3 | 0.08 | 6.5e-05 | 0.00059 |
| P, U, annuals | GO:0016998 | cell wall macromolecule catabolic process | 46 | 3 | 0.08 | 7.9e-05 | 0.00059 |
| P, U, annuals | GO:0098754 | detoxification | 250 | 5 | 0.46 | 8.3e-05 | 0.00059 |
| P, U, annuals | GO:0032876 | negative regulation of DNA endoreduplication | 18 | 2 | 0.03 | 0.0005 | 0.003 |
| P, U, annuals | GO:0009809 | lignin biosynthetic process | 90 | 3 | 0.16 | 0.0006 | 0.00322 |
| P, U, annuals | GO:0009808 | lignin metabolic process | 100 | 3 | 0.18 | 0.0008 | 0.00391 |
| P, U, annuals | GO:0005976 | polysaccharide metabolic process | 415 | 5 | 0.76 | 0.0009 | 0.00391 |
| P, U, annuals | GO:0010275 | NAD(P)H dehydrogenase complex assembly | 25 | 2 | 0.05 | 0.0009 | 0.00392 |
| P, U, annuals | GO:0000272 | polysaccharide catabolic process | 113 | 3 | 0.21 | 0.0011 | 0.00431 |
| P, U, annuals | GO:0009636 | response to toxic substance | 451 | 5 | 0.82 | 0.0013 | 0.00446 |
| P, U, annuals | GO:0071456 | cellular response to hypoxia | 30 | 2 | 0.05 | 0.0014 | 0.0045 |
| P, U, annuals | GO:0009833 | plant-type primary cell wall biogenesis | 32 | 2 | 0.06 | 0.0015 | 0.00481 |
| P, U, annuals | GO:0036294 | cellular response to decreased oxygen levels | 34 | 2 | 0.06 | 0.0017 | 0.00483 |
| P, U, annuals | GO:0071453 | cellular response to oxygen levels | 34 | 2 | 0.06 | 0.0017 | 0.00483 |
| P, U, annuals | GO:0048588 | developmental cell growth | 299 | 4 | 0.54 | 0.002 | 0.00537 |
| P, U, annuals | GO:0008156 | negative regulation of DNA replication | 41 | 2 | 0.07 | 0.0025 | 0.00573 |
| P, U, annuals | GO:0032875 | regulation of DNA endoreduplication | 41 | 2 | 0.07 | 0.0025 | 0.00573 |
| P, U, annuals | GO:2000104 | negative regulation of DNA-templated DNA replication | 41 | 2 | 0.07 | 0.0025 | 0.00573 |
| P, U, annuals | GO:0009699 | phenylpropanoid biosynthetic process | 161 | 3 | 0.29 | 0.0031 | 0.00658 |
| P, U, annuals | GO:0032436 | positive regulation of proteasomal ubiquitin-dependent protein catabolic process | 46 | 2 | 0.08 | 0.0032 | 0.00658 |
| P, U, annuals | GO:2000060 | positive regulation of ubiquitin-dependent protein catabolic process | 48 | 2 | 0.09 | 0.0034 | 0.00674 |
| P, U, annuals | GO:0090549 | response to carbon starvation | 2 | 1 | 0 | 0.0036 | 0.00674 |
| P, U, annuals | GO:2000786 | positive regulation of autophagosome assembly | 2 | 1 | 0 | 0.0036 | 0.00674 |
| P, U, annuals | GO:0010383 | cell wall polysaccharide metabolic process | 175 | 3 | 0.32 | 0.0039 | 0.00695 |
| P, U, annuals | GO:0030488 | tRNA methylation | 52 | 2 | 0.09 | 0.004 | 0.00695 |
| P, U, annuals | GO:0009698 | phenylpropanoid metabolic process | 180 | 3 | 0.33 | 0.0042 | 0.00703 |
| P, U, annuals | GO:0030244 | cellulose biosynthetic process | 59 | 2 | 0.11 | 0.0052 | 0.00832 |
| P, U, annuals | GO:0032434 | regulation of proteasomal ubiquitin-dependent protein catabolic process | 60 | 2 | 0.11 | 0.0053 | 0.00833 |
| P, U, annuals | GO:2000058 | regulation of ubiquitin-dependent protein catabolic process | 62 | 2 | 0.11 | 0.0057 | 0.00846 |
| P, U, annuals | GO:0051274 | beta-glucan biosynthetic process | 63 | 2 | 0.11 | 0.0059 | 0.00846 |
| P, U, annuals | GO:0044036 | cell wall macromolecule metabolic process | 205 | 3 | 0.37 | 0.0061 | 0.00846 |
| P, U, annuals | GO:0045454 | cell redox homeostasis | 65 | 2 | 0.12 | 0.0062 | 0.00846 |
| P, U, annuals | GO:0071554 | cell wall organization or biogenesis | 657 | 5 | 1.2 | 0.0063 | 0.00846 |
| P, U, annuals | GO:1901800 | positive regulation of proteasomal protein catabolic process | 67 | 2 | 0.12 | 0.0066 | 0.00846 |
| P, U, annuals | GO:1903052 | positive regulation of proteolysis involved in protein catabolic process | 67 | 2 | 0.12 | 0.0066 | 0.00846 |
| P, U, annuals | GO:0030243 | cellulose metabolic process | 68 | 2 | 0.12 | 0.0068 | 0.00849 |
| P, U, annuals | GO:0044090 | positive regulation of vacuole organization | 4 | 1 | 0.01 | 0.0073 | 0.00864 |
| P, U, annuals | GO:2000785 | regulation of autophagosome assembly | 4 | 1 | 0.01 | 0.0073 | 0.00864 |
| P, U, annuals | GO:0031331 | positive regulation of cellular catabolic process | 221 | 3 | 0.4 | 0.0075 | 0.00866 |
| P, U, annuals | GO:0009896 | positive regulation of catabolic process | 229 | 3 | 0.42 | 0.0082 | 0.009 |
| P, U, annuals | GO:0045732 | positive regulation of protein catabolic process | 76 | 2 | 0.14 | 0.0084 | 0.009 |
| P, U, annuals | GO:0016052 | carbohydrate catabolic process | 232 | 3 | 0.42 | 0.0085 | 0.009 |
| P, U, annuals | GO:0015979 | photosynthesis | 233 | 3 | 0.42 | 0.0086 | 0.009 |
| P, U, annuals | GO:0051273 | beta-glucan metabolic process | 77 | 2 | 0.14 | 0.0086 | 0.009 |
| P, U, annuals | GO:0090329 | regulation of DNA-templated DNA replication | 80 | 2 | 0.15 | 0.0093 | 0.00949 |
| P, U, annuals | GO:0051053 | negative regulation of DNA metabolic process | 83 | 2 | 0.15 | 0.01 | 0.00998 |
| P, annuals | GO:0046902 | regulation of mitochondrial membrane permeability | 3 | 1 | 0 | 0.0037 |  |
| P, annuals | GO:0090559 | regulation of membrane permeability | 4 | 1 | 0 | 0.0049 |  |
| P, annuals | GO:0009961 | response to 1-aminocyclopropane-1-carboxylic acid | 7 | 1 | 0.01 | 0.0085 |  |
| P, annuals | GO:0048367 | shoot system development | 747 | 4 | 0.91 | 0.0096 |  |
| P, annuals | GO:0015867 | ATP transport | 8 | 1 | 0.01 | 0.0097 |  |
| U, annuals | GO:0002090 | regulation of receptor internalization | 6 | 2 | 0.01 | 4.60E-05 | 0.00035 |
| U, annuals | GO:0002092 | positive regulation of receptor internalization | 6 | 2 | 0.01 | 4.60E-05 | 0.00035 |
| U, annuals | GO:0031623 | receptor internalization | 6 | 2 | 0.01 | 4.60E-05 | 0.00035 |
| U, annuals | GO:0048260 | positive regulation of receptor-mediated endocytosis | 6 | 2 | 0.01 | 4.60E-05 | 0.00035 |
| U, annuals | GO:0048259 | regulation of receptor-mediated endocytosis | 7 | 2 | 0.01 | 6.40E-05 | 0.00043 |
| U, annuals | GO:0045807 | positive regulation of endocytosis | 8 | 2 | 0.01 | 8.50E-05 | 0.00046 |
| U, annuals | GO:0006753 | nucleoside phosphate metabolic process | 238 | 4 | 0.43 | 0.0007 | 0.00203 |
| U, annuals | GO:0009152 | purine ribonucleotide biosynthetic process | 110 | 3 | 0.2 | 0.0009 | 0.0024 |
| U, annuals | GO:0006898 | receptor-mediated endocytosis | 26 | 2 | 0.05 | 0.001 | 0.00243 |
| U, annuals | GO:0006164 | purine nucleotide biosynthetic process | 119 | 3 | 0.21 | 0.0012 | 0.0026 |
| U, annuals | GO:0030100 | regulation of endocytosis | 29 | 2 | 0.05 | 0.0012 | 0.0026 |
| U, annuals | GO:0055086 | nucleobase-containing small molecule metabolic process | 284 | 4 | 0.51 | 0.0014 | 0.0026 |
| U, annuals | GO:0009260 | ribonucleotide biosynthetic process | 127 | 3 | 0.23 | 0.0014 | 0.0026 |
| U, annuals | GO:0046390 | ribose phosphate biosynthetic process | 127 | 3 | 0.23 | 0.0014 | 0.0026 |
| U, annuals | GO:0072522 | purine-containing compound biosynthetic process | 127 | 3 | 0.23 | 0.0014 | 0.0026 |
| U, annuals | GO:0006182 | cGMP biosynthetic process | 33 | 2 | 0.06 | 0.0016 | 0.0026 |
| U, annuals | GO:0009187 | cyclic nucleotide metabolic process | 33 | 2 | 0.06 | 0.0016 | 0.0026 |
| U, annuals | GO:0009190 | cyclic nucleotide biosynthetic process | 33 | 2 | 0.06 | 0.0016 | 0.0026 |
| U, annuals | GO:0046068 | cGMP metabolic process | 33 | 2 | 0.06 | 0.0016 | 0.0026 |
| U, annuals | GO:0052652 | cyclic purine nucleotide metabolic process | 33 | 2 | 0.06 | 0.0016 | 0.0026 |
| U, annuals | GO:0018282 | metal incorporation into metallo-sulfur cluster | 1 | 1 | 0 | 0.0018 | 0.00277 |
| U, annuals | GO:0018283 | iron incorporation into metallo-sulfur cluster | 1 | 1 | 0 | 0.0018 | 0.00277 |
| U, annuals | GO:1903329 | regulation of iron-sulfur cluster assembly | 1 | 1 | 0 | 0.0018 | 0.00277 |
| U, annuals | GO:0007267 | cell-cell signaling | 41 | 2 | 0.07 | 0.0024 | 0.0036 |
| U, annuals | GO:0009150 | purine ribonucleotide metabolic process | 159 | 3 | 0.29 | 0.0027 | 0.00373 |
| U, annuals | GO:0009165 | nucleotide biosynthetic process | 159 | 3 | 0.29 | 0.0027 | 0.00373 |
| U, annuals | GO:1901293 | nucleoside phosphate biosynthetic process | 159 | 3 | 0.29 | 0.0027 | 0.00373 |
| U, annuals | GO:0009259 | ribonucleotide metabolic process | 176 | 3 | 0.32 | 0.0036 | 0.00485 |
| U, annuals | GO:0019693 | ribose phosphate metabolic process | 186 | 3 | 0.34 | 0.0042 | 0.00548 |
| U, annuals | GO:0006163 | purine nucleotide metabolic process | 187 | 3 | 0.34 | 0.0042 | 0.00548 |
| U, annuals | GO:0009060 | aerobic respiration | 55 | 2 | 0.1 | 0.0043 | 0.00548 |
| U, annuals | GO:0045007 | depurination | 3 | 1 | 0.01 | 0.0054 | 0.00642 |
| U, annuals | GO:0006304 | DNA modification | 62 | 2 | 0.11 | 0.0054 | 0.00642 |
| U, annuals | GO:0072521 | purine-containing compound metabolic process | 205 | 3 | 0.37 | 0.0055 | 0.00642 |
| U, annuals | GO:0019637 | organophosphate metabolic process | 417 | 4 | 0.75 | 0.0056 | 0.00644 |
| U, annuals | GO:0045333 | cellular respiration | 69 | 2 | 0.12 | 0.0067 | 0.00755 |
| U, annuals | GO:0051050 | positive regulation of transport | 72 | 2 | 0.13 | 0.0073 | 0.00797 |
| U, annuals | GO:0051128 | regulation of cellular component organization | 450 | 4 | 0.81 | 0.0073 | 0.00797 |
| U, annuals | GO:0009117 | nucleotide metabolic process | 236 | 3 | 0.43 | 0.0081 | 0.00864 |
| U, annuals | GO:0060627 | regulation of vesicle-mediated transport | 77 | 2 | 0.14 | 0.0083 | 0.00868 |
| U, annuals | GO:0006307 | DNA dealkylation involved in DNA repair | 5 | 1 | 0.01 | 0.009 | 0.0093 |
| U, annuals | GO:0006897 | endocytosis | 85 | 2 | 0.15 | 0.01 | 0.00998 |
| P, U, perennials | GO:0042773 | ATP synthesis coupled electron transport | 51 | 5 | 0.09 | 2.3e-08 | 9.1e-07 |
| P, U, perennials | GO:0006119 | oxidative phosphorylation | 52 | 5 | 0.09 | 2.6e-08 | 9.1e-07 |
| P, U, perennials | GO:0022904 | respiratory electron transport chain | 64 | 5 | 0.11 | 7.4e-08 | 1.73e-06 |
| P, U, perennials | GO:0043953 | protein transport by the Tat complex | 10 | 3 | 0.02 | 5.4e-07 | 9.45e-06 |
| P, U, perennials | GO:0009060 | aerobic respiration | 100 | 5 | 0.17 | 7.0e-07 | 9.8e-06 |
| P, U, perennials | GO:0009558 | embryo sac cellularization | 13 | 3 | 0.02 | 1.3e-06 | 1.52e-05 |
| P, U, perennials | GO:0045333 | cellular respiration | 130 | 5 | 0.22 | 2.6e-06 | 2.6e-05 |
| P, U, perennials | GO:0007349 | cellularization | 17 | 3 | 0.03 | 3.0e-06 | 2.625e-05 |
| P, U, perennials | GO:0046034 | ATP metabolic process | 152 | 5 | 0.26 | 5.5e-06 | 3.55e-05 |
| P, U, perennials | GO:0022900 | electron transport chain | 156 | 5 | 0.27 | 6.3e-06 | 3.55e-05 |
| P, U, perennials | GO:0009126 | purine nucleoside monophosphate metabolic process | 160 | 5 | 0.27 | 7.1e-06 | 3.55e-05 |
| P, U, perennials | GO:0009144 | purine nucleoside triphosphate metabolic process | 160 | 5 | 0.27 | 7.1e-06 | 3.55e-05 |
| P, U, perennials | GO:0009167 | purine ribonucleoside monophosphate metabolic process | 160 | 5 | 0.27 | 7.1e-06 | 3.55e-05 |
| P, U, perennials | GO:0009205 | purine ribonucleoside triphosphate metabolic process | 160 | 5 | 0.27 | 7.1e-06 | 3.55e-05 |
| P, U, perennials | GO:0009199 | ribonucleoside triphosphate metabolic process | 169 | 5 | 0.29 | 9.3e-06 | 4.34e-05 |
| P, U, perennials | GO:0015980 | energy derivation by oxidation of organic compounds | 175 | 5 | 0.3 | 1.1e-05 | 4.8125e-05 |
| P, U, perennials | GO:0009141 | nucleoside triphosphate metabolic process | 179 | 5 | 0.31 | 1.2e-05 | 4.94e-05 |
| P, U, perennials | GO:0009161 | ribonucleoside monophosphate metabolic process | 180 | 5 | 0.31 | 1.3e-05 | 5.06e-05 |
| P, U, perennials | GO:0009123 | nucleoside monophosphate metabolic process | 204 | 5 | 0.35 | 2.3e-05 | 8.47e-05 |
| P, U, perennials | GO:0090307 | mitotic spindle assembly | 35 | 3 | 0.06 | 2.9e-05 | 0.0001 |
| P, U, perennials | GO:0018343 | protein farnesylation | 6 | 2 | 0.01 | 4.2e-05 | 0.00014 |
| P, U, perennials | GO:0051225 | spindle assembly | 54 | 3 | 0.09 | 0.0001 | 0.00035 |
| P, U, perennials | GO:0018344 | protein geranylgeranylation | 10 | 2 | 0.02 | 0.0001 | 0.0004 |
| P, U, perennials | GO:0009150 | purine ribonucleotide metabolic process | 301 | 5 | 0.51 | 0.0002 | 0.00043 |
| P, U, perennials | GO:0007052 | mitotic spindle organization | 62 | 3 | 0.11 | 0.0002 | 0.00043 |
| P, U, perennials | GO:1902850 | microtubule cytoskeleton organization involved in mitosis | 62 | 3 | 0.11 | 0.0002 | 0.00043 |
| P, U, perennials | GO:0009259 | ribonucleotide metabolic process | 336 | 5 | 0.57 | 0.0002 | 0.00062 |
| P, U, perennials | GO:0018342 | protein prenylation | 15 | 2 | 0.03 | 0.0003 | 7e-04 |
| P, U, perennials | GO:0097354 | prenylation | 15 | 2 | 0.03 | 0.0003 | 7e-04 |
| P, U, perennials | GO:0019693 | ribose phosphate metabolic process | 356 | 5 | 0.61 | 0.0003 | 0.00075 |
| P, U, perennials | GO:0006163 | purine nucleotide metabolic process | 359 | 5 | 0.61 | 0.0003 | 0.00075 |
| P, U, perennials | GO:0048768 | root hair cell tip growth | 17 | 2 | 0.03 | 0.0004 | 0.00083 |
| P, U, perennials | GO:0007051 | spindle organization | 91 | 3 | 0.16 | 0.0005 | 0.00106 |
| P, U, perennials | GO:0072521 | purine-containing compound metabolic process | 399 | 5 | 0.68 | 0.0005 | 0.00109 |
| P, U, perennials | GO:0002098 | tRNA wobble uridine modification | 21 | 2 | 0.04 | 0.0006 | 0.00116 |
| P, U, perennials | GO:0002097 | tRNA wobble base modification | 25 | 2 | 0.04 | 0.0008 | 0.00161 |
| P, U, perennials | GO:0006091 | generation of precursor metabolites and energy | 455 | 5 | 0.78 | 0.001 | 0.00176 |
| P, U, perennials | GO:0009117 | nucleotide metabolic process | 459 | 5 | 0.79 | 0.001 | 0.00176 |
| P, U, perennials | GO:0009561 | megagametogenesis | 116 | 3 | 0.2 | 0.001 | 0.00176 |
| P, U, perennials | GO:0000911 | cytokinesis by cell plate formation | 117 | 3 | 0.2 | 0.001 | 0.00176 |
| P, U, perennials | GO:0006753 | nucleoside phosphate metabolic process | 463 | 5 | 0.79 | 0.001 | 0.00176 |
| P, U, perennials | GO:0000070 | mitotic sister chromatid segregation | 118 | 3 | 0.2 | 0.0011 | 0.00176 |
| P, U, perennials | GO:0065002 | intracellular protein transmembrane transport | 119 | 3 | 0.2 | 0.0011 | 0.00176 |
| P, U, perennials | GO:0032506 | cytokinetic process | 124 | 3 | 0.21 | 0.0012 | 0.0019 |
| P, U, perennials | GO:1902410 | mitotic cytokinetic process | 124 | 3 | 0.21 | 0.0012 | 0.0019 |
| P, U, perennials | GO:0071806 | protein transmembrane transport | 128 | 3 | 0.22 | 0.0013 | 0.00204 |
| P, U, perennials | GO:0016310 | phosphorylation | 1663 | 9 | 2.84 | 0.0014 | 0.0021 |
| P, U, perennials | GO:0090421 | embryonic meristem initiation | 36 | 2 | 0.06 | 0.0017 | 0.00251 |
| P, U, perennials | GO:0006796 | phosphate-containing compound metabolic process | 2463 | 11 | 4.21 | 0.0018 | 0.00251 |
| P, U, perennials | GO:0007018 | microtubule-based movement | 144 | 3 | 0.25 | 0.0019 | 0.00263 |
| P, U, perennials | GO:0006793 | phosphorus metabolic process | 2497 | 11 | 4.27 | 0.002 | 0.0027 |
| P, U, perennials | GO:0000819 | sister chromatid segregation | 156 | 3 | 0.27 | 0.0024 | 0.00318 |
| P, U, perennials | GO:0055086 | nucleobase-containing small molecule metabolic process | 563 | 5 | 0.96 | 0.0025 | 0.00326 |
| P, U, perennials | GO:0042744 | hydrogen peroxide catabolic process | 45 | 2 | 0.08 | 0.0027 | 0.00338 |
| P, U, perennials | GO:0009653 | anatomical structure morphogenesis | 1825 | 9 | 3.12 | 0.0027 | 0.00338 |
| P, U, perennials | GO:0009880 | embryonic pattern specification | 47 | 2 | 0.08 | 0.0029 | 0.00359 |
| P, U, perennials | GO:0140014 | mitotic nuclear division | 176 | 3 | 0.3 | 0.0033 | 0.00399 |
| P, U, perennials | GO:0030488 | tRNA methylation | 52 | 2 | 0.09 | 0.0036 | 0.00422 |
| P, U, perennials | GO:0000281 | mitotic cytokinesis | 188 | 3 | 0.32 | 0.004 | 0.00466 |
| P, U, perennials | GO:0010014 | meristem initiation | 57 | 2 | 0.1 | 0.0043 | 0.00489 |
| P, U, perennials | GO:0061640 | cytoskeleton-dependent cytokinesis | 202 | 3 | 0.35 | 0.0049 | 0.0055 |
| P, U, perennials | GO:0008360 | regulation of cell shape | 62 | 2 | 0.11 | 0.005 | 0.00558 |
| P, U, perennials | GO:0048507 | meristem development | 427 | 4 | 0.73 | 0.0058 | 0.00632 |
| P, U, perennials | GO:0042743 | hydrogen peroxide metabolic process | 71 | 2 | 0.12 | 0.0065 | 0.00704 |
| P, U, perennials | GO:0000910 | cytokinesis | 229 | 3 | 0.39 | 0.0069 | 0.00731 |
| P, U, perennials | GO:0000226 | microtubule cytoskeleton organization | 238 | 3 | 0.41 | 0.0077 | 0.008 |
| P, U, perennials | GO:0001708 | cell fate specification | 82 | 2 | 0.14 | 0.0086 | 0.00889 |
| P, U, perennials | GO:0009553 | embryo sac development | 252 | 3 | 0.43 | 0.009 | 0.00909 |
| P, U, perennials | GO:0140694 | non-membrane-bounded organelle assembly | 255 | 3 | 0.44 | 0.0093 | 0.00925 |
| P, perennials | GO:0006338 | chromatin remodeling | 290 | 7 | 0.96 | 3.50E-05 | 0.00322 |
| P, perennials | GO:0031508 | pericentric heterochromatin formation | 6 | 2 | 0.02 | 0.0002 | 0.00491 |
| P, perennials | GO:0140719 | constitutive heterochromatin formation | 6 | 2 | 0.02 | 0.0002 | 0.00491 |
| P, perennials | GO:0032467 | positive regulation of cytokinesis | 7 | 2 | 0.02 | 0.0002 | 0.00506 |
| P, perennials | GO:0045814 | negative regulation of gene expression | 108 | 4 | 0.36 | 0.0004 | 0.0056 |
| P, perennials | GO:0031055 | chromatin remodeling at centromere | 10 | 2 | 0.03 | 0.0005 | 0.0056 |
| P, perennials | GO:0006325 | chromatin organization | 457 | 7 | 1.52 | 0.0006 | 0.0056 |
| P, perennials | GO:0040029 | epigenetic regulation of gene expression | 217 | 5 | 0.72 | 0.0007 | 0.0056 |
| P, perennials | GO:0071824 | protein-DNA complex organization | 499 | 7 | 1.66 | 0.001 | 0.00774 |
| P, perennials | GO:0045787 | positive regulation of cell cycle | 62 | 3 | 0.21 | 0.0011 | 0.00778 |
| P, perennials | GO:0010467 | gene expression | 2405 | 16 | 7.99 | 0.0017 | 0.0086 |
| P, perennials | GO:0031323 | regulation of cellular metabolic process | 2160 | 15 | 7.18 | 0.0017 | 0.0086 |
| P, perennials | GO:0010468 | regulation of gene expression | 1718 | 13 | 5.71 | 0.0019 | 0.0086 |
| P, perennials | GO:0031398 | positive regulation of protein ubiquitination | 20 | 2 | 0.07 | 0.002 | 0.0086 |
| P, perennials | GO:1903322 | positive regulation of protein modification by small protein conjugation or removal | 20 | 2 | 0.07 | 0.002 | 0.0086 |
| P, perennials | GO:0019222 | regulation of metabolic process | 2207 | 15 | 7.34 | 0.0021 | 0.0086 |
| P, perennials | GO:0032465 | regulation of cytokinesis | 21 | 2 | 0.07 | 0.0022 | 0.0086 |
| P, perennials | GO:0034508 | centromere complex assembly | 23 | 2 | 0.08 | 0.0026 | 0.0086 |
| P, perennials | GO:0060255 | regulation of macromolecule metabolic process | 2006 | 14 | 6.67 | 0.0026 | 0.0086 |
| P, perennials | GO:0010556 | regulation of macromolecule biosynthetic process | 1803 | 13 | 5.99 | 0.003 | 0.0086 |
| P, perennials | GO:0012502 | induction of programmed cell death | 1 | 1 | 0 | 0.0033 | 0.0086 |
| P, perennials | GO:0016557 | peroxisome membrane biogenesis | 1 | 1 | 0 | 0.0033 | 0.0086 |
| P, perennials | GO:0045046 | protein import into peroxisome membrane | 1 | 1 | 0 | 0.0033 | 0.0086 |
| P, perennials | GO:0051926 | negative regulation of calcium ion transport | 1 | 1 | 0 | 0.0033 | 0.0086 |
| P, perennials | GO:0097352 | autophagosome maturation | 1 | 1 | 0 | 0.0033 | 0.0086 |
| P, perennials | GO:1900130 | regulation of lipid binding | 1 | 1 | 0 | 0.0033 | 0.0086 |
| P, perennials | GO:1900131 | negative regulation of lipid binding | 1 | 1 | 0 | 0.0033 | 0.0086 |
| P, perennials | GO:1901019 | regulation of calcium ion transmembrane transporter activity | 1 | 1 | 0 | 0.0033 | 0.0086 |
| P, perennials | GO:1901020 | negative regulation of calcium ion transmembrane transporter activity | 1 | 1 | 0 | 0.0033 | 0.0086 |
| P, perennials | GO:1901339 | regulation of store-operated calcium channel activity | 1 | 1 | 0 | 0.0033 | 0.0086 |
| P, perennials | GO:1901340 | negative regulation of store-operated calcium channel activity | 1 | 1 | 0 | 0.0033 | 0.0086 |
| P, perennials | GO:1903170 | negative regulation of calcium ion transmembrane transport | 1 | 1 | 0 | 0.0033 | 0.0086 |
| P, perennials | GO:0071704 | organic substance metabolic process | 6059 | 27 | 20.14 | 0.0038 | 0.0086 |
| P, perennials | GO:0031396 | regulation of protein ubiquitination | 28 | 2 | 0.09 | 0.0038 | 0.0086 |
| P, perennials | GO:0031326 | regulation of cellular biosynthetic process | 1865 | 13 | 6.2 | 0.0041 | 0.0086 |
| P, perennials | GO:0051674 | localization of cell | 30 | 2 | 0.1 | 0.0044 | 0.0086 |
| P, perennials | GO:0051781 | positive regulation of cell division | 30 | 2 | 0.1 | 0.0044 | 0.0086 |
| P, perennials | GO:0090068 | positive regulation of cell cycle process | 30 | 2 | 0.1 | 0.0044 | 0.0086 |
| P, perennials | GO:1903320 | regulation of protein modification by small protein conjugation or removal | 30 | 2 | 0.1 | 0.0044 | 0.0086 |
| P, perennials | GO:0009889 | regulation of biosynthetic process | 1878 | 13 | 6.24 | 0.0044 | 0.0086 |
| P, perennials | GO:0044237 | cellular metabolic process | 6113 | 27 | 20.32 | 0.0046 | 0.0086 |
| P, perennials | GO:0010025 | wax biosynthetic process | 31 | 2 | 0.1 | 0.0047 | 0.0086 |
| P, perennials | GO:0009058 | biosynthetic process | 3760 | 20 | 12.5 | 0.0049 | 0.0086 |
| P, perennials | GO:0010166 | wax metabolic process | 33 | 2 | 0.11 | 0.0053 | 0.0086 |
| P, perennials | GO:0006457 | protein folding | 108 | 3 | 0.36 | 0.0054 | 0.0086 |
| P, perennials | GO:0008152 | metabolic process | 6613 | 28 | 21.98 | 0.0061 | 0.0086 |
| P, perennials | GO:1901570 | fatty acid derivative biosynthetic process | 36 | 2 | 0.12 | 0.0063 | 0.0086 |
| P, perennials | GO:0010629 | negative regulation of gene expression | 523 | 6 | 1.74 | 0.0066 | 0.0086 |
| P, perennials | GO:0008631 | intrinsic apoptotic signaling pathway in response to oxidative stress | 2 | 1 | 0.01 | 0.0066 | 0.0086 |
| P, perennials | GO:0010799 | regulation of peptidyl-threonine phosphorylation | 2 | 1 | 0.01 | 0.0066 | 0.0086 |
| P, perennials | GO:0010800 | positive regulation of peptidyl-threonine phosphorylation | 2 | 1 | 0.01 | 0.0066 | 0.0086 |
| P, perennials | GO:0045007 | depurination | 2 | 1 | 0.01 | 0.0066 | 0.0086 |
| P, perennials | GO:0046521 | sphingoid catabolic process | 2 | 1 | 0.01 | 0.0066 | 0.0086 |
| P, perennials | GO:1902175 | regulation of oxidative stress-induced intrinsic apoptotic signaling pathway | 2 | 1 | 0.01 | 0.0066 | 0.0086 |
| P, perennials | GO:1903071 | positive regulation of ER-associated ubiquitin-dependent protein catabolic process | 2 | 1 | 0.01 | 0.0066 | 0.0086 |
| P, perennials | GO:1903169 | regulation of calcium ion transmembrane transport | 2 | 1 | 0.01 | 0.0066 | 0.0086 |
| P, perennials | GO:1904294 | positive regulation of ERAD pathway | 2 | 1 | 0.01 | 0.0066 | 0.0086 |
| P, perennials | GO:1905898 | positive regulation of response to endoplasmic reticulum stress | 2 | 1 | 0.01 | 0.0066 | 0.0086 |
| P, perennials | GO:0043067 | regulation of programmed cell death | 119 | 3 | 0.4 | 0.007 | 0.00894 |
| P, perennials | GO:0006355 | regulation of DNA-templated transcription | 1318 | 10 | 4.38 | 0.0079 | 0.00994 |
| P, perennials | GO:2001141 | regulation of RNA biosynthetic process | 1349 | 10 | 4.48 | 0.0093 | 0.00994 |
| P, perennials | GO:0051276 | chromosome organization | 565 | 6 | 1.88 | 0.0096 | 0.00994 |
| P, perennials | GO:0009059 | macromolecule biosynthetic process | 2822 | 16 | 9.38 | 0.0097 | 0.00994 |
| P, perennials | GO:0044249 | cellular biosynthetic process | 3665 | 19 | 12.18 | 0.0099 | 0.00994 |
| P, perennials | GO:0006419 | alanyl-tRNA aminoacylation | 3 | 1 | 0.01 | 0.0099 | 0.00994 |
| P, perennials | GO:0006850 | mitochondrial pyruvate transmembrane transport | 3 | 1 | 0.01 | 0.0099 | 0.00994 |
| P, perennials | GO:0010447 | response to acidic pH | 3 | 1 | 0.01 | 0.0099 | 0.00994 |
| P, perennials | GO:2000306 | positive regulation of photomorphogenesis | 3 | 1 | 0.01 | 0.0099 | 0.00994 |
| U, perennials | GO:0042773 | ATP synthesis coupled electron transport | 28 | 3 | 0.14 | 0.0004 | 0.00347 |
| U, perennials | GO:0006119 | oxidative phosphorylation | 29 | 3 | 0.15 | 0.0004 | 0.00347 |
| U, perennials | GO:0090230 | regulation of centromere complex assembly | 7 | 2 | 0.04 | 0.0005 | 0.00347 |
| U, perennials | GO:0022904 | respiratory electron transport chain | 35 | 3 | 0.18 | 0.0007 | 0.00355 |
| U, perennials | GO:0005992 | trehalose biosynthetic process | 15 | 2 | 0.08 | 0.0025 | 0.00826 |
| U, perennials | GO:0009060 | aerobic respiration | 55 | 3 | 0.28 | 0.0027 | 0.00826 |
| U, perennials | GO:0005991 | trehalose metabolic process | 16 | 2 | 0.08 | 0.0029 | 0.00826 |
| U, perennials | GO:0034508 | centromere complex assembly | 19 | 2 | 0.1 | 0.0041 | 0.00838 |
| U, perennials | GO:0046351 | disaccharide biosynthetic process | 20 | 2 | 0.1 | 0.0045 | 0.00838 |
| U, perennials | GO:0000097 | sulfur amino acid biosynthetic process | 21 | 2 | 0.11 | 0.005 | 0.00838 |
| U, perennials | GO:0045333 | cellular respiration | 69 | 3 | 0.35 | 0.0051 | 0.00838 |
| U, perennials | GO:0080163 | regulation of protein serine/threonine phosphatase activity | 1 | 1 | 0.01 | 0.0051 | 0.00838 |
| U, perennials | GO:0032886 | regulation of microtubule-based process | 22 | 2 | 0.11 | 0.0055 | 0.00838 |
| U, perennials | GO:0022900 | electron transport chain | 76 | 3 | 0.39 | 0.0066 | 0.00919 |
| U, perennials | GO:0046034 | ATP metabolic process | 80 | 3 | 0.41 | 0.0076 | 0.00919 |
| U, perennials | GO:0009126 | purine nucleoside monophosphate metabolic process | 83 | 3 | 0.42 | 0.0085 | 0.00919 |
| U, perennials | GO:0009167 | purine ribonucleoside monophosphate metabolic process | 83 | 3 | 0.42 | 0.0085 | 0.00919 |
| U, perennials | GO:0009144 | purine nucleoside triphosphate metabolic process | 84 | 3 | 0.43 | 0.0087 | 0.00919 |
| U, perennials | GO:0009205 | purine ribonucleoside triphosphate metabolic process | 84 | 3 | 0.43 | 0.0087 | 0.00919 |
| U, perennials | GO:0009199 | ribonucleoside triphosphate metabolic process | 88 | 3 | 0.45 | 0.0099 | 0.00992 |

**Supplementary Table 3.**
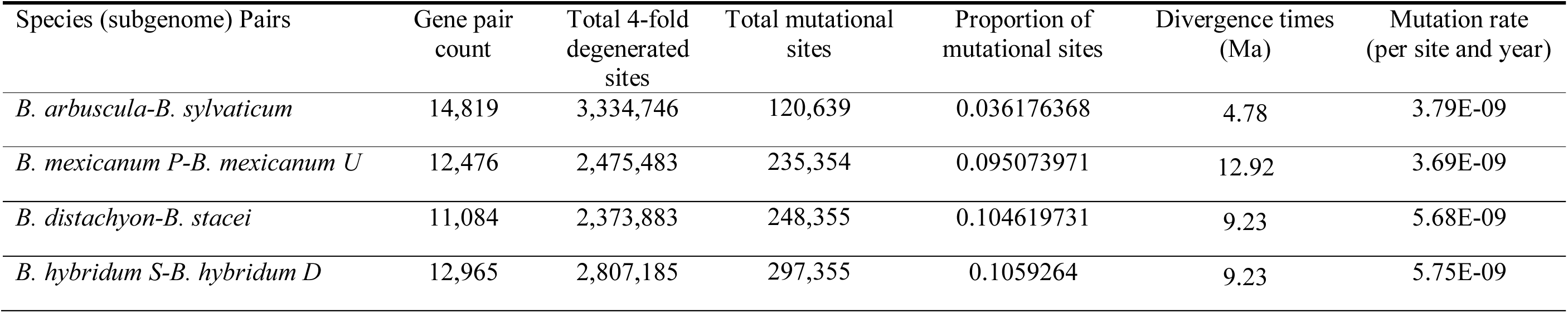
Mutation rate parameters among the studied *Brachypodium* (sub)genomes and groups.

**Supplementary Table 4.** Statistics on the proportion of pan-gene count (%) of each core, softcore, dispensable and private categories, in the studied *Brachypodium* genomes. Newly assembled *B. arbuscula* and *B. mexicanum* (P, U) (sub)genomes are highlighted in bold.

| (Sub)Genomes | Core | Softcore | Dispensable | Private |
| --- | --- | --- | --- | --- |
| ***B. arbuscula*** | **54.78** | **21.51** | **17.11** | **1.30** |
| ***B. mexicanum P*** | **49.27** | **17.46** | **18.94** | **3.27** |
| ***B. mexicanum U*** | **51.31** | **17.11** | **17.84** | **2.98** |
| *B. distachyon* | 65.96 | 16.61 | 13.29 | 0.40 |
| *B. hybridum D* | 60.30 | 23.66 | 12.09 | 0.55 |
| *B. hybridum S* | 60.19 | 23.91 | 12.80 | 0.23 |
| *B. stacei* | 58.85 | 23.54 | 13.69 | 0.33 |
| *B. sylvaticum* | 57.25 | 20.27 | 14.26 | 1.27 |
| Average | 57.24 | 20.51 | 15.00 | 1.29 |

**Supplementary Table 5.** Proportion (%) of repetitive elements in the eight sub(genomes) of *Brachypodium* under study. Newly assembled *B. arbuscula* and *B. mexicanum* (P, U) (sub)genomes are highlighted in bold.

| (Sub)Genomes | LTR | TIR | LINE | Helitron | repeat_region | Total |
| --- | --- | --- | --- | --- | --- | --- |
| ***B. arbuscula*** | **16.44** | **11.37** | **0.04** | **10.09** | **0.33** | **38.27** |
| ***B. mexicanum* P** | **60.00** | **7.86** | **0** | **4.91** | **0.12** | **72.89** |
| ***B. mexicanum* U** | **55.98** | **7.79** | **0** | **7.19** | **0.36** | **71.32** |
| *B. distachyon* | 17.50 | 8.81 | 0.04 | 9.56 | 0.60 | 36.51 |
| *B. hybridum* D | 20.90 | 8.54 | 0.16 | 10.39 | 0.45 | 40.44 |
| *B. hybridum* S | 12.16 | 8.42 | 0.1 | 10.12 | 4.66 | 35.46 |
| *B. stacei* | 14.37 | 7.86 | 0.02 | 10.17 | 5.83 | 38.25 |
| *B. sylvaticum* | 27.59 | 9.9 | 0.06 | 11.45 | 0.6 | 49.6 |
| Average | 28.12 | 8.82 | 0.05 | 9.23 | 1.62 | 47.84 |

**Supplementary Table 6.** *Solo*/*intact* TE ratio in the eight sub(genomes) of *Brachypodium* under study. Newly assembled *B. arbuscula* and *B. mexicanum* (P, U) (sub)genomes are highlighted in bold.

| Sub/genomes | total *solo* TE<br>count | total *intact*<br>TE count | *solo/intact* ratio |
| --- | --- | --- | --- |
| ***Brachypodium arbuscula*** | **4,334** | **1,718** | **2.52** |
| ***Brachypodium mexicanum* P** | **57,786** | **12,941** | **4.47** |
| ***Brachypodium mexicanum* U** | **54,117** | **12,010** | **4.51** |
| *Brachypodium distachyon* | 9,745 | 1,055 | 9.24 |
| *Brachypodium hybridum* D | 7,940 | 1,717 | 4.62 |
| *Brachypodium hybridum* S | 5,300 | 837 | 6.33 |
| *Brachypodium stacei* | 5,437 | 1,047 | 5.19 |
| *Brachypodium sylvaticum* | 9,653 | 2,544 | 3.79 |

**Supplementary Table 7.** Proportion of Pan-TE classes with respect to the total TEs (%) for the eight sub(genomes) of *Brachypodium* under study. Newly assembled *B. arbuscula* and *B. mexicanum* (P, U) (sub)genomes are highlighted in bold.

| (Sub)Genomes | core | softcore | dispensable | private |
| --- | --- | --- | --- | --- |
| ***B. arbuscula*** | **27.77** | **12.50** | **58.15** | **1.58** |
| ***B. mexicanum* P** | **16.73** | **5.98** | **71.60** | **5.69** |
| ***B. mexicanum* U** | **11.49** | **3.82** | **67.68** | **17.02** |
| *B. distachyon* | 27.74 | 17.93 | 53.84 | 0.49 |
| *B. hybridum* D | 28.36 | 17.73 | 52.85 | 1.05 |
| *B. hybridum* S | 16.35 | 10.09 | 73.28 | 0.28 |
| *B. stacei* | 24.98 | 15.24 | 59.29 | 0.49 |
| *B. sylvaticum* | 27.76 | 16.49 | 53.52 | 2.23 |
| Average | 22.65 | 12.47 | 61.28 | 3.60 |

**Supplementary Table 8.** Pan-TE length (bp) and proportion (%) in genome length of the eight *Brachypodium* sub(genomes) under study. Newly assembled *B. arbuscula* and *B. mexicanum* (P, U) (sub)genomes are highlighted in bold.

| (Sub)Genomes | core<br>length | softcore<br>length | dispensable<br>length | private<br>length | core<br>proportion | softcore<br>proportion | dispensable<br>proportion | private<br>proportion |
| --- | --- | --- | --- | --- | --- | --- | --- | --- |
| ***B. arbuscula*** | **25,807,770** | **19,077,944** | **42,417,313** | **1,037,236** | **9.51** | **7.03** | **15.63** | **0.38** |
| ***B. mexicanum* P** | **66,586,074** | **16,590,135** | **358,823,412** | **21,414,606** | **8.26** | **2.06** | **44.51** | **2.66** |
| ***B. mexicanum* U** | **49,252,522** | **10,470,984** | **307,336,892** | **47,278,704** | **7.11** | **1.51** | **44.35** | **6.82** |
| *B. distachyon* | 27,784,657 | 24,408,390 | 35,214,051 | 971,982 | 10.29 | 9.04 | 13.04 | 0.36 |
| *B. hybridum* D | 32,201,801 | 30,240,379 | 36,633,891 | 1,648,381 | 11.70 | 10.99 | 13.31 | 0.60 |
| *B. hybridum* S | 19,734,039 | 12,908,877 | 48,854,790 | 554,674 | 7.73 | 5.06 | 19.15 | 0.22 |
| *B. stacei* | 22,028,043 | 16,428,846 | 51,123,984 | 303,231 | 9.03 | 6.73 | 20.96 | 0.12 |
| *B. sylvaticum* | 40,736,638 | 38,625,357 | 57,626,227 | 2,403,918 | 11.92 | 11.30 | 16.87 | 0.70 |
| Average | 35,516,443 | 21,093,864 | 117,253,820 | 9,451,592 | 9.44 | 6.72 | 23.48 | 1.48 |

**Supplementary data 1.** Details on the genome assembly and annotation of *Brachypodium arbuscula* and *B. mexicanum*.

## Genome assembly

Genome assemblies o*f B. mexicanum* and *B. arbuscula* were generated using PacBio Sequel long reads assembled with MECAT^1^ and subsequently polished using ARROW^2^. Hi-C Illumina reads were aligned to the initial contig assemblies using bwa mem^3^, and chromosome-scale scaffolding was performed based on Hi-C contact maps visualized in Juicebox^4^. Manual inspection of the Hi-C interaction maps identified 17 and 4 assembly misjoins in the *B. mexicanum* and *B. arbuscula* assemblies, respectively. Following correction, 670 joins were introduced to generate the 20 chromosomes of *B. mexicanum*, whereas 180 joins were applied to construct the nine chromosomes of *B. arbuscula.* Chromosomes of *B. mexicanum* were oriented using the Bd21 genome as reference and numbered according to the published karyotype, whereas chromosomes of *B. arbuscula* were ordered and oriented according to the version 2.0 assembly of *Brachypodium sylvaticum* var. AIN1. Scaffold junctions were padded with 10,000 Ns. Contigs terminating with telomeric repeat sequences ((TTTAGGG)n) were used to verify chromosome orientation. Unplaced scaffolds were screened against bacterial proteins, organelle genomes, and the GenBank nonredundant (nr) database, and contaminant sequences were removed. To eliminate redundant overlaps between adjacent contigs within chromosomes, contig ends were aligned using BLAT^5^, and overlapping duplicated sequences were collapsed. This procedure collapsed 198 and 72 adjacent contig pairs in the *B. mexicanum* and *B. arbuscula* assemblies, respectively. The chromosome-scale assemblies were further polished using Illumina paired-end reads (2 × 250 bp; 400-bp insert size). Reads were aligned with bwa mem, and homozygous SNPs and INDELs were identified using the UnifiedGenotyper module of GATK^6^. For *B. mexicanum*, polishing was performed using 40× Illumina coverage, correcting 309 homozygous SNPs and 12,379 homozygous INDELs. For *B. arbuscula,* 120× Illumina coverage was used, correcting 306 homozygous SNPs and 12,219 homozygous INDELs. The final *B. arbuscula* v3.1 assembly contained 283.6 Mb of sequence assembled into 204 contigs with a contig N50 of 4.1 Mb, with 96.0% of assembled bases integrated into chromosomes. The final *B. mexicanum* v2.1 assembly consisted of 1,530.5 Mb of sequence assembled into 812 contigs with a contig N50 of 6.4 Mb, with 97.94% of assembled bases anchored to chromosomes. Assembly completeness was evaluated by aligning available RNA-seq reads to the final assemblies. RNA-seq mapping rates reached 100.0% for *B. mexicanum* and 98.71% for *B. arbuscula*, indicating high completeness of the euchromatic gene space.

## Genome annotation

Gene annotation of *Brachypodium mexicanum* and *B. arbuscula* was performed using an integrated pipeline combining transcriptome evidence, protein homology, and *ab initio* gene prediction. Genome-guided transcript assemblies were generated from stranded Illumina RNA-seq data using PERTRAN, which aligns short reads to the genome with GSNAP^7^, followed by alignment validation, realignment, correction, and splice graph construction. Approximately one billion and 414 million paired-end (2 × 150 bp) RNA-seq read pairs were used for *B. mexicanum* and *B. arbuscula*, respectively.

Full-length transcript evidence was generated from PacBio Iso-Seq data using a genome-guided correction pipeline. Circular consensus sequence (CCS) reads were aligned to the genome using GMAP^8^, corrected for small splice-junction indels when necessary, and collapsed into nonredundant transcript models by clustering alignments sharing identical intron structures or exhibiting ≥95% overlap for single-exon transcripts. Approximately 18 million CCS reads from *B. mexicanum* and 5 million CCS reads from *B. arbuscula* generated ∼1.3 million and 319,000 putative full-length transcripts, respectively. RNA-seq transcript assemblies were subsequently refined with PASA^9^, producing approximately one million transcript assemblies for *B. mexicanum* and 227,916 transcript assemblies for *B. arbuscula*.

Candidate gene loci were defined based on transcript alignments and EXONERATE^10^ alignments of proteins from representative angiosperm species together with the Swiss-Prot database to repeat-soft-masked genomes. Repeat masking was performed using RepeatMasker^11^ with a repeat library composed of de novo repeats identified by RepeatModeler^12^ and RepBase sequences. Gene models were predicted using FGENESH+, FGENESH_EST^13^, EXONERATE, PASA-derived ORFs, and AUGUSTUS^14^. For *B. mexicanum*, AUGUSTUS gene prediction was implemented through BRAKER1^15^, whereas for *B. arbuscula* AUGUSTUS was trained using high-confidence PASA-derived ORFs together with intron hints generated from RNA-seq alignments. For each candidate locus, the highest-scoring gene model was selected based on transcript support, protein homology, and overlap with repetitive sequences. Selected models were further refined using PASA to improve splice junctions, annotate untranslated regions (UTRs), and identify alternative splice isoforms. The refined proteins were compared against reference proteomes to calculate Cscore, defined as the ratio between the BLASTP score and the mutual best-hit BLASTP score, together with protein coverage. Gene models were retained if they satisfied one of the following criteria: (i) Cscore ≥0.5 and protein coverage ≥50%; or (ii) supported by transcript evidence with coding sequences overlapping repeats by less than 20%. Gene models whose coding sequences overlapped repetitive regions by more than 20% were retained only if Cscore ≥0.9 and protein coverage ≥70%. Finally, all predicted proteins were searched against the Pfam database, and gene models with proteins containing more than 30% transposable element-related Pfam domains were removed. Additional manual curation eliminated incomplete gene models, poorly supported predictions lacking transcript evidence, and short single-exon genes (<300 bp coding sequence) without conserved protein domains or detectable expression.

